# Hidden immune memory niches in inflammatory skin diseases

**DOI:** 10.64898/2026.03.20.713219

**Authors:** Lloyd Steele, April R Foster, Kenny Roberts, Chloe Admane, Sebastian Birk, Pavel V Mazin, Amir Akbarnejad, Catherine Tudor, Benjamin Rumney, Hon Man Chan, Bayanne Olabi, Treasa Jiang, Victoria Rowe, David Baudry, Christine Hale, Elena Winheim, Elias Farr, Joe McWilliam, Laure Gambardella, Keerthi Priya Chakala, Nusayhah H Gopee, Aljes Binkevich, Alexander Predeus, Martin Prete, Amirhossein Vahidi, Edel A O’Toole, Daniela Basurto-Lozada, David Horsfall, Emily Stephenson, Vijaya Baskar Mahalingam Shanmugiah, Raif S Geha, Ramnik J. Xavier, Catherine Smith, Satveer Mahil, Mohammad Lotfollahi, Muzlifah Haniffa

## Abstract

Disease-associated histopathological features are widely used to identify tissue microenvironments or niches for diagnostics and treatment response in clinical practice. However, despite its widespread use, histopathology does not reveal the full cellular and molecular composition of known pathological niches. Furthermore, the existence of pathological niches that may not be histologically discernible remains unknown. In this study, we generated a spatially-resolved multi-modal molecular atlas of ∼5 million human skin cells (including 113 skin sections profiled using Xenium-5k) and applied deep learning to unbiasedly decode 26 skin niches in health and disease. Several disease-associated niches corresponded to known histopathological features, and we defined their cellular and molecular features, co-localisations, and interactions. Additionally, we discovered an immunologically active role for skin appendageal structures in disease mechanisms, potentially contributing to inflammatory memory, that was not identifiable using standard histopathological analysis. These include a resident memory T cell-rich niche in the sebaceous gland and a plasma cell-rich niche in the sweat gland, analogous to the gland-associated immune niche in lung. Finally, we illustrate how our atlas can be used to generate high-resolution representations using transfer learning, resolving rare T cell and sebocyte subsets not possible in the original studies, validating niche identification, and the spatial enrichment of candidate genes linked to disease-associated genetic variants. Overall, our study links histopathology and atlas-scale genomics to reveal novel insights into inflammatory disease pathogenesis, chronicity, and potentially curative therapeutic avenues, using skin as an exemplar tissue for this approach.

## INTRODUCTION

Our understanding of human tissues in health and disease has been recently accelerated by high-throughput single cell multiomics technologies^1^. We now have the cellular catalogue of most tissues^2–5^, but how these cells come together to form tissue ecosystems or niches that perform diverse structural and functional roles is poorly understood. Haematoxylin and eosin (H&E)-based histopathology is widely used to identify tissue niches for diagnostics, but does not capture the cellular and molecular features of known niches in health and disease. Furthermore, clinically relevant processes are not always identifiable from standard H&E analysis^1^. For instance, inflammatory memory, which causes the relapse of inflammatory disease following treatment cessation, is predicted to exist in histologically normal tissue^6–8^. If identified and understood, targeting memory-related niches promises curative therapeutic strategies for many patients and significant healthcare savings.

Spatial transcriptomics technologies can now profile thousands of genes at single-cell resolution *in situ*. Linking these high-plexity *in situ* tissue profiles with H&E-based clinical histopathology analysis presents a unique opportunity to dissect known histopathological structures in unprecedented detail and potentially identify novel hidden niches. In particular, the identification of novel disease-associated niches that are not evident from H&E would generate new perspectives into disease and help understand latent processes such as inflammatory memory. Furthermore, while recent studies have linked disease susceptibility genes to manually annotated disease-associated histopathological features^9–11^, it is unknown whether such genes are enriched within disease-associated niches not evident from H&E.

To date, high-plexity single-cell resolution spatial atlases harmonised with single-cell RNA-sequencing (scRNA-seq) data have not been assembled for human tissue in health and disease at scale, and thus tissue niches have not been systematically decoded. Prior *in situ* maps of human tissue have predominantly been low-plexity, precluding a deep understanding of tissue niches^12–15^. The human skin is a tissue that is highly accessible and for which histopathology is routinely performed for clinical diagnosis, including inflammatory disorders. Atopic dermatitis (AD) and psoriasis are two highly prevalent inflammatory skin diseases, affecting >300 million individuals globally^16,17^, that: exhibit polarised inflammatory states (Th2 vs Th17/Th1); respond to distinct targeted therapies (IL-4/IL-13 inhibition vs IL-17/IL-23 inhibition); and are characterized by clinical recurrence in the same anatomical site, supporting the notion of tissue-residing inflammatory memory^7,18,19^. Studying human skin in health and AD/psoriasis thus provides the opportunity to understand how immune responses are enacted and potentially stored *in vivo* with spatial resolution.

Here we link high-plexity single cell resolution spatial genomics, H&E-based tissue imaging, and genetic susceptibility variants, utilising generative artificial intelligence (AI) approaches to identify and understand tissue niches in health and disease across more than 160 tissue sections. We use human skin as an exemplar tissue, with a framework that presents a new way to investigate human biology in health and disease through establishing a harmonised *in situ* molecular atlas. We use cutting-edge deep learning methods to construct a comprehensive spatially-resolved single cell atlas of human skin to unbiasedly define histopathologically-known and hidden niches in healthy and diseased skin^20,21^. We reveal 26 niches, including niches associated with known histopathological features, such as a *Tzone-like* niche corresponding to perivascular inflammation, and two “novel” disease niches associated with inflammatory memory that are not evident from histopathology: i) a sebaceous gland-associated niche enriched in CD8^+^ resident memory T (T_RM_) cells and ii) an eccrine sweat gland-associated niche enriched in plasma cells that likely reflects a human cross-tissue immune niche. These hidden niches are enriched in candidate genes associated with disease risk. Within AD epidermal niches, we also identify type 2 T_RM_ and polarised group 3 innate lymphoid cells (*ILC3_CCL1+PTGDS+)*, which we validate in a range of type 2 inflammatory skin diseases, providing new therapeutic targets.

Our work represents a unique AI-served multi-modal resource that facilitates mapping new single-cell and spatial data to compare tissue niches and cell types across diseases, which we envisage to be a standard workflow for interpreting future datasets. To illustrate this, we show how transfer learning can resolve fine cell types and niches in unseen datasets, and how spatially-informed annotations can inform the interpretation of disease-associated genetic variants. Our datasets, AI models, and systematic workflow are publicly available for download and interactive exploration through the following user-friendly webportal [to add]^22^, creating a foundational representation of human skin.

## RESULTS

### Constructing a spatially-resolved skin cell atlas

To generate a spatially-informed skin cell atlas in health and inflammatory disease, we first assembled a scRNA-seq atlas, followed by joint integration with Xenium-5k data (Fig. 1a and Extended Data Fig. 1a).

**Figure 1.**
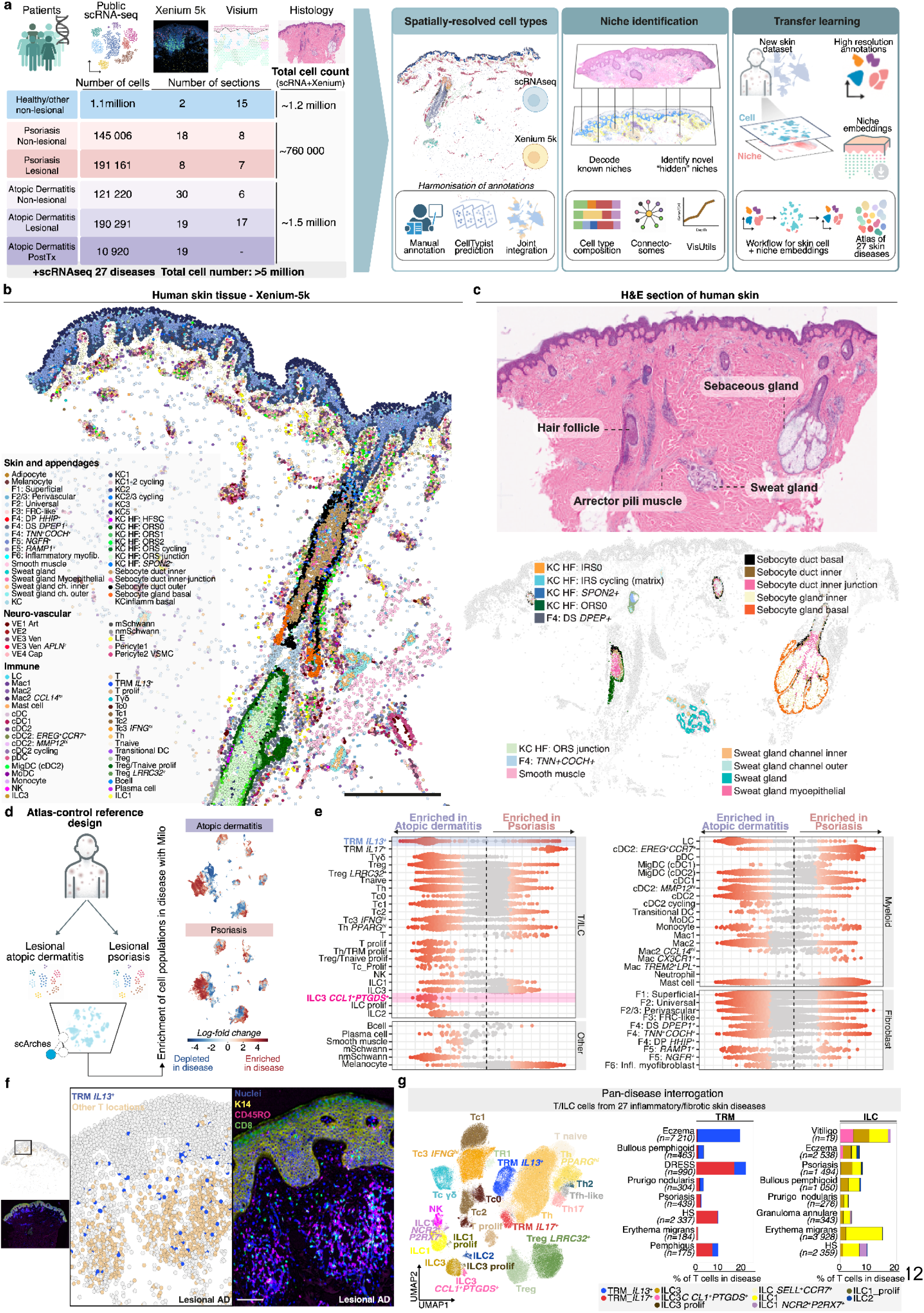
Constructing a spatially-informed atlas of human skin. **a,** Overview of study methodology, including skin atlas integration, niche identification and analysis, definition of zonation patterns in human skin, and a resource for transfer learning including an atlas of 27 skin diseases. Tx: treatment. **b,** Human adult skin from Xenium-5k data with each cell coloured by cell type. Scale bar = 500 µm. **c,** Haematoxylin and eosin-stained tissue (top) and the same tissue profiled by Xenium 5k, with structural cells from the pilosebaceous unit and sweat gland shown (bottom). HF: hair follicle. IRS: internal root sheath. ORS: outer root sheath. **d,** The ACR design first trains an scVI model using samples from a reference atlas (non-lesional), followed by scArches fine-tuning with a query dataset (here, lesional psoriasis and AD). Milo is then used to perform differential abundance testing. **e,** Differential abundance results from Milo for lesional AD skin vs lesional psoriasis skin. Coloured dots represent significant differences (false discovery rate 0.1). **f,** Lesional human skin profiled by Xenium 5k with T_RM_ cells coloured. AkoyaPhenocyler profiling of AD skin in lesional (matching Xenium) skin, showing CD8^+^ T cells in epidermis. Scale bar = 100 µm. **g,** UMAP visualisation of skin T cell atlas. Bar plot showing diseases with highest cell counts for *T_RM__IL13+* and *ILC3+_CCL1+PTGDS+* cells. AD: atopic dermatitis.

For scRNA-seq data, we curated and integrated 338 skin samples derived from 37 published studies, including healthy, non-lesional (macroscopically normal/unaffected skin in individuals with disease elsewhere), and lesional AD and psoriasis samples (Supplementary Table 1). To uniformly process the data, we remapped raw sequencing data and applied the same pre-processing pipeline (Methods). For the ∼1.7 million high-quality scRNA transcriptomes identified, we performed integration using single-cell variational inference (scVI)^23^.

We defined 107 cell types at the highest granularity of annotation in scRNA-seq data (Supplementary Fig. 1a and Methods). As the integration provided higher cell numbers and multiple anatomical sites in comparison to single studies, our analysis identified for the first time transcriptome-wide profiles for rare cell types, such as *ASIC2+_mechanoreceptors* (*ASIC2^+^*, *DCC^+^*)^24^ in healthy and non-lesional steady-state adult skin, as well as cell types not previously resolved in inflamed AD and/or psoriasis skin, including eosinophils (*CLC^+^*, *CCR3^+^, IL4^+^*) and perineural macrophages (*Mac_CX3CR1^+^; P2RY12^+^*) (Extended Data Fig. 2a).

To spatially resolve cell types and define tissue niches, we generated single-cell resolution spatial transcriptomics data from 113 skin sections using a 5000-plex gene panel (10x Genomics Xenium). We included healthy, non-lesional, and lesional skin from AD and psoriasis, split into core and validation datasets, comprising ∼1.7 million cells (Extended Data Fig. 1a). To investigate spatial niches during severe active disease and remission, we included matched pre-treatment and week 12 post-treatment (dupilumab (IL4/IL13 blockade) or methotrexate (broad immunosuppressant)) samples from patients with AD who achieved a good treatment response (clear/nearly clear skin; Methods). To orthogonally validate cell type locations and investigate zonation patterns (gene expression by tissue depth) in human skin, we generated spot-level transcriptome-wide data from 53 sections (10x Genomics Visium) (patient demographics in Supplementary Table 2-3).

Using high-plexity single-cell resolution molecular profiles collected at scale, we were able to harmonise annotations between spatial transcriptomic and scRNA-seq data in the same latent space, providing for the first time a spatially-resolved, integrated atlas of human skin (Fig. 1b). To ensure robust harmonised annotations, we tested three approaches: *de novo* annotation of spatial transcriptomic data using marker genes derived from scRNA-seq, a label transfer approach using a logistic regression classifier trained on our scRNA-seq reference^25^, and joint integration of spatial transcriptomic and scRNA-seq data using scANVI^26^ (Fig. 1a and Methods). Joint integration generated the best cellular representations, being the only method that resolved rare cell types, including perineural macrophages (*Mac_CX3CR1^+^*), *Merkel cells*, and ILC subsets (Extended Data Fig. 2b)^27^. Overall, we identified 99 out of the 107 cell types from scRNA-seq in the spatial transcriptomic data. We comprehensively categorised cell type populations across both modalities using the same marker gene sets (Supplementary Fig. 2-11). The remaining eight cell types were likely not sampled due to tissue depth (*Skeletal muscle, Cartilage, Satellite cell)*, anatomical region of skin analysed (*ASIC2^+^_Mechanoreceptor),* and transcriptomic similarity to other immune cells (*Th_PPARGhi, type 1 regulatory T cell (TR1), Tnaive_CD8, Eosinophil*) (discussed in Supplementary Note 1).

We next manually confirmed that the location of spatially-resolved skin cell types was in agreement with their predicted location from histology using two well-defined microanatomical structures: the pilosebaceous unit and sweat glands. This identified the expected location of known populations, such as the internal root sheath (*KC_HF: IRS0/1*) and dermal papilla fibroblasts (*F4: DP_HHIP^+^*) (Extended Data Fig. 3a). Our spatial information also revealed fine detail within *in situ* microanatomical structures for the first time, including inner, junctional, and outer sebaceous duct cells, as well as inner and outer eccrine sweat gland channels (Fig. 1c). From the differentially expressed genes for each population (Methods), we observed anatomical specialisation for distinct biochemical processes (Supplementary Fig. 8c), such as the activation and inactivation of androgens occurring as an interplay between the sebocyte subsets^28^ (Extended Data Fig. 3b), further supporting their distinct properties.

Next, we used our integrated atlas to investigate how active inflammation alters the skin’s cellular composition in AD and psoriasis. We utilised an Atlas-Control-Reference design^29^ to identify differentially abundant cell populations using scRNA-seq data, which first trains a model using a reference (atlas) dataset followed by fine-tuning using a query dataset via scArches (Fig. 1d)^30^. This design has been shown to provide a higher sensitivity (minimises loss of biological variation) to identify disease-associated cells compared to standard integration^29^. Using this approach, we compared lesional AD and lesional psoriasis after reference (non-lesional) atlas training. We confirmed the expected features of these two inflammatory diseases and identified new features in inflamed AD skin (Fig. 1e and Supplementary Fig. 12a). As expected, the *ILC2* (*PTGDR2^+^*, *IL1RL1^+^*, *IL13^+^, GATA3^+^*) population was enriched in AD compared to psoriasis. Additionally, we identified that a novel *ILC3_CCL1^+^PTGDS^+^* population (*KLRF2^+^*^31^*, CSF2^+^*, *TNFSF4^+^*, *SPINK2^+^, ID3^+^, CCL1^+^*), and a similar proliferative population (*ILC_Prolif*; *KLRF2^+^*, *CSF2^+^*, *MKI67^+^*)), were enriched in AD (Fig. 1e and Supplementary Fig. 3c,d). We used Xenium data and complementary spatial transcriptomic data (10x Visium) to confirm that *ILC3_CCL1^+^PTGDS^+^*cells were enriched in AD and additionally reveal their location in superficial skin (Extended Data Fig. 2c-d). Supporting this predicted location, *ILC3_CCL1^+^PTGDS^+^* cells showed the highest expression of *CCL1* (Extended Data Fig. 2e), and *CCL1* expression was specifically increased in AD in superficial skin (Extended Data Fig. 2d). These results suggest that the *ILC3_CCL1^+^PTGDS^+^*cell state represents a novel polarisation of ILCs in human skin in AD, identified across modalities.

Epidermal CD8*^+^* T_RM_s producing IL17 and IFNγ (termed T_RM_17 and T_RM_1 cells) are recognised in skin in psoriasis and vitiligo^32–35^, and we observed that T_RM_17 cells (expressing *IL17A*, *IL17F, IL23R,* and *CXCL13*) were enriched in psoriasis (Fig 1e). While polarised CD8^+^ T cells have been described in AD skin by ourselves and others^5,34,36–38^, their location is uncertain, and IL13-producing T_RM_s (T_RM_2) are not recognised^32,39^. We recently reported that the microenvironment-induced gene expression of T cells located in the epidermis resembled T_RM_ cells in AD^21^, consistent with spatial imprinting of the T_RM_ phenotype^40^. Here, we identified IL13-producing T_RM_ cells (expressing *IL13*, *CCL1,* and *IL9R*) as a distinct cell type across multiple AD tissue sections, characterised by epidermal T_RM_-associated genes (*CD8A^+^*, *ITGAE^+^, ZNF683^+^, CCR8^+^*)^41,42^. Consistently, we located these *T_RM__IL13^+^* cells predominantly in the epidermis (Fig. 1f), and we validated their presence using multiplex protein staining in lesional and post-treatment AD skin (Fig. 1f and Extended Data Fig. 2f). Furthermore, we were able to utilise our high-resolution annotations to identify co-localisation with *ILC3_CCL1^+^PTGDS^+^* cells (Extended Data Fig. 2g and Supplementary Fig. 12b). A co-localisation with *ILC3_CCL1^+^PTGDS^+^*cells was further supported by a CCL1/CCR8 interaction inferred from cell-cell communication analysis (Extended Data Fig. 2h and Methods). Targeting the T_RM_ IL23 receptor has proven highly efficacious in psoriasis treatment^43^, with IL23 required for the proliferation and retention of skin-resident *T_RM__IL17^+^*cells^44^, and our results suggest that *T_RM__IL13^+^*and *ILC3_CCL1^+^PTGDS^+^* are novel therapeutic target populations in AD.

Finally, we sought to validate these novel *T_RM__IL13^+^*and *ILC3_CCL1^+^PTGDS^+^* in different disease datasets. We mapped 27 inflammatory and fibrotic skin diseases to our existing skin scRNA-seq atlas in a semi-supervised manner, generating ∼3.6 million scRNA-seq cells overall (totalling >5 million cells including Xenium data), including ∼400,000 T cells (Supplementary Fig. 13a and Methods). We identified *T_RM__IL13^+^*and *ILC3_CCL1^+^PTGDS^+^* cells in other skin diseases associated with type 2 inflammation, including bullous pemphigoid (Fig. 1g and Supplementary Data Fig. 13b-c). In addition to validating *T_RM__IL13^+^* and *ILC3_CCL1^+^PTGDS^+^* cells across different type 2 inflammation-associated diseases, we detected cell states not present in our original atlas through manual annotation of newly mapped cells. Among newly identified cells were T follicular helper-like (Tfh-like) cells (*IL21^+^*, *CXCR5^+^*, and *CXCL13^+^*), which were identified in hidradenitis suppurativa, in which tertiary lymphoid structures are reported, as well as other diseases such as lichen planus (Supplementary Data Fig. 13d-e).

Overall, we demonstrate the value of our spatially-resolved atlas to identify and spatially localise novel rare pathogenic cell states (*ILC3_CCL1+PTGDS+* cells represent 0.04% of 1.7 million scRNA-seq cells), revealing potential colocalisation and interactions of disease-associated *ILC3_CCL1+PTGDS+* with *T_RM__IL13^+^* cells in type 2 inflammation.

### Defining human skin niches using deep learning

The function of a tissue is more than the sum of its constituent cells, as organized assemblies of different cell types enact distinct functions^45^. We next used a deep learning-based method (NicheCompass^20^) to provide an unsupervised decomposition of human skin into distinct niches. NicheCompass considers both a cell’s gene expression and its surrounding neighbourhood, and applies a graph neural network to these neighbourhoods to derive niche embeddings, thus identifying spatially-recurring cellular neighbourhoods across tissue sections (Extended Data Fig. 4a,b). We then compared Xenium profiles to tissue H&E and utilised our high-resolution cell annotations to explore the composition of each niche (Fig. 2a-c and Extended Data Fig. 4c-d).

**Figure 2.**
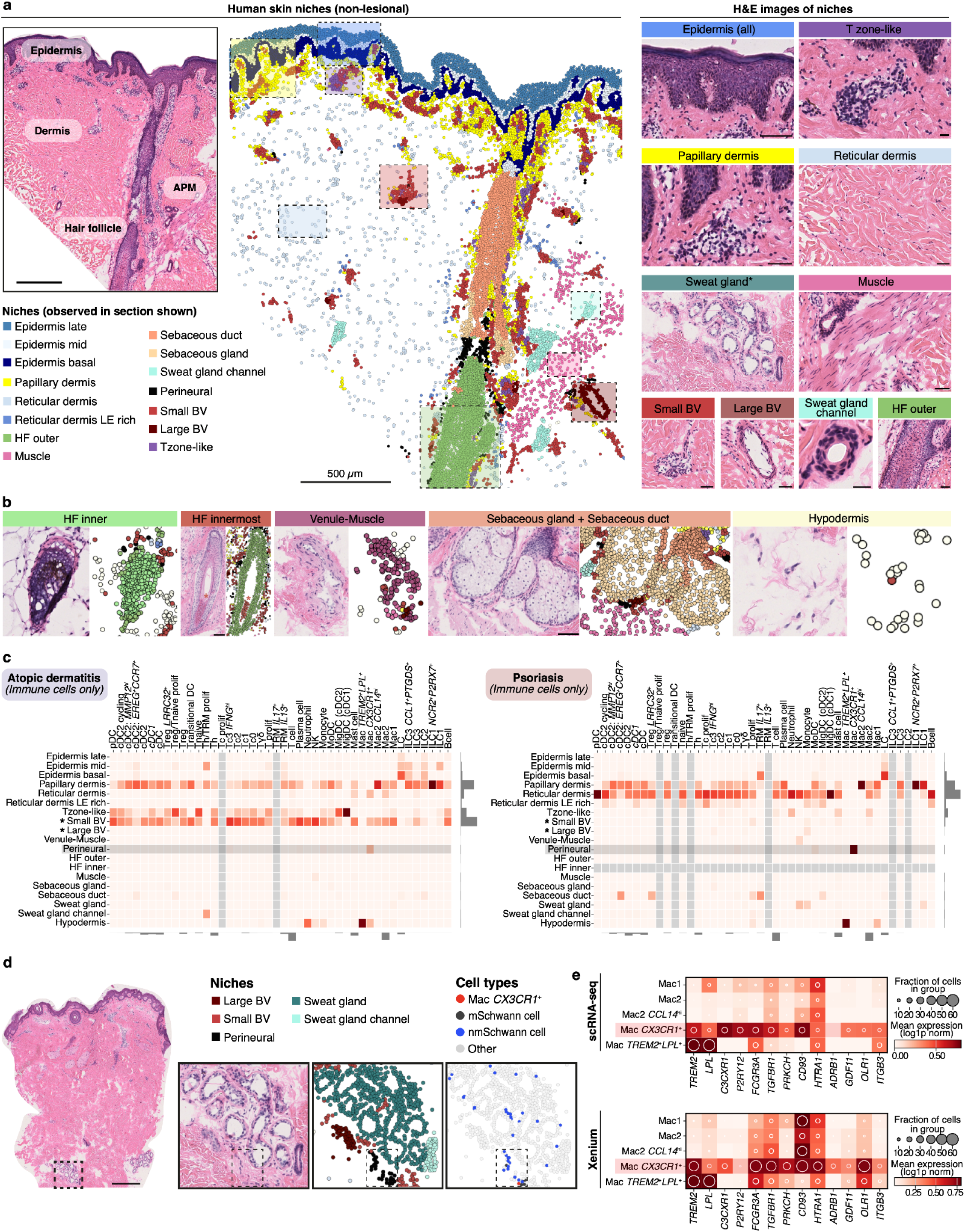
Defining human skin niches using deep learning. **a,** H&E-stained skin tissue (left), and same tissue profiled by Xenium 5k with cells coloured by niche cluster (centre), where coloured boxes indicate regions shown in the right panel. Asterisk indicates a niche not observed in sample shown. APM: Arrector pili muscle, Scale bar = 500µm. **b,** H&E-stained skin tissue and Xenium 5k with cells coloured by niche for select niches not evident in panel a. The HF_innermost niche was not found with >10 cells in non-lesional skin, but this likely reflects frequency of the niche as it contained rare hair follicle cell types found in healthy states from scRNA-seq data. Full tissue H&Es are shown in Extended Fig. 4d **c,** Heatmap showing the niche distribution of immune cells in AD and psoriasis non-lesional skin (niches with minimum of 10 cells), normalized by column. Bars indicate cell (bottom) and niche (right) abundance. The *Small_BV* niche contained abundant immune cells, but immune cells were rare in the *Large_BV* niche (marked with asterisks). *Mac_CX3CR1+* cells (consistent with a perineural macrophage phenotype) were observed in the perineural niche. **d,** H&E-stained skin tissue showing sweat gland region (dashed box). For this region, we show zoomed H&E (left), corresponding niche annotations (middle), and location of Schwann cells and perineural macrophages (*Mac_CX3CR1+*) within this region (right), where dashed boxes mark the perineural niche region where *Mac_CX3CR1+* is located. Scale bar = 500µm. **e,** Gene expression for *Mac_CX3CR1+* cells by disease in scRNA-seq and Xenium 5k data. AD: atopic dermatitis. BV: blood vessel. H&E: Haematoxylin and eosin, HF: Hair Follicle. scRNA-seq: single-cell RNA sequencing.

We first constructed an atlas of non-inflamed (healthy and non-lesional) niches (50 sections), which we later leveraged to map and understand niches in lesional/inflamed and post-treatment skin. In non-inflamed skin, we identified 20 distinct niches found in at least 3 tissue sections (Extended Data Fig. 4a,e). These deep learning-identified niches largely correlated with known histological skin structures (Fig. 2a-b). Six niches formed sequential layers in the epidermis (*Epidermis_basal*, *Epidermis_mid*, *Epidermis_late*) and dermis/hypodermis *(Papillary_dermis, Reticular_dermis, Hypodermis*). The other 14 niche categories were more focal and included immune (*T zone-like*), sweat gland (*Sweat_gland*, *Sweat_gland_channel*), pilosebaceous unit (*HF_inner*, *HF_outer*, *Muscle*, *Sebaceous_gland, Sebaceous_duct*), neurovascular (*Large_blood_vessel, Small_blood_vessel*, *Perineural*), and other (*Reticular_dermis_LErich, VenuleMuscle, EpidermisInflamm_mid*) structures. The *EpidermisInflamm_mid* niche exhibited an increased proportion of Langerhans cells (*LC)* and *TRM_IL13^+^*cells compared to the *Epidermis_mid* niche (Extended Data Fig. 4f), and was predominantly present in two donors with AD with the most severe inflammation at the time of biopsy (Extended Data Fig. 4g). This finding is consistent with the presence of microscopic inflammation in macroscopically normal (non-lesional) skin in patients with severe inflammatory skin disease^37,46–48^.

Our high-resolution cell annotations allowed us to further explore the fine cell populations within niches not possible with selected protein markers (Fig. 2c and Supplementary Data Fig. 14a). Most immune cell populations in human skin, particularly T cell subsets, were restricted to three dermal niches: *Small_blood_vessel*; *Papillary_dermis*; and/or the perivascular *Tzone-like* niche (Fig. 2c). The deeper reticular dermis contained fewer immune cells, predominantly mast cells and macrophages. This enrichment of immune cells in the superficial dermis is consistent with previous findings based on select protein markers^49,50^. To validate distinct composition by tissue depth in different tissue sections and orthogonal technologies, we developed VisUtils: a tool to analyze spot-level spatial transcriptomic data that enables quantitative comparison of cell type abundance and gene expression by tissue depth (Fig. 1a and Methods). VisUtils utilises a manually annotated common tissue reference point, such as the dermo-epidemal junction in the skin, and then calculates cell type abundance and gene expression by tissue depth (Methods). VisUtils validated the distinct immune composition of superficial compared to deep dermis (Supplementary Data Fig. 15a).

We also identified distinct immune cell densities in the vascular niches. Whereas the *Small_blood_vessel niche* was immune-rich, the *Large_blood_vessel* niche was immune-sparse (Fig. 2c). This difference may be mediated by fibroblast and pericyte populations as the *Small_blood_vessel* niche contained fibroblasts (*F2/3: Perivascular* and *F3: Fibroblast reticular cell-like* (*FRC-like*)) and *Pericyte1* cells (Supplementary Fig. 14a), which both expressed genes involved in chemotaxis and the maintenance of immune cells (*CCL19^+^*, *CXCL12^+^*, *CH25H^+^*) (Extended Data Fig. 4h). In contrast, the *Large_blood_vessel* niche contained *Pericyte2_VSMC* (vascular smooth muscle cells) that expressed muscle-related genes (*CASQ2^+^*, *MYH11^+^*, *SCGA^+^*). These findings suggest distinct adaptation of stromal cells to support specialized functions within these niches, achieving fine control of T cell localisation within the skin.

We also revealed heterogeneous populations within other niches evident from H&E. For instance, the *Perineural* niche contained not only myelinated (*mSchwann*) and non-myelinated Schwann cells (*nmSchwann*), but also *F5: NGFR+* fibroblasts (Schwann-like or nerve-associated fibroblasts)^51^, *F2/3: Perivascular* fibroblasts, and the rare perineural macrophage *Mac_CX3CR1+* population (Fig. 2d,e and Supplementary Fig. 14a,b), which is posited to contribute to axon growth in mice^27,52^. *Mac_CX3CR1+* populations have recently been reported in human skin utilising our scRNA-seq atlas^5,52^, and here we illustrate how our spatial atlas can now resolve this rare cell type.

Overall, we decompose our *in situ* atlas of human skin into fine-grained niches utilising a biology-agnostic AI approach. Identified niches correspond to unique compositions of cell types and gene expression patterns in distinct microanatomical locations, serving as a baseline for investigating changes in disease.

### Inflammatory skin disease-related niches

We next interrogated tissue architectural changes in lesional AD and psoriasis skin. We derived niche embeddings for lesional and post-treatment skin using a reference-mapping approach with NicheCompass (Fig. 3a and Methods). We utilised histopathological annotations (Fig. 3b) and high-resolution cell type distributions (Extended Data Fig. 5a) to understand tissue niches, with validation of cell populations using VisUtils (Supplementary Fig. 16a). Overall, we identified 26 niches in lesional and post-treatment skin (Fig. 3a). We first focused on two niches that appeared expanded in lesional skin and corresponded to recognised focal histopathological abnormalities: *Tzone-like* and *Epidermal_Antigen-presenting cell^hi^*(*APC^hi^*) (Fig. 3a-b and Extended Data Fig. 6a,b).

**Figure 3.**
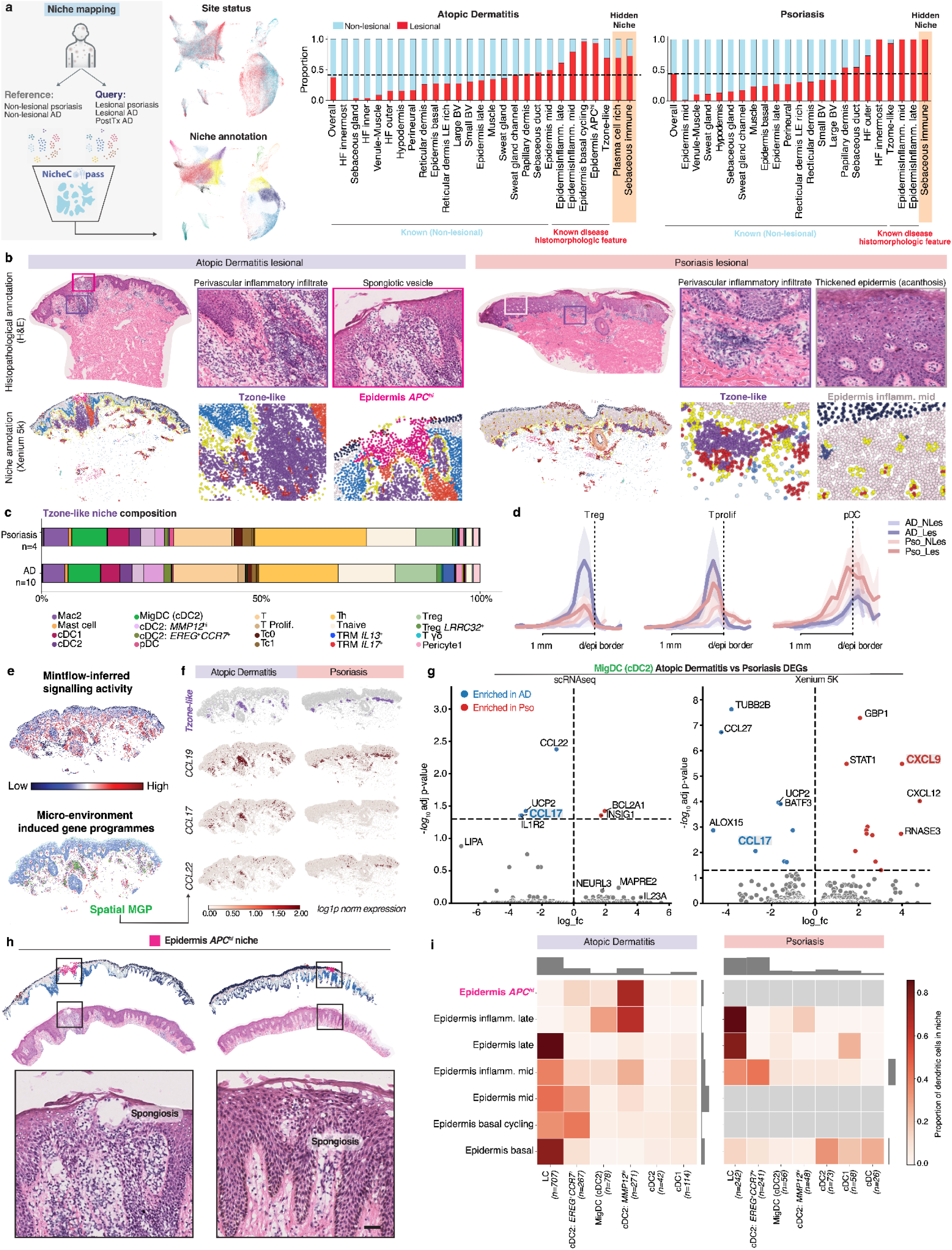
Inflammatory skin disease-related niches. **a,** Schematic of reference mapping at a niche level using NicheCompass, and UMAP visualization of niche embedding coloured by site status (lesional and non-lesional) and niche cluster name (left). Bar plot of niche composition by skin status (lesional vs non-lesional baseline), with the leftmost bar indicating the overall proportion of cells for each status. With increased cell numbers, we identified small numbers of the *Sebaceous_immune* cellniches in non-lesional skin. **b,** H&E stained tissue of inflamed AD and psoriasis with histopathological annotations (top row) and Xenium 5k of same tissue coloured by niche (bottom row). **c,** Composition of *Tzone-like* niche by cell type in inflamed AD and psoriasis (baseline lesional). **d,** VisUtils plots for *Treg*, *T_prolif*, and *pDC* abundance by disease status in Visium data. Mean +- 2 standard deviations of mean. **e,** Mintflow-inferred signalling activity (top, see Methods), and microenvironment-induced (spatial) gene programme detected by Mintflow (bottom). A microenvironment-induced (spatial) gene programme (green) corresponded to the *Tzone-like* niche. **f,** Inflamed AD and psoriasis sections highlighting the *Tzone-like* niche (top) and with all cells coloured by expression of the chemokines *CCL19*, *CCL17*, and *CCL22* (chemokines from the microenvironment-induced gene programme). **g,** Volcano plots from differential gene expression testing for *MigDC (cDC2)* cells for lesional AD vs lesional psoriasis skin in scRNA-seq and Xenium data. **h,** H+E and epidermis coloured by niche (Xenium 5k) for AD lesional samples (left), showing spongiosis/spongiotic vesicle. **i,** Heatmap showing distribution of epidermal dendritic cells (row) by niche in AD and psoriasis. Grey boxes indicate absence of niche (<10 cells). AD, atopic dermatitis. H&E: Haematoxylin and eosin.

The *Tzone-like* niche was found in almost all lesional tissue sections (Extended Data Fig. 6a) and corresponded to histopathological perivascular infiltrate (Fig. 3b and Extended Data Fig. 6b), which represents the site of extravasation of immune cells from the blood to the skin (Supplementary Data Fig. 17a). While perivascular infiltrate is known to be expanded in inflammatory states in both AD and psoriasis skin, its H&E appearance is identical for these two diseases (Fig. 3b). We therefore utilised our single-cell resolution spatial atlas to identify the shared and distinct features of this niche by disease.

The overall composition of the *Tzone-like* niche was notably similar across 13 sections in both AD and psoriasis. Shared features of the *Tzone-like* niche included the presence of *F3: Fibroblast reticular cell (FRC)-like* fibroblasts, *Tnaive*, *T helper* (Th) cells, and *MigDC (cDC2s)* (Fig. 3c), consistent with a lymphoid-like tissue region as previously reported^51^.

Cellular differences could also be noted in the *Tzone-like* niche. A higher proportion of *Tregs, T_Prolif*, *cDC2: MMP12hi*, and *Pericyte1* cells was observed in AD. In contrast, a higher proportion of *pDC* and cytotoxic T (*Tc2* and *Tc3_IFNGhi*) cells was observed in psoriasis. We validated these populations through consensus with VisUtils (Fig. 3d and Extended Data Fig. 6c) and through assessment in a further 11 lesional Xenium sections (Extended Data Fig. 6d).

Next, we investigated how distinct signalling within the *Tzone-like* niche in each disease may mediate these distinct cellular compositions. We dissected microenvironment-induced gene programs (MGPs) in the *Tzone-like* niche of AD and psoriasis using MintFlow - a generative AI framework that we developed in orthogonal work^21^. For AD, the MGP in the cells of the *Tzone-like* niche included the chemokines *CCL19*, *CCL17*, and *CCL22* (Fig. 3e and Extended Data Fig. 6e). We queried these chemokines in psoriasis to identify shared and disease-specific features. *CCL19,* a chemoattractant for CCR7^+^ *T_naive* and dendritic cells (DCs) (including *MigDC* subtypes and *LCs*; Supplementary Fig. 10c)^51,57^, was similarly expressed in both AD and psoriasis, consistent with the overall similar composition of the *Tzone-like* niche across AD and psoriasis. *CCL17* appeared more specific to AD (Fig. 3f and Extended Data Fig. 6f,g), consistent with its role as an available clinical biomarker for AD^58^. CCL17 and CCL22 have been reported to attract *Tregs* to DCs^59–62^, and thus distinct *CCL17* expression may potentially contribute to the increased prevalence of *Tregs* in AD (Fig. 1f and 3c). CCL17 has also been reported to be essential for the migration of CCR7+ cutaneous *DCs* to CCL19^hi^ regions^63^, which was the *Tzone-like* niche in skin (Fig. 3f), which may partially explain differential *cDC2: MMP12hi* abundance in this niche.

Studies have correlated an upregulation of *CCL17* and *CCL22* with AD and asthma severity^64^, but here we could uniquely investigate expression across 107 cell types. We identified that *CCL17* was most highly expressed by *MigDC (cDC2)* cells (Extended Data Fig. 5h). We confirmed statistically significant increased CCL17 expression in *MigDC (cDC2)* cells in AD in both scRNA-seq and Xenium data (Fig. 3g and Methods), suggesting that distinct chemokine gradients within the histopathologically-identical *Tzone-like* niche contribute to a distinct cellular composition of this niche in AD and psoriasis.

Studies of epidermal myeloid populations in psoriasis have produced conflicting results, with reports of both enrichment and depletion of *LC*s^67^. Furthermore, in AD, the identity of the immune cell associated with the histopathological feature of spongiosis (epithelial intercellular oedema) is unclear as previous studies have used select protein markers, which likely miss fine differences between myeloid cell subpopulations^68^. We aimed to resolve this uncertainty by utilising our spatially-resolved, high-resolution cell types and niches to comprehensively define epidermal DC composition in AD and psoriasis (Fig. 3h,i and Extended Data Fig. 7a-b).

We observed that *cDC2: EREG^+^CCR7^+^* cells were relatively enriched in psoriasis compared to AD, whereas *LC*s were enriched in AD compared to psoriasis (Fig. 3i). We validated this finding in both scRNA-seq (Fig. 1d) and Visium data (Extended Data Fig. 7c). *cDC2: EREG^+^CCR7^+^*cells shared transcriptomic similarity to the *LC* population (Supplementary Fig. 10c) and thus these two populations may not be resolved as distinct populations in smaller studies or targeted protein studies, leading to previous reports of conflicting results for LC abundance^67^.

We identified that the disease-associated *Epidermal_APC^hi^*niche corresponded to focal spongiosis in five tissue sections (Fig. 3h and Extended Data Fig. 7a-b), highlighting the granularity of our niche capture and allowing us to define its DC composition. The *Epidermis_APC^hi^* niche was expectedly only observed in AD tissue sections (Extended Data Fig. 7b), including non-lesional skin of two patients with the most severe inflammation scores (Extended Data Fig. 7d). Interestingly, specifically within the *Epidermal_APC^hi^* niche, we identified different DC subset abundances compared to other epidermal niches. *LC*s were depleted in the *Epidermal_APC^hi^*, whereas *cDC2: MMP12hi* cells were highly enriched (Fig. 3i and Extended Data Fig. 7e). Our analysis thus identifies a focal myeloid adaptation (*cDC2: MMP12hi*) corresponding to epidermal spongiosis in AD, adding a novel cellular perspective to this well-defined histopathological feature.

Finally, we assessed broader epidermal differences between AD and psoriasis. *Epidermis_basal* and *Epidermis_basal_cycling* niches were increased in AD compared to psoriasis (Extended Data Fig. 7a), which were characterised by basal and cycling keratinocyte populations (Extended Data Fig. 5a). Consistent with this observation, an accumulation of basal and hyperproliferating epithelial cells has been linked to pathologic type 2 immune responses in other tissues^69^. Cycling keratinocyte populations were also increased in psoriasis (Supplementary Data Fig. 12a), but these were located within niches with more mature keratinocyte populations, namely *EpidermisInflamm_Mid* and *EpidermisInflamm_Late* (Extended Data Fig. 5a, Supplementary Fig. 7a).

In conclusion, through unbiased niche identification, we capture known disease-associated histomorphologic niches of inflammatory skin disease, such as perivascular infiltrate and spongiosis, allowing us to decompose their cellular composition at a high–resolution and resolve fine cellular differences between AD and psoriasis.

### Inflammatory myofibroblasts recruit distinct granulocytes in AD and psoriasis

We next illustrated the value of our dataset in understanding chemokine gradients for recruiting immune cells. Eosinophils are blood-derived and enriched in AD, but have not previously been captured by scRNA-seq in AD studies^5,36,70–73^. Using VisUtils, we identified that the eotaxin *CCL26*, which is the ligand for *CCR3^+^* eosinophils, was specifically increased in AD in the superficial dermis and epidermis (Extended Data Fig. 8a). Using cell-cell communication analysis, we identified that the major cellular sources of *CCL26* signalling were *F3: FRC-like* fibroblasts and *F6: Inflammatory myofibroblasts* (Extended Data Fig. 8b). This analysis also predicted that eosinophils signalled to fibroblasts via the type 2 cytokines *IL4* and *IL13* (Extended Data Fig. 8b), with eosinophils being uniquely high producers of *IL4* in human skin (Extended Data Fig. 8c).

Given reports that *F6: Inflammatory myofibroblasts* can recruit neutrophils in wounds and disease^51,74^, we hypothesised that *F6: Inflammatory myofibroblasts* may recruit eosinophils if exposed to type 2 cytokines. In AD skin, *F6: Inflammatory myofibroblasts* expressed high amounts of the eotaxin *CCL26*, whereas higher amounts of the neutrophil recruitment ligand *CXCL8* were observed in psoriasis (Extended Data Fig. 8d). We experimentally validated *F6: Inflammatory myofibroblasts* polarisation by stimulating primary human skin fibroblasts with a panel of cytokines *in vitro* (IL-13, IL-17, IL-1β, TNF) associated with AD and psoriasis (Extended Data Fig. 8e). IL-1β-stimulated fibroblasts generated the highest neutrophil-tropic activity (*CXCL8* expression), consistent with prior work in inflammatory bowel disease^74^ and further supporting the cross-tissue nature of this fibroblast state^51,75^. In contrast, IL-13-stimulated fibroblasts induced uniquely high levels of the eotaxin *CCL26*, supporting the differential ability of *F6: Inflammatory myofibroblasts* to recruit eosinophils or neutrophils in AD and psoriasis, respectively (Extended Data Fig. 8f,g).

### Inflammatory memory niches within skin sebaceous and sweat glands

AD and psoriasis cannot currently be cured, costing billions of pounds in direct treatment costs^76^. Disease persistence has been attributed to inflammatory memory, but this has been challenging to study, particularly in the absence of any distinguishable features seen in H&E-stained skin. To understand inflammatory memory, we sought to identify niches that were enriched in inflamed states in AD and psoriasis, did not correspond to known histopathological abnormalities, and persisted in week 12 post-dupilumab treated skin. Two skin niches matched these criteria - *Sebaceous_immune* and *Plasma_cell_rich* (Extended Data Fig. 9a) - raising the possibility of these niches harbouring or contributing to the formation of inflammatory memory. The *Sebaceous_immune* niche was observed in nine tissue sections in both AD and psoriasis, and the *Plasma_cell_rich* niche was observed in six tissue sections in AD only (Extended Data Fig. 6a and 9b).

The *Sebaceous_immune* niche was formed of immune cell populations within the sebaceous duct, including *T_RM_* cells, *Th* cells, *Tregs*, *MigDCs (cDC2)*, *cDC2* subsets, and *ILC* subsets (Fig. 4a and Extended Data Fig. 9c). Strikingly, the *Sebaceous_immune* niche contained the highest prevalence of T_RM_ cells across all skin niches (Fig. 4b,c). Given this high T_RM_ concentration, and previous work showing that the T_RM_ phenotype is spatially imprinted^40^, we hypothesised that the *Sebaceous_immune* niche fosters a unique epidermal microenvironment for supporting T_RM_ cells that contributes to inflammatory memory.

**Fig. 4.**
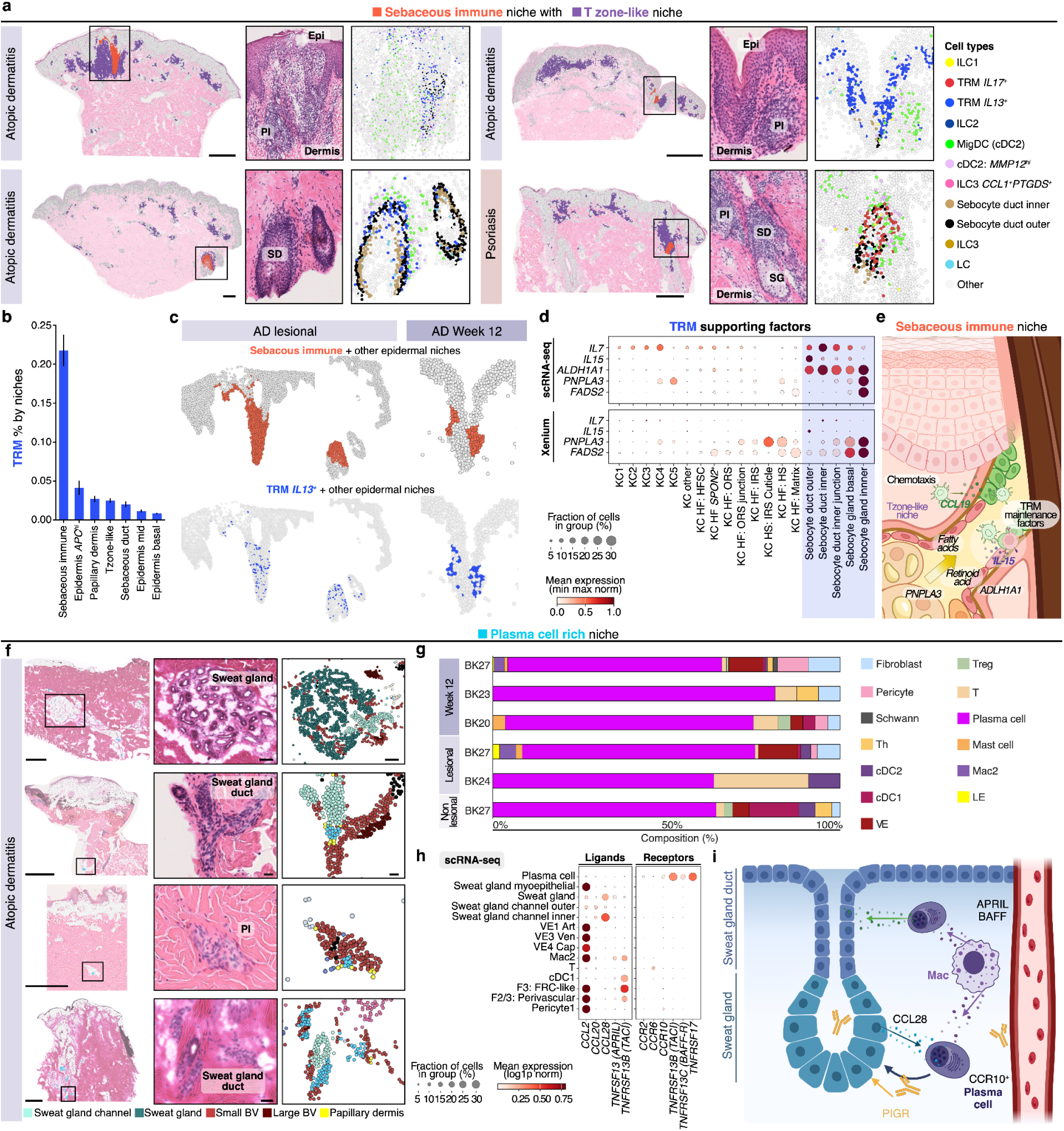
Hidden inflammatory skin niches within skin sebaceous and sweat glands. **a,** H&E stained human skin tissue with overlying Xenium 5k data indicating position of the *Sebaceous_immune* niche. Inlet panels: zoomed in regions of the H&E and the Xenium 5k data (coloured by select cell types). Scale bar 500µm. **b,** Prevalence of T_RM_ cells by niche (showing niches with highest prevalence). **c,** Skin tissue profiled by Xenium 5k showing epidermal and sebaceous niches, with *Sebaceous_immune* niche coloured (top row). Same tissue coloured by *T_RM__IL13+* cell type. **d,** Dot plots of expression of genes linked to generation of T_RM_ support factors in keratinocyte populations. **e,** Schematic of *Sebaceous_immune* niche. **f,** H&E stained human skin tissue and overlying Xenium 5k showing regions corresponding to the *Plasma_cell_rich* niche. Zoomed in regions show H&E of plasma cell niche region and Xenium tissue coloured by niche. Scale bar 500µm. **g,** Composition of *Plasma_cell_rich* niche by tissue section (minimum 10 cells) (simplified annotations, with highest level annotations in Supp. Fig. 18a). Cells with <10 cell counts excluded: **h,** Expression of receptors and ligands related to gland-associated immune niche in human lung tissue in cell types within the *Plasma_cell_rich* niche. **i,** Schematic of *Plasma_cell_rich* niche. H&E: haematoxylin and eosin. PI: perivascular infiltrate. SG: Sebaceous gland. SD: Sebaceous duct. TRM: resident memory T cell.

Well-described supporting factors for T_RM_ cells include cytokines (IL-7, IL-15)^777879^, retinoic acid^80,81^, and fatty acids^82^. We investigated expression of genes involved in the generation of these products in sebocyte populations relative to interfollicular epidermal populations (Methods). Compared to interfollicular epidermis, sebocytes expressed increased *IL7* and *IL15*; the retinoic-acid generating enzyme *ALDH1A1*^83,84^; and genes linked to fatty acid desaturation (*FADS2*) and lipid biosynthesis (*PNPLA3*)^85^ (Fig. 4d and Extended Data Fig. 9d-e). *IL15* has been shown to profoundly alter the program and function of T_RM_ cells^79^, and was specifically increased in *KC_Sebocyte_OuterDuct* cells in both scRNA-seq and Xenium data (Fig. 4d and Extended Data Fig. 9d-e), supporting this niche as a unique microenvironment for T_RM_ cells. PD-1 has also recently been reported as a requisite for skin T_RM_ cell formation^86^, but the cellular partners that express PD-L1 or PD-L2 during the various stages of the T_RM_ cell life cycle were uncertain^86^. We found that, within the *Sebaceous_immune* niche, *T_RM_* cells co-localised with *MigDCs (cDC2)* cells that express PD-L1 (encoded by *CD274*) (Extended Data Fig. 9c and Supplementary Fig. 10c), suggesting that they might perform such a partner role.

We observed that the *Sebaceous_immune* niche was located adjacent to the immune-dense *Tzone-like* niche across all patients (Fig. 4a and Extended Fig. 9f). We therefore considered that it represented a site of epidermotropism, where immune cells move from the vascularised dermis to the avascular epidermis. Supporting this role, lymphocyte and granulocyte chemotaxis processes were the highest enriched processes in the *Sebaceous_immune* niche (Extended Data Fig. 9g and Methods). Cell-cell communication analysis (Methods) also suggested active recruitment of immune cells, including *MigDC (cDC2)*, *cDC2: EREG+CCR7+*, and *T_naive* cells, to the *Sebaceous_immune* niche through *CCL19* expression by *KC_Sebocyte_DuctOuter* cells (Extended Data Fig. 9h).

Overall, we identify a novel hidden disease-associated immune niche enriched in AD and psoriasis that is not recognised as a histopathological feature. This niche contains the highest prevalence of T_RM_ cells across skin niches through provision of a unique epidermal microenvironment, distinct from interfollicular epidermis (Fig. 4e).

### Gland-associated plasma cell niche in human skin

A further niche not recognised as a histopathological feature was the *Plasma_cell_rich* niche (Fig. 4f), which was enriched in lesional and post-treatment samples in AD skin, but also present in a single non-lesional AD sample (Extended Data Fig. 9b). The *Plasma_cell_rich* niche contained predominantly *Plasma cells*, in addition to vascular/perivascular structures (*Pericyte1*, *VE3_Ven, F3: FRC-like, F2/3: Perivascular*) and small populations of other immune cells (*Mac2*, *T*, and *cDC1*) (Fig. 4g and Supplementary Fig. 18a). Plasma cells were also present in psoriasis, but they did not form a specific niche and were instead distributed across the *Tzone-like*, *Papillary_dermis*, and *Small_BV* niches (Extended Data Fig. 5a).

Plasma cells are a source of protective antibodies^88^ and have been identified in perivascular regions in mouse and human skin^89^. More recently, autonomous antibody production from plasma cells has been reported in a hair follicle niche in mouse skin^90^. Whether a similar hair follicle niche exists in humans is uncertain. Here, we identified that the *Plasma_cell_rich* niche was enriched near sweat glands/ducts and small blood vessels in human skin in AD (Fig. 4f), not the hair follicle. Notably, eccrine sweat glands are limited to the footpad in mice^91^, whereas in humans they are present across almost all anatomical regions, suggesting a species-specific difference in immune niches associated with glandular structures. A location of plasma cells near to sweat glands is also in keeping with previous observations of antibodies in human sweat and antibodies coating commensal organisms on the skin surface^92^. Further supportive of this finding of gland-associated plasma cells, we observed that a proportion of sweat gland cells expressed *PIGR* (Supplementary Data Fig. 18b), encoding polymeric immunoglobulin receptor (pIgR)^93–95^. pIgR enables transcytosis of immunoglobulin in lung and gut, facilitating the transport of antibodies to the mucosal surfaces.

A “gland-associated immune niche” (GAIN), in which plasma cells co-localise with pIgR^+^ glandular cells, is recognised in human lung^93^. Given the co-occurrence of plasma cells and eccrine glandular cells in skin, we next asked whether the same chemotactic and maintenance factors reported for lung GAIN - *CCL28*, *CCL20*, *CCL2, TNFSF13* (APRIL), and *TNFSF13B* (BAFF) - were present in human skin. We identified these same factors to be present in the *Plasma_cell_rich* niche in skin, including CCL28 (a chemokine for CCR10+ plasma cells) and *CCL20* (a chemokine for CCR10+ plasma cells^96^) from sweat gland and sweat gland channel/duct cells. We also identified B-cell maintenance factors (BAFF and APRIL) from macrophages and *F2/3: Perivascular* fibroblasts (Fig. 4h,i and Supplementary Data Fig. 18e). Similar chemoattractants and survival factors have also been reported for plasma cells in the gut^94,96–98^. Overall, our results suggest that the *Plasma_cell_rich* niche represents a common GAIN between skin and other human barrier tissues, and suggests increased complexity of AD pathogenesis beyond the paradigmatic view of a T-cell mediated disease.

### Spatially restricted expression of genetic risk loci genes within inflammatory memory niches

Inflammatory skin diseases are common and have well-established disease susceptibility loci^99,100^. Recent work has highlighted that the expression of disease-associated genetic variants is enriched within known histopathological niches, including lymphoid follicles in inflammatory bowel disease^10,1110^. However, whether genetic risk variants are highly expressed within hidden disease-associated niches is unknown as previous work has relied only upon manual histopathological annotations, not detecting hidden immune niches. We therefore selected disease-associated genes reported from recent genome-wide association studies (GWAS) in AD and psoriasis and scored these gene sets through gene module scoring (Methods) in cells across our defined skin niches^100–102^. We also used genes from the Open Targets platform, with a genetic association with AD and psoriasis, as a secondary gene set for validation (Methods)^103^.

We observed that candidate gene expression was expectedly enriched in known histopathological niches, namely *Tzone-like* (perivascular infiltrate) and *Epidermal_APC^hi^* (spongiosis). We also observed enrichment of candidate genes in the hidden niches (*Sebaceous_immune* and *Plasma_cell_rich*) (Fig. 5a and Extended Data Fig. 10a-c). This spatial correlation of disease-associated genes and hidden immune niches further supports their role as disease-associated niches not discernible from H&E. Across cell types, a higher expression of candidate genes was observed in cells frequently located within these pathogenic niches, including *MigDC (cDC2)* and/or *cDC2: EREG+CCR7+*, *Treg*/*Treg_LRRC32^+^*, *Tnaive*, *cDC2: MMP12hi*, and *T_RM__IL13^+^*/*T_RM__IL17^+^*cells (Extended Data Fig. 10d and Supplementary Fig. 19a).

**Figure 5.**
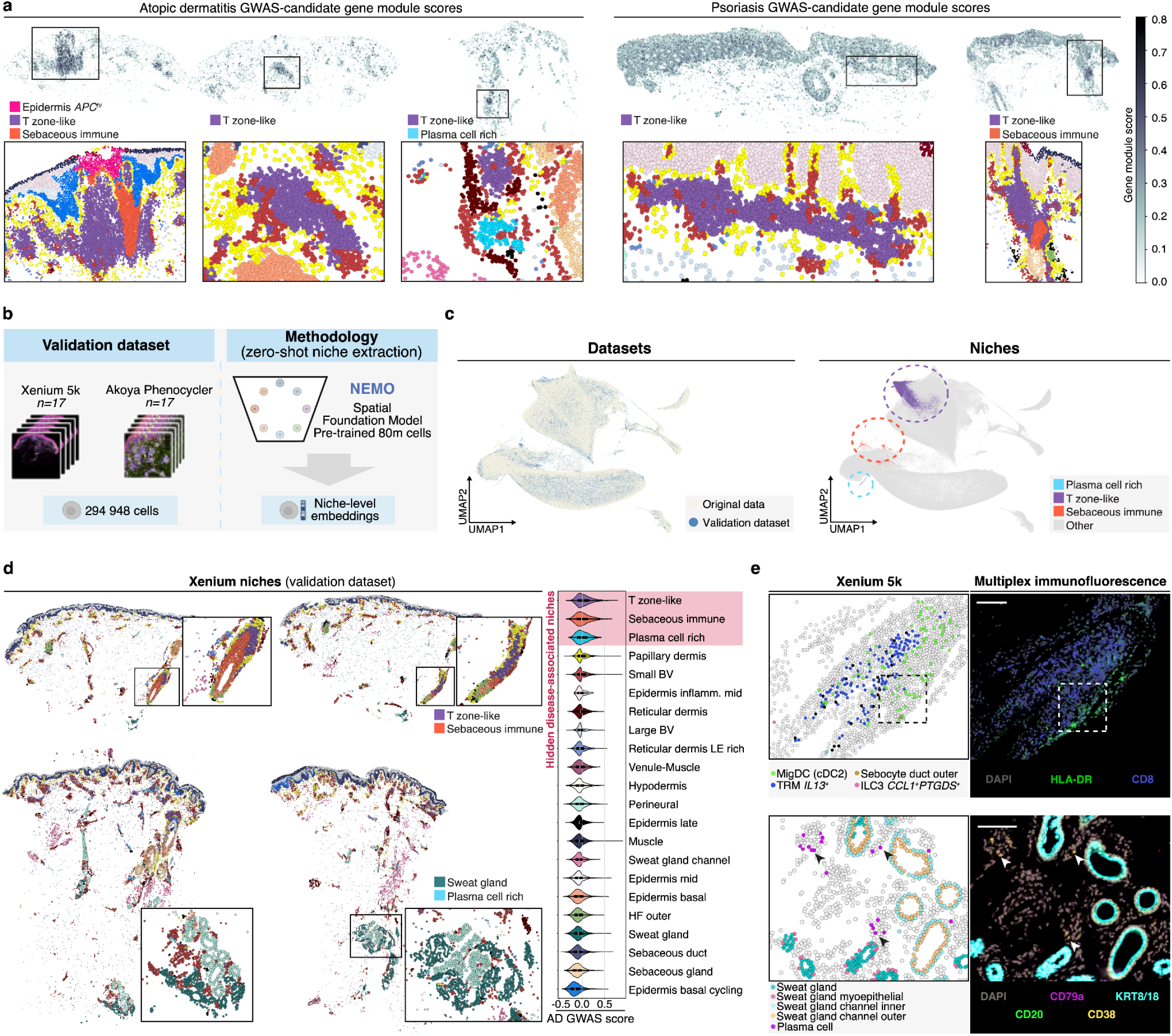
Spatially restricted expression of genetic risk loci genes within inflammatory memory niches. **a,** Spatial plots of gene module scores for each cell calculated using candidate genes for AD and psoriasis (Methods). Zoomed in regions show corresponding region to boxes, coloured by niche. **b,** Schematic of methodology used for validation data. **c,** UMAP visualisation for niche embeddings coloured by dataset (left) and niche label (right). **d,** Unseen validation dataset coloured by niche (left), and violin plots of gene module scores (calculated from candidate gene list) by tissue niche in validation dataset. **e,** Zoomed in regions of tissue shown in the panel d coloured by cell type (Xenium 5k) and AkoyaPhenocycler applied to the same tissue section (right). AD: atopic dermatitis.

Having established enrichment of genetic variants within our defined niches, we next sought to: 1) validate the existence of these disease-associated niches in an unseen dataset, 2) validate enrichment of genetic variants associated with these disease-associated niches, and 3) validate hidden niches at the protein level. To this end, we generated a new validation dataset that comprised 17 additional tissue sections generated from patients within our cohort, profiled by Xenium 5k (∼300 000 cells post-QC) and Akoya Phenocycler (multiple protein immunofluorescence) (Fig. 5b and Methods). To extract niche embeddings, we utilised a spatial transcriptomic foundation model developed in orthogonal work, pre-trained across 80 million single-cell resolution spatial transcriptomic human cells (Methods).

The new unseen validation dataset integrated well with our existing data (Methods), and both known (*Tzone-like*) and hidden (*Sebaceous_immune*, *Plasma_cell_rich*) disease-associated niches were recovered (Fig. 5c,d). While we did not identify the *Epidermal_APC^hi^* niche, overall this suggests robustness of our identified niches across both different datasets and niche identification methodologies. Known and hidden disease-associated niches again showed the highest gene module scores for genetic risk variants (Fig. 5d), consistent with our prior analysis. Using multiplex immunofluorescence on the same tissue sections, we were also able to confirm key cellular populations within these hidden immune niches at the protein level. Within the *Sebaceous_immune* niche, we identified a notably high CD8*^+^*T cell abundance (corresponding to *TRM_IL13+* cells from Xenium analysis) (Fig. 5e). We also validated the colocalisation of CD8*^+^*T cells with PD-L1*^+^ MigDCs* predicted from Xenium (Fig. 5e). Within the *Plasma_cell_rich* niche, we demonstrated plasma cells (CD79A*^+^*CD20^-^) adjacent to sweat glands and blood vessels within the skin (Fig. 5e).

Overall, we highlight the role of novel hidden disease-associated niches as sites associated with genetic risk variants for immune-mediated disease and we validate these niches in a novel dataset, across niche identification methodologies, and at the protein level, revealing new disease-associated structures beyond H&E.

### AI-Served skin resource for mapping new single-cell & spatial data

Generating millions of high-plexity single-cell data that enable high-resolution cell annotations and detailed insights into human disease can be prohibitively expensive, particularly for rarer skin diseases. Transfer learning is a widely used paradigm in machine learning, in which pre-training on large-scale data is performed prior to fine-tuning for a specific downstream task. We hypothesised that our harmonised, integrated atlas could be used to generate improved cellular and niche representations for future skin studies.

For cellular embeddings we provide two frameworks for utilizing our atlas: 1) scArches-skin, a resource-lite approach using fine-tuning of a pre-trained model (as described previously, Fig. 1d), and 2) integrative transfer, a resource-intensive approach using semi-supervised integration (Fig. 6a and Methods). The latter additionally can be used for different sized gene panels. To demonstrate our proposed methodologies, we initially used two scRNA-seq datasets of diseases not included in our original atlas (disease agnostic). First, we used inflamed alopecia areata skin data, motivated by the difficulty in cell type assignment of genetic risk variants due to the limited cell types resolved in the original study^104,105^. Secondly, we selected inflamed acne skin data because the original study did not detect sebocytes^106^, which were well characterised in our study in both spatial and scRNA-seq data and are associated with acne pathogenesis from histopathology.

**Figure 6.**
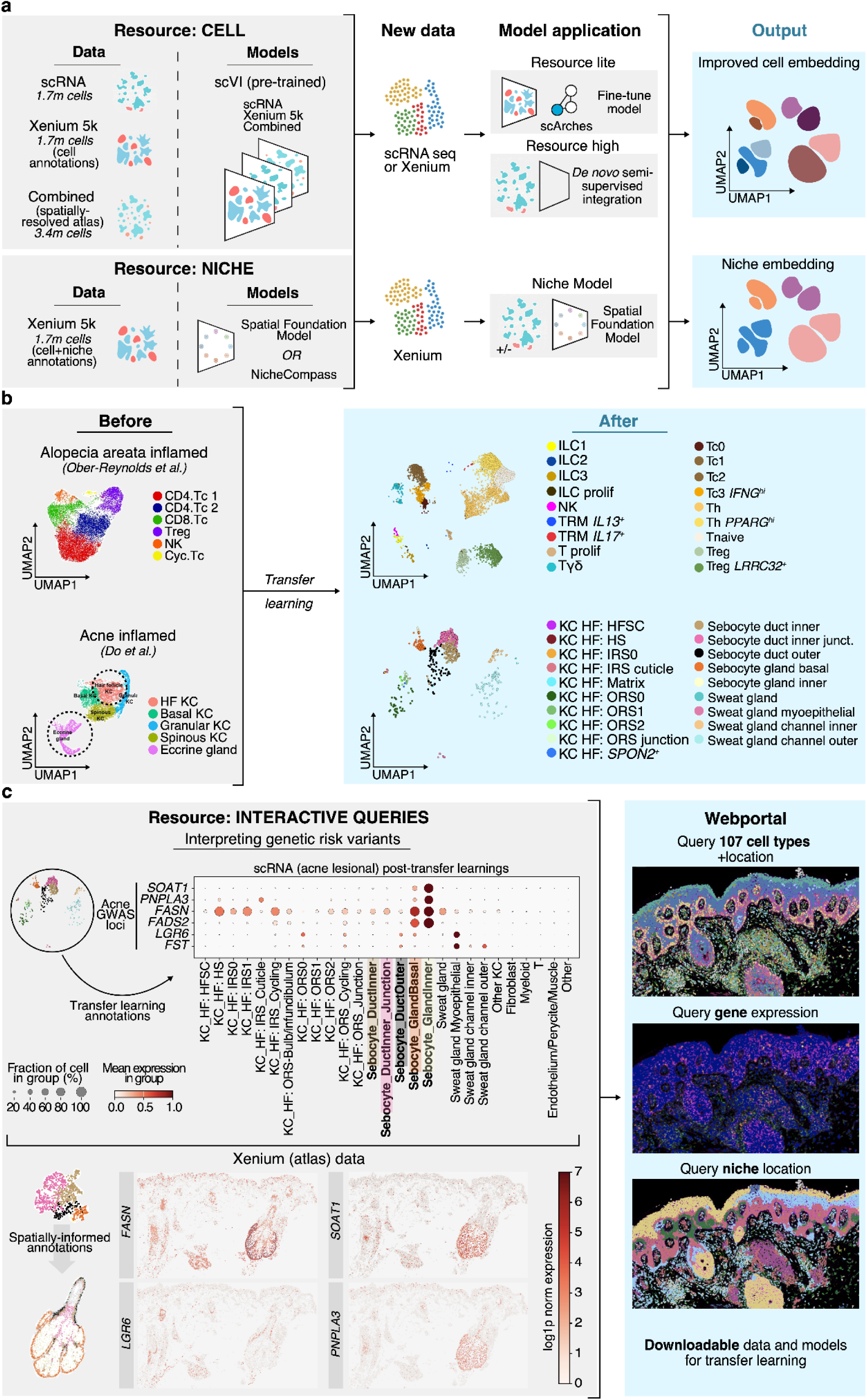
I-Served skin resource for mapping new single-cell & spatial data. **a,** Schematic showing resources available and strategies to generate cell and niche embeddings for novel/unseen skin data. **b,** UMAP visualisation for skin scRNA-seq data from original studies (left) and after transfer learning (right) for inflamed alopecia areata data (top row) and inflamed acne data (bottom row), showing improved granularity of annotations. **c,** Dot plot of candidate genes for acne genetic risk loci utilising improved annotations obtained for sebocyte cell populations in inflamed acne scRNA-seq. As these annotations were spatially-informed from our prior work, we could generate spatial plots of expression of select genes in Xenium-5k (atlas) data, showing how these genes can be spatially contextualised. Our webportal can be used to perform similar queries and to download data and models.

In the alopecia areata dataset, we were able to resolve rare immune cell types, including ILC and Treg subsets (Fig. 6b and Supplementary Data Fig. 20a), which the original study failed to identify. This included an ILC2 population of only 7 cells (<0.1% of T cells).

For acne, we resolved highly granular keratinocyte appendage cells that were not identified in the original study, including sebocyte subtypes (Fig. 6b and Supplementary Data Fig. 20c). We used these high-resolution annotations to further understand proposed genetic risk variants in acne^99,108^. Several candidate genes, not discussed in the original reports, were highly expressed in the *SebaceousGland_Inner* population (Fig. 6c and Supplementary Fig. 20d). As our annotations were spatially-informed, we could predict localisation of these genes to specific regions of the sebaceous gland, showing the value of our atlas to both: 1) resolve fine cell types that enables the cell type assignment of genetic risk variants, and 2) to understand the potential biological processes through which candidate genes from GWAS mediate disease risk from their spatial localisation (Fig. 6c).

For niche embeddings, we provide a pretrained NicheCompass model for reference-query mapping and a spatial foundation model for zero-shot niche embeddings (i.e. without model fine-tuning) (Fig. 6a) via our web portal (to be made available) to facilitate niche identification for the community. Our annotated niche data can also be used to contextualise future skin niche findings, as previously illustrated (Fig. 5b-d). In addition, our data is available as a queryable resource (Fig. 6c), which can be used for cell-specific investigation of drug targets or investigation (discussed in Supplementary Note 1 and Supplementary Fig. 21).

Overall, we demonstrate how our data and models can be used for knowledge transfer at a cell- and niche-level to maximise findings in other skin diseases using scRNA-seq data, providing a new paradigm for interpreting skin transcriptomic data.

## DISCUSSION

To identify and understand pathological molecular circuits, an improved understanding of tissue niches is needed. Here, we demonstrate a new strategy to do so by dissecting human skin niches in health and diverse inflammatory diseases. We link H&E images, dissociated transcriptome-wide single-cell profiles, and sparse spatially-resolved single-cell profiles at scale, overcoming the technical limitations of the respective technologies. Through this approach, we catalogue and spatially resolve cell types in human skin across ∼5 million cells, decode disease-associated histomorphologic niches, and reveal cellular signalling pathways in AD and psoriasis that likely contribute to their distinct cellular abundance and co-localisation patterns. We identify “novel” niches in inflamed human skin that are not discernible from H&E alone, including a T_RM_-rich sebaceous niche (*Sebaceous_immune*) and a gland-associated *Plasma_cell_rich* niche. These niches redefine the role of tissue appendages, and the cells forming them that were previously considered to have limited immunological activity, such as outer sebocyte duct cells and sweat gland duct cells, highlighting these as key cells that sculpt novel microenvironments in disease settings.

A role for the sebaceous gland in the maintenance of T_RM_ cells has not previously been recognised in humans or mice. Skin T_RM_ cells have predominantly been defined based on flow cytometry^6,33,35^, and thus are not well localized across different skin niches. While lymphoid-sebaceous cell interactions have been reported in murine studies, this has only been in the context of sebaceous gland size and sebum secretion^109,110^. We thus define a new association for the sebaceous gland unit with inflammatory memory, identifying this structure as a likely microenvironment to maintain T_RM_ cells given its unique concentrations of lipids and retinoic acid in the skin^82,111^. Furthermore, the sebaceous gland is clinically linked to AD as seborrhoeic dermatitis frequently precedes early-onset AD^112^; follicular AD is a recognised clinical phenotype of AD^113^; and AD affecting the head and neck region, which is characterised by a high sebaceous gland density, is particularly difficult to treat^114,38^. Previously, epidermotropism has been reported to be mediated by the suprabulbar and infundibular keratinocytes of the hair follicle in mice^77^. Using our single-cell resolution spatial map of skin, we pinpoint sebocytes in the outer layer of the sebaceous duct as a major source of *IL15*, as well as of the chemokine *CCL19*, in the human epidermis, raising the possibility of the sebaceous gland/duct as a site of epidermotropism in skin.

Plasma cells contribute to a B cell memory wall that has classically been associated with secondary lymphoid organs^88^. A major conceptual advance, established in mice, was that B cell antibody responses can occur *de novo* in skin, independent of secondary lymphoid organs^90,115^, with bacterial colonization of skin shown to induce IgG antibody production from skin autonomously. This finding of autonomous antibody production was leveraged to induce protective inflammatory memory through application of select antigen to the skin surface^115^. Whether this same effect occurs in humans is uncertain. Our work raises the possibility of a similar function in humans, with an impaired skin barrier in AD contributing to development of the *Plasma_cell_rich* niche. We also identified evidence that the gland-associated *Plasma_cell_rich* niche in human skin was analogous to the plasma cell-rich GAIN in human lung^93^, and potentially intestine^116^, suggesting that this niche represents a cross-tissue immune niche. While GAIN was only studied in non-diseased lung, this niche was suggested to play an important role in common lung diseases^93^.

Through defining hidden immune niches in human skin, we were able to identify enrichment of candidate genes mediating genetic risk of inflammatory skin disease within these regions. Until now, such genetic variant analyses by space have utilised broad tissue localisations or histopathological annotations^9,10^, and thus do not capture hidden disease-associated immune niches that are likely critical to inflammatory memory formation.

Our study provides a foundational resource for future studies of skin and other tissues, at a cell and tissue niche level. While AD and psoriasis have been relatively extensively studied^5,34,71,73,118–123^, many other skin diseases remain sparsely profiled or completely unprofiled. Our data and workflow will allow researchers to achieve much more granular cell and niche annotations in existing and newly generated skin data than was previously achievable, which we illustrate in two external skin diseases. Through generation of spatial transcriptomic data and reference-mapping, we propose that “hidden” niches will be found across skin diseases, heralding a new era of disease understanding beyond traditional histopathology. Our work also provides an *in situ* atlas of human skin at scale, linked to high-resolution cell types and histopathology, which has relevance for parallel fields through comprehensively defining the spatial architecture of skin. This resource can also be used for studying cross-tissue mechanisms through our comprehensively profiling of disease. For instance, we resolved and identified potential interactions between rare *F6: Inflammatory myofibroblasts* and *Eosinophil* populations. *F6: Inflammatory myofibroblasts* have not been previously reported in type 2 inflammation, but an *F6: Inflammatory myofibroblast* marker (*IL13RA2*) has been identified in the broad fibroblast population in eosinophilic esophagitis^125^, suggesting relevance of this cross-tissue^51,75^ inflammatory myofibroblast population in other diseases and a shared mechanism of eosinophil recruitment.

A strength of our study is that scRNA-seq and Xenium data are integrated and defined with the same marker genes, providing *in situ* transcriptomic validation of scRNA-seq findings and confidence that reported cell types do not arise from dissociation effects. We also profile skin at scale with spatial transcriptomics (>150 sections), allowing us to capture even rare niches. A limitation is that Xenium had lower resolution for cell types due to lower sensitivity of profiling genes, and thus some cells could not be captured or assigned a specific cell type. We also included only week 12 data for AD and thus cannot determine the longevity of the *Plasma_cell_rich* niche. For reported candidate genes from GWAS, we could only assess genes included in the Xenium-5k panel, and thus do not fully capture all genetic variants.

Overall, our study provides the most comprehensive spatially-resolved profile of human tissue to date, including >5 million cells in total and >150 spatial transcriptomic slides. Our work represents an important step, using skin as an exemplar, towards a systematic understanding of human tissues in health and disease through integration of histopathology, atlas-scale genomics, and AI-based frameworks. A comprehensive end-to-end decomposition of human tissue into modular niches and deep understanding of their resultant cellular and molecular properties provides a basis to systematically transform histopathological interpretation of disease to advance therapeutic strategies.

---

**Extended Data Figure 1.**
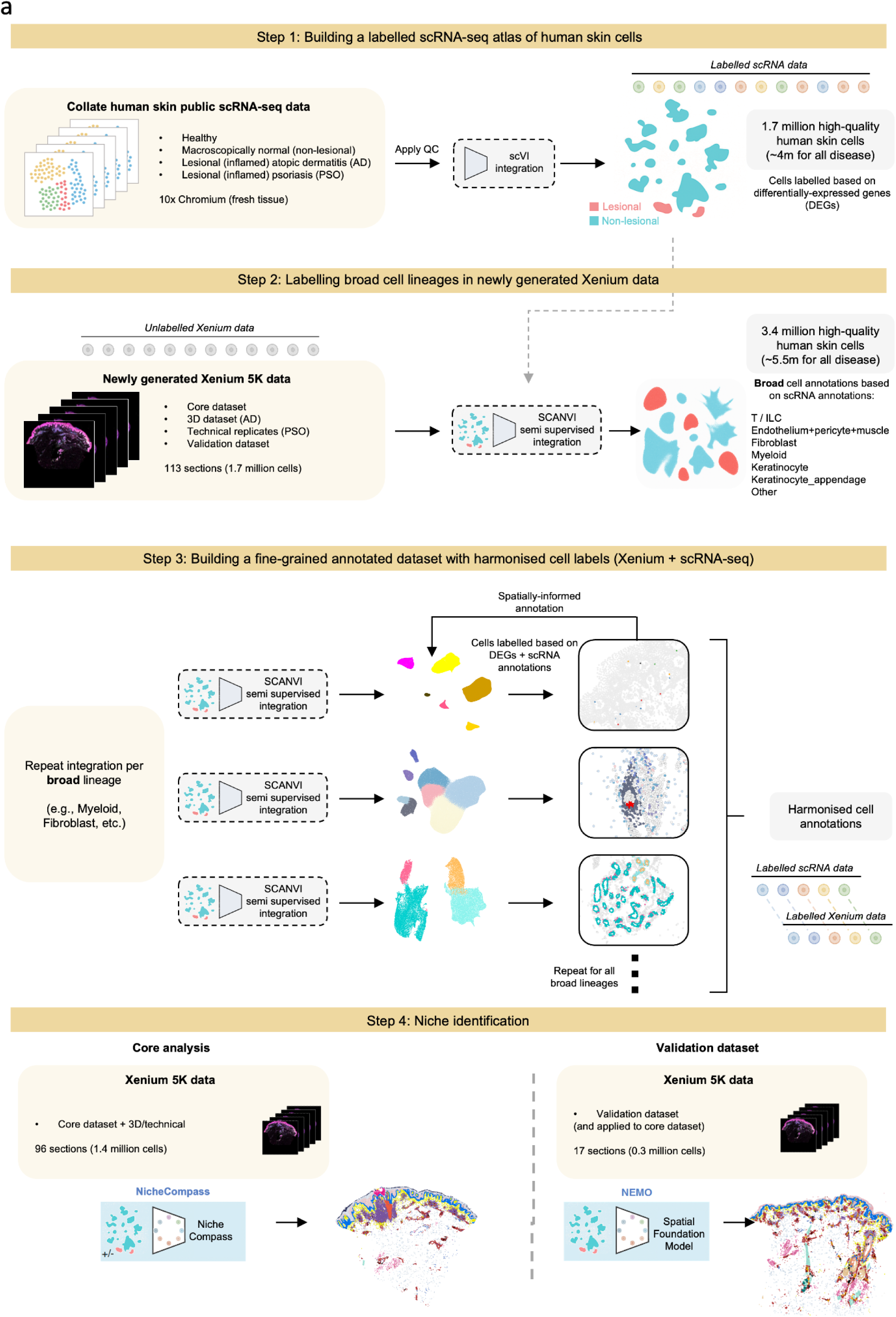
Study overview for joint integration and niche identification. **a,** Overview of study methodology and datasets for skin atlas integration to identify harmonised cell types across modalities and identify niches within Xenium-5k data.

**Extended Data Figure 2.**
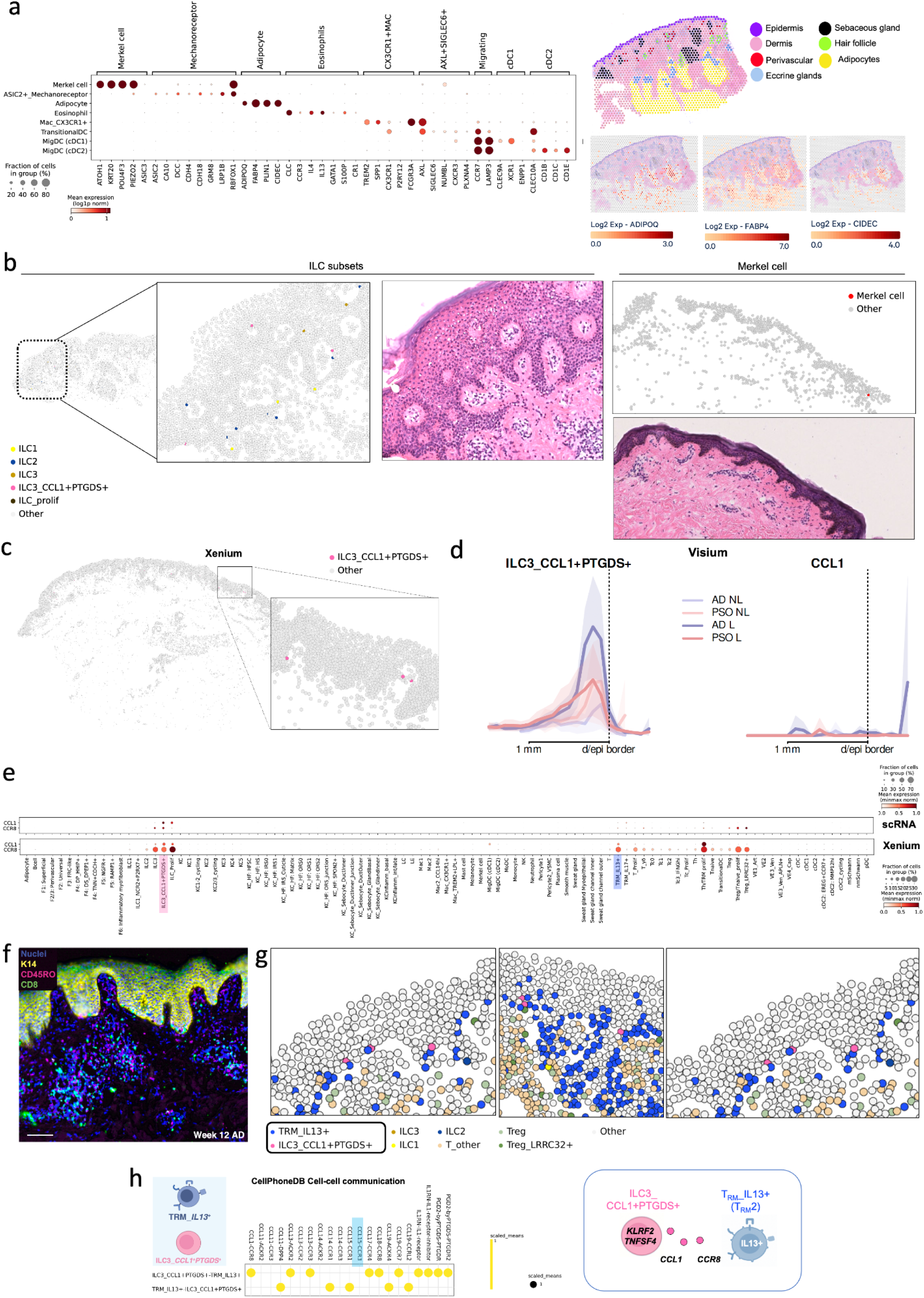
Skin atlas high-resolution cell types. **a,** Dot plot of rare cell types in skin (left) and validation of adipocyte genes using Visium data (right). **b,** Human skin tissue (AD lesional) profiled by Xenium 5k coloured by cell type, with matching haematoxylin and eosin stain, for Merkel cells and ILC subsets.**c** Human skin tissue (AD lesional) profiled by Xenium 5k showing *ILC3_CCL1^+^PTGDS^+^* location. **d,** VisUtils plot of *ILC3_CCL1^+^PTGDS^+^* cell type abundance (left) and *CCL1* expression by tissue depth (right) (see Methods). Mean +- 2 standard deviations of mean. **e,** Dot plot of expression of *CCL1* and *CCR8* across all cell types in Xenium and scRNA-seq dataset. **f,** AkoyaPhenocyler profiling of AD skin post-treatment (week 12), showing persistence of CD8+ T cells at week 12. Scale bar = 100 µm. **g**, Human skin tissue (AD lesional) profiled by Xenium 5k showing *ILC3_CCL1^+^PTGDS^+^* and *T_RM__IL13+ loc*ation, suggesting co-localisation. Neighbourhood enrichment matrix in Supp. Fig. 12b . **h,** CellPhoneDB results for interactions of *T_RM__IL13^+^* and *ILC_CCL1^+^PTGDS^+^* cell types. AD: atopic dermatitis. ILC: innate lymphoid cell. scRNA-seq: single-cell RNA sequencing, T_RM_: Resident memory T cell

**Extended Data Figure 3.**
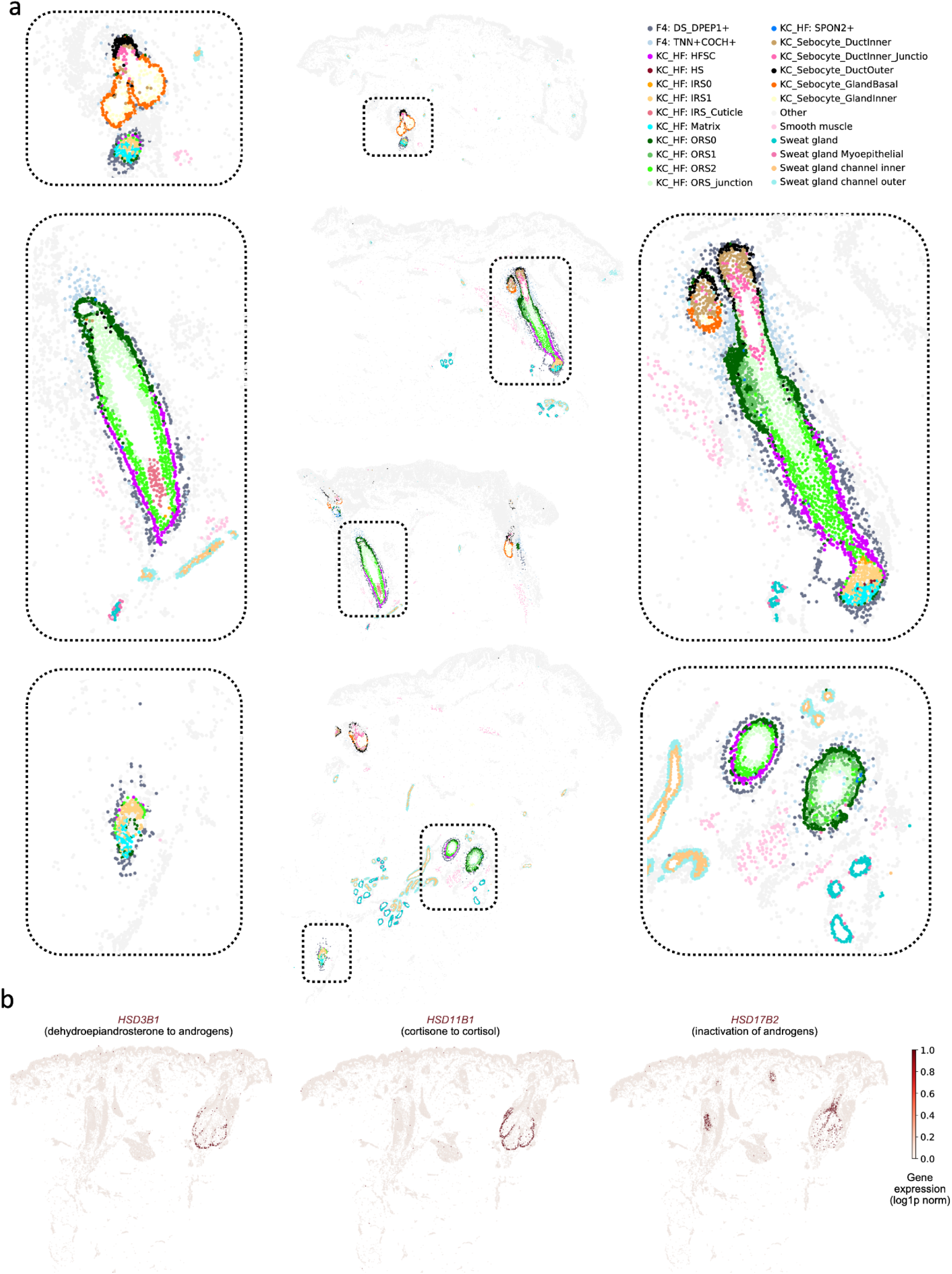
Hair follicle subtypes in skin and distinct localization of enzyme expression within the sebaceous gland. **a,** Xenium 5k coloured by keratinocyte appendage cell types. **b,** Xenium 5k coloured by gene expression showing spatial restriction of biochemical processes related to androgen activation and inactivation.

**Extended Data Figure 4.**
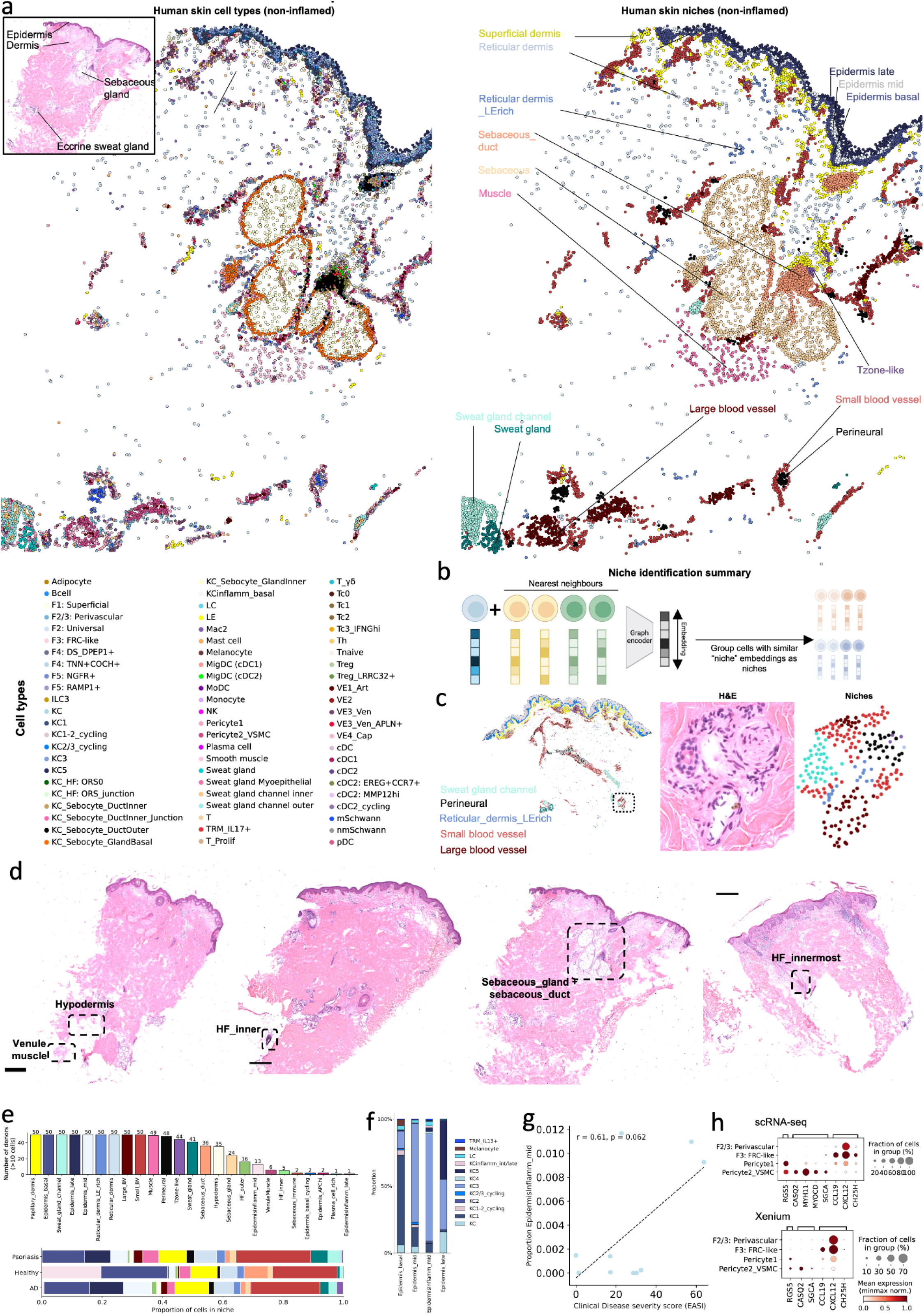
Non-lesional niche composition. **a,** Xenium 5k (non-lesional), with matched H&E stain (top left), coloured by cell type (left) and niche (right). **b,** Simplified schematic of approach to generate niche embeddings, in which input is both the index cell and its set of *k* neighbours. **c,** Xenium 5k slide coloured by tissue niches, where dashed box shows neurovascular bundle (left). Zoomed in H&E stained tissue and Xenium tissue capturing the *Perineural*, *Small_Blood_Vessel*, and *Large_Blood_Vessel* niches within the neurovascular bundle, in addition to a nearby *Sweat_gland_channel*. **d,** Full tissue H&E sections showing the location of niches shown in Fig. 2b. Scale bar = 500µm. **e,** Bar plot showing number of donor samples that contain the niche (minimum 10 cells, hence HF_innermost not shown) (top) and bar plot showing proportions of tissue niche composition by patient status (psoriasis non-lesional, AD, healthy (no inflammatory skin disease but aged)). Healthy skin contained the same niches but with expanded *EpidermisInflamm_mid*, which may reflect this was aged skin or different processing (FFPE vs frozen). **f,** Proportion of cell types within each epidermal niche in non-lesional skin. **G,** Correlation between inflammation severity score (EASI) and number of cells annotated as *EpidermisInflamm_Mid* niche in non-lesional skin by donor. Pearson’s correlation (two-tailed) with fitted least-squares regression line shown. **h,** Expression of chemotaxis-related genes (*CCL19*, *CXCL12*, *CH25H*) and muscle-related genes (*MYH11*, *MYOCD*, *SGCA*) in stromal cells within blood vessel niches, showing increased expression of chemotaxis genes in populations enriched in *Small_Blood_Vessel* niche (*F2/3: Perivascular*, *F3: FRC-like*, and *Pericyte1*) compared to the *Large_Blood_Vessel* niche (*Pericyte2*). AD: atopic dermatitis, EASI: eczema activity and severity index, FFPE: Formalin-Fixed Paraffin-Embedded. H&E: haematoxylin and eosin.

**Extended Data Figure 5.**
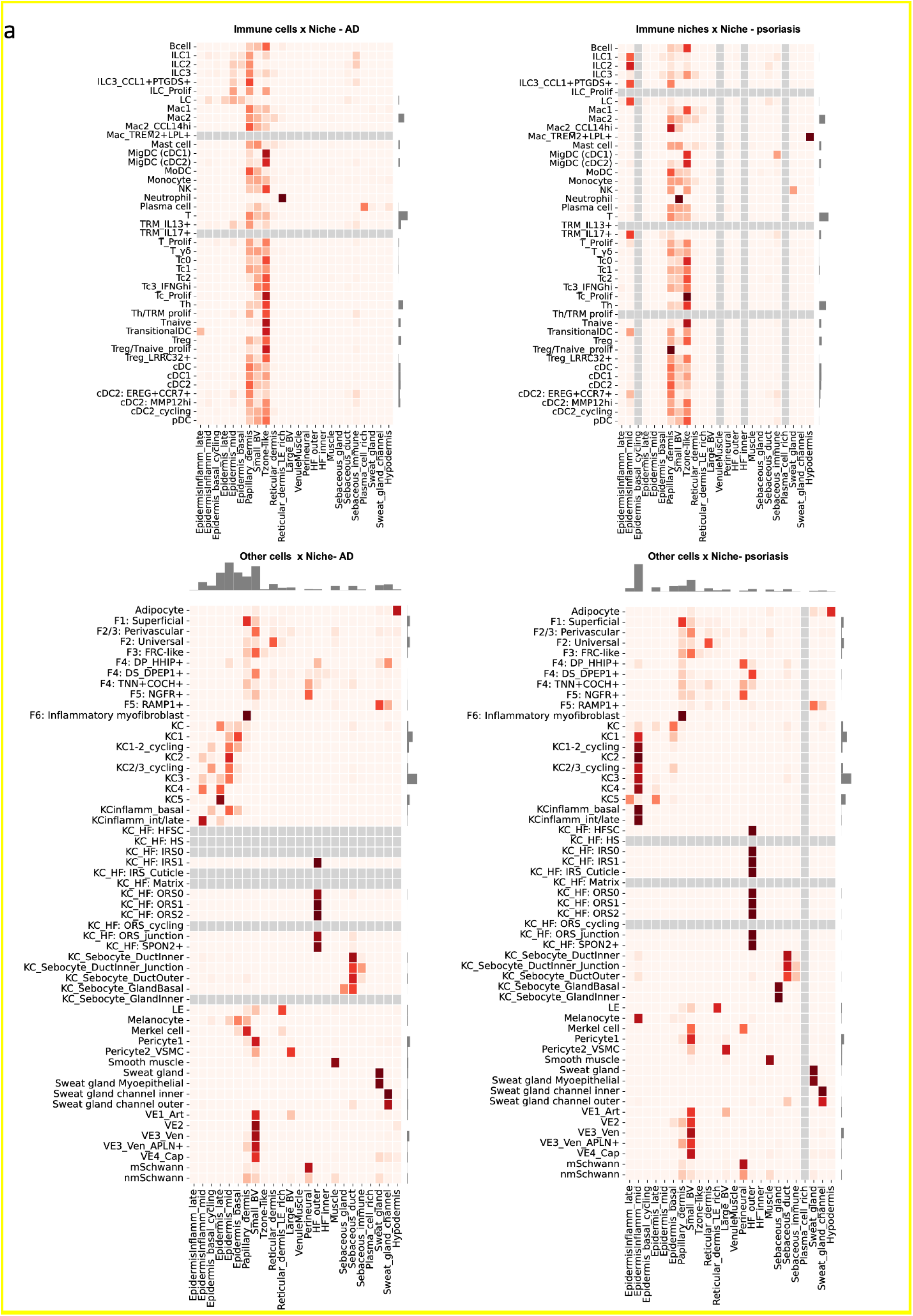
Cellular distribution in inflammation. **a,** Heatmap showing cell type distribution by niche for immune cells (top) and non-immune cells (bottom) for lesional AD and lesional psoriasis skin in Xenium 5k data, normalised by row. AD: atopic dermatitis.

**Extended Data Figure 6.**
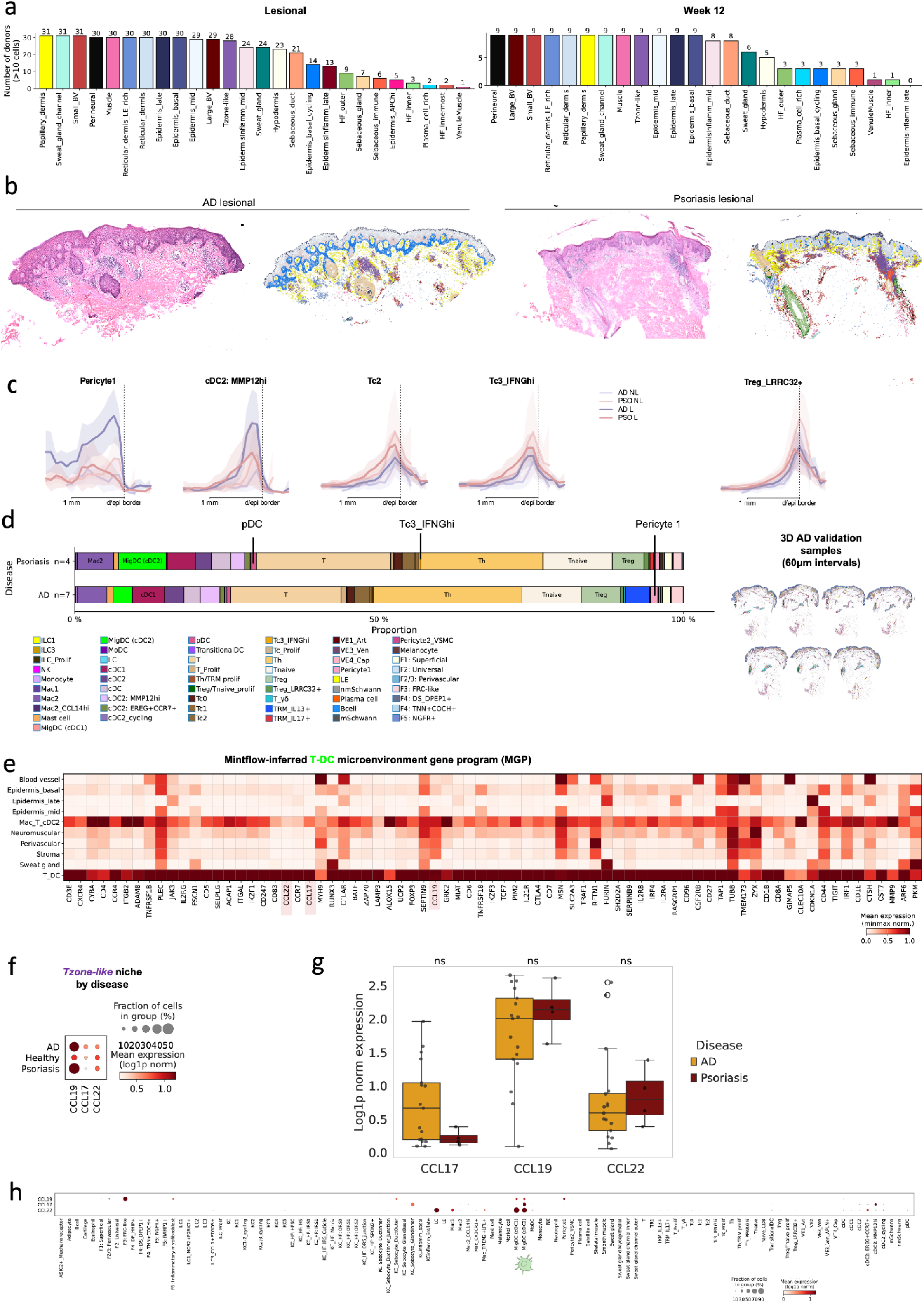
Tzone-like niche composition. **a,** Number of donor samples with each niche in lesional and week 12 samples. **b,** H&E stained tissue of inflamed AD (left) and psoriasis (right) with Xenium 5k of same tissue coloured by niche.**c,** VisUtils plots showing cell type abundance by tissue depth. While *Treg* cells showed a significantly increased abundance in AD (see Fig. 3d), *Treg_LRRC32+* cells did not. **d,** Composition of *Tzone-like* niche in 11 further sections not included in original analysis: we used 3D samples for AD (i.e. consecutive sections at 60µm intervals from the same tissue block) (shown to right of plot, coloured by cell type) and technical replicates for psoriasis (see Extended Data Fig. 1 for overview of datasets).**e**, Expression of genes in microenvironment-induced gene programme corresponding to *Tzone-like* niche in AD, highlighting chemokines. **f,** Dotplot of *CCL17*/*CCL22*/*CCL19* expression in Tzone-like niche by disease status. **g,** Boxplot of *CCL17*/*CCL22*/*CCL19* expression in *Tzone-like* niche by disease status. Statistical significance (Mann-Whitney U for donor-level expression) was annotated as * for p < 0.05 and ns (not significant) otherwise. Differences were non-significant across all cells within the *Tzone-like* niche, and we therefore later performed differential gene expression testing only in *MigDC (cDC2)* cells (see Fig. 3g) after localising the expression of *CCL17* to this population. **h,** Expression of chemokines from *Tzone-like* microenvironment-induced gene programme across all cell types (scRNA-seq data). AD: Atopic dermatitis. H&E: Haematoxylin and eosin scRNA-seq: single-cell RNA sequencing

**Extended Data Figure 7.**
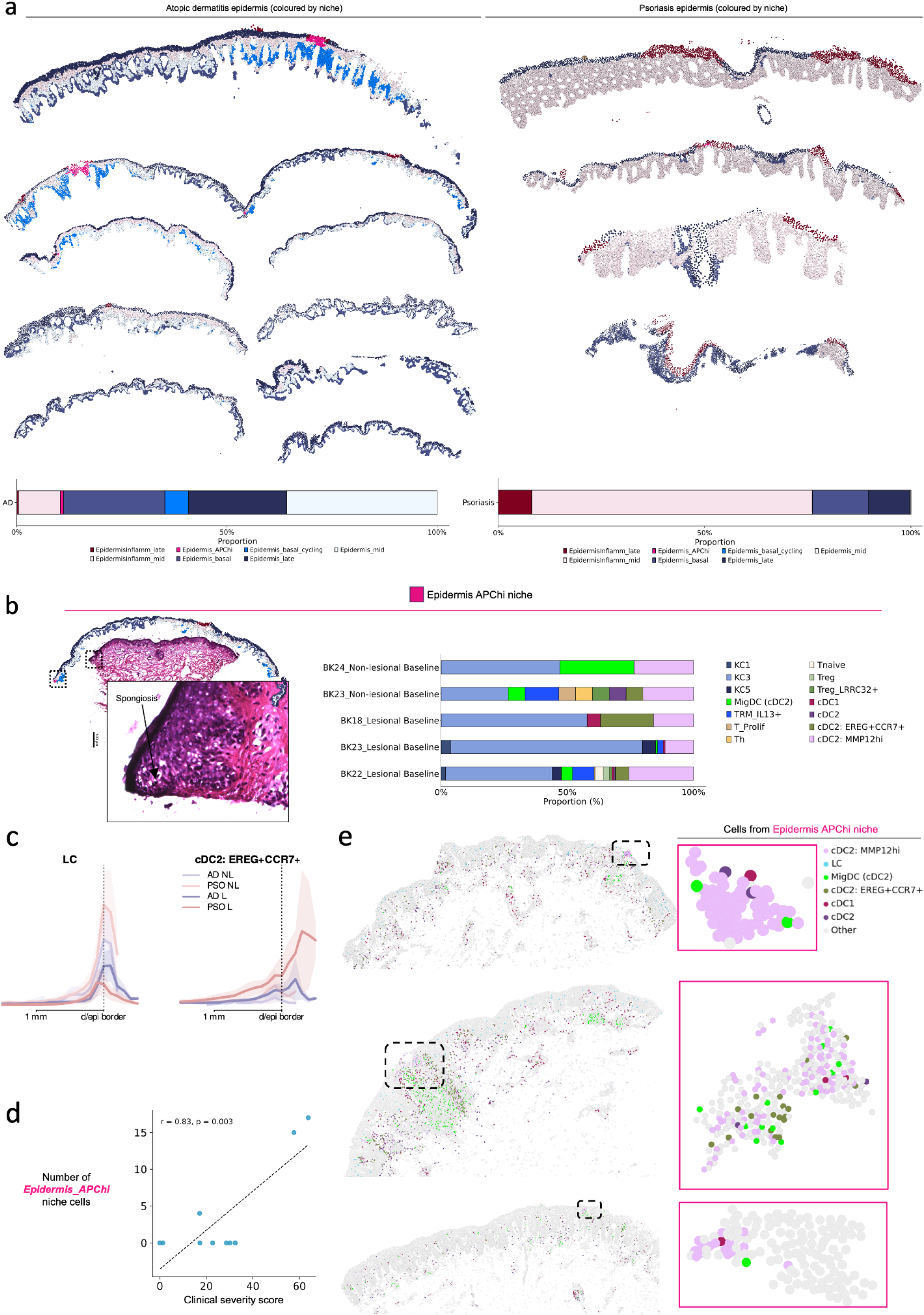
Epidermis during inflammation in atopic dermatitis and psoriasis. **a,** Xenium 5k coloured by niche (epidermal niches only) for AD and psoriasis core dataset (not 3D or validation data) and proportion of epidermal composition by niche. **b,** Xenium 5k samples containing *Epidermal_APC^hi^* niche and their cell composition (right). Haematoxylin and eosin-stained tissue and Xenium 5k section, coloured by niche, for lesional AD skin sample with *Epidermal_APC^hi^* niche that is not shown in Fig 3h, showing spongiosis corresponds to the *Epidermal_APC^hi^*niche (dotted box) (left). **c,** VisUtils plot of Langerhans cell (*LC*) and *cDC2: EREG+CCR7+* cell abundance by tissue depth. Mean +- 2 standard deviations of mean. **d,** Correlation between inflammation severity score (EASI) and number of cells annotated as *Epidermal_APC^hi^*niche in non-lesional skin by donor. Pearson’s correlation (two-tailed) with fitted least-squares regression line shown. **e,** Lesional Xenium 5k samples containing *Epidermal_APC^hi^* niche, showing DC populations. Dashed boxes highlight *Epidermal_APC^hi^* niche. Inlet shows *Epidermal_APC^hi^* niche cells only, coloured by cell type for DCs (vs other in grey). AD: atopic dermatitis. APC: antigen-presenting cell. DC: dendritic cell. EASI: Eczema Area and Severity Index

**Extended Data Figure 8.**
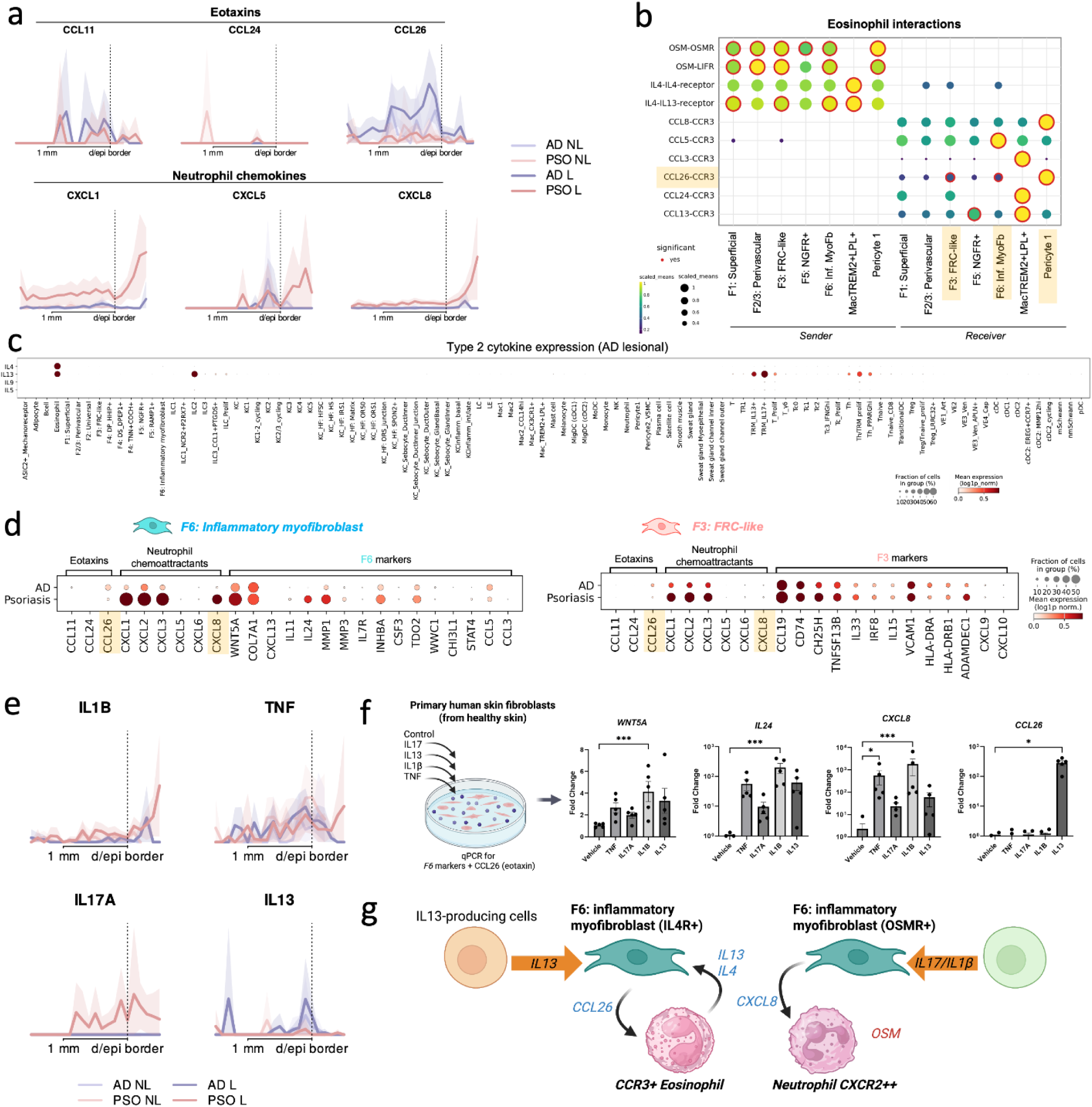
Inflammatory myofibroblasts recruit distinct granulocytes in AD and psoriasis. **a,** VisUtils plot of eotaxin and neutrophil-attracting chemokine expression by tissue depth. Mean +- 2 standard deviations of mean. **b,** Cell-cell communications results for eosinophils with other cell types. **c,** Dotplot of type 2 chemokine expression by all cell types (scRNA-seq data), showing uniquely high *IL4* expression in eosinophils. *IL5* was not highly expressed, and consistently, anti-IL-5 agents have not shown clinical efficacy in treating AD. **d,** Dotplot of eotaxins, neutrophil-attracting chemokine expression, and fibroblast subtype marker genes in scRNA-seq data in *F6: Inflammatory myofibroblasts* and *F3: FRC-like fibroblasts*. **e,** VisUtils plot of panel used for fibroblast stimulation by tissue depth. Mean +- 2 standard deviations of mean. **f,** Schematic of fibroblast stimulation experiment (left). qPCR of inflammatory myofibroblast markers and eotaxin from primary human fibroblasts stimulated with TNF, IL-17A, IL-1β and IL-13. n = 5 biologically independent samples; data shown as mean ± SEM; * P < 0.05, ** P < 0.01, *** P < 0.001 (Friedman test with Dunn’s post-test). Fold change calculated relative to GAPDH. **g,** Schematic summary of different polarisation of *F6: Inflammatory myofibroblasts* by chemokine milieu. AD: atopic dermatitis. qPCR: quantitative polymerase chain reaction. scRNA-seq: single-cell RNA sequencing. SEM: standard error of mean.

**Extended Data Figure 9.**
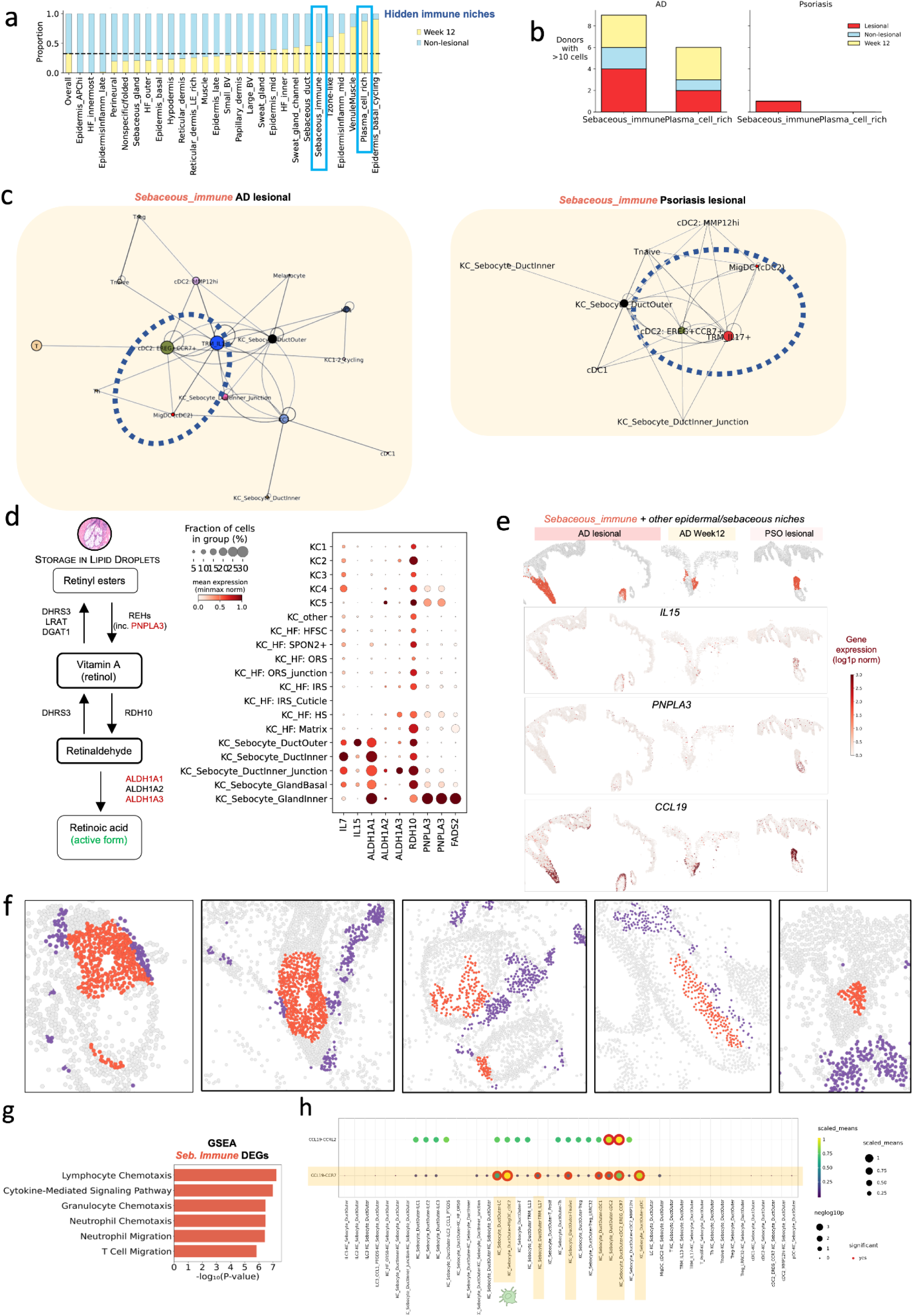
Hidden immune niches. **a,** Bar plot showing proportion of cells from non-lesional and week 12 skin for each niche. Far left bar indicates overall composition. **b,** Barplot showing number of tissue sections containing the hidden immune niches by disease and site status (non-lesional, lesional, week 12). **c,** “Connectosomes”, which were recently utilized^40^ to provide a network representation of cell type co-localisation enrichments within a given niche, averaged across samples (see Methods) for the *Sebaceous_immune* niches in AD and psoriasis. Nodes represent cell types. Edges between cell types indicate cell types that co-localise / are found in close proximity. **e,** Schematic of genes involved in retinoic acid generation (left). Expression of these genes in epidermal cell types in scRNA-seq data (right). **f,** *Sebaceous_immune* niche and other epidermal niches (top row). Expression of chemokines in Xenium 5k data within these regions, showing enrichment of *IL15* and *CCL19* in *Sebaceous_immune* niche. *PNPLA3* expression was highest in the sebaceous gland. **g,** Gene set enrichment analysis, using differentially expressed genes for the *Sebaceous_immune* niche, identified the most enriched processes in the *Sebaceous_immune* niche related to chemotaxis of immune cells. **h,** Cell-cell communications results showing significant interaction for *KC_Sebocyte_OuterDuct* cells (CCL19+) and *MigDC (cDC2)* (CCR7+) cells. AD: atopic dermatitis.

**Extended Data Figure 10.**
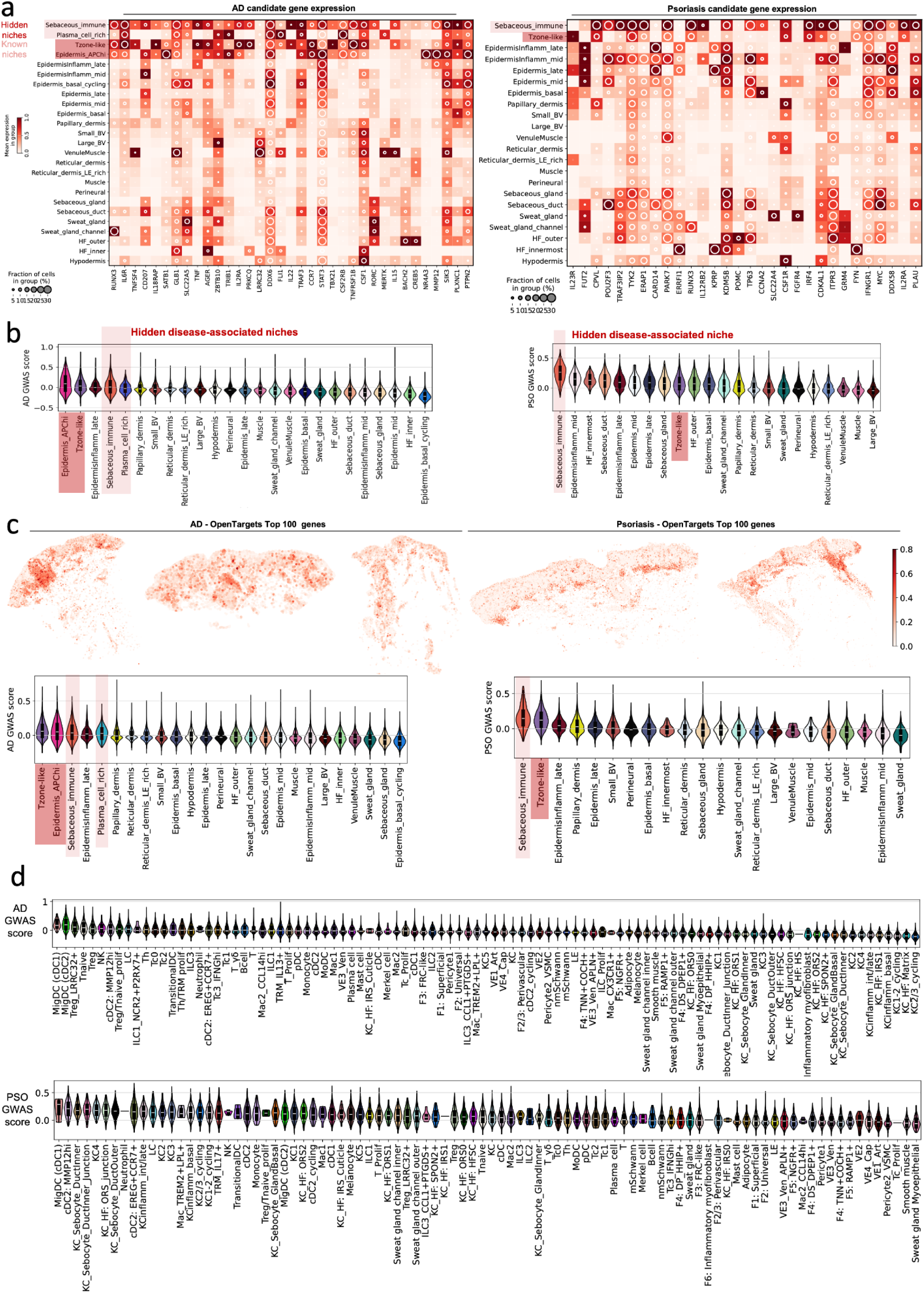
Genetic variants within skin cell types and niches. **a,** Dotplot-heatmap combined of candidate genes for disease susceptibility genes (from genome wide association studies (see Methods)) for AD and psoriasis by tissue niche. **b,** Violin plots of gene module scores (calculated from candidate gene list) in tissue niche, sorted from highest scores, in AD (left) and psoriasis (right). **c,** Spatial plots of gene module scores for each cell calculated using candidate genes for AD and psoriasis from OpenTargets gene set (Methods) (top). Violin plots of gene module scores (calculated from OpenTargets gene set) in each niche, sorted from highest scores. **d,** Violin plots of gene module scores (calculated from gene set derived from genome wide association studies) in each cell type by disease, sorted from highest scores.

## Online methods

### Human tissue samples

Human adult skin biopsies from healthy volunteers or atopic dermatitis (AD) and psoriasis patients processed for data generation in this study were collected as 6mm punch biopsies. Clinical diagnosis was made by dermatologists. Samples were then processed into formalin fixed paraffin embedded (FFPE) for healthy skin samples or fresh frozen OCT-embedded for AD and psoriasis samples for further downstream tissue processing. All research ethics committees and regulatory approvals were in place for the collection and storage of research samples at St John’s Institute of Dermatology, Guy’s Hospital, London (REC reference number: EC00/128).

### Clinically severity scores

At the time of biopsy, for patients with AD, disease severity scores were clinically rated using the validated eczema area and severity index (EASI) score^126^. We defined a treatment responder as a >50% improvement in EASI score as this is frequently used in AD clinical trials.

### Data generation

#### Xenium

OCT-embedded fresh frozen AD or psoriasis human skin tissue was used to generate 10µm sections. FFPE healthy human skin tissue was used to generate 5µm sections. We generated *in situ* gene expression data generation using the 10x Genomics Xenium *in situ* 5k-plex platform. Tissue sections were placed into the active capture area on Xenium slides and stored accordingly until being processed for *in situ* gene expression according to the manufacturers’ protocol. All Xenium slides were processed using the Xenium Prime 5K Human Pan Tissue & Pathways Panel, including the standard cell segmentation antibody staining. Post Xenium imaging slides were kept in PBS at 4°C to continue with post-Xenium H&E immunohistochemistry and/or multiplex protein immunofluorescence using the Akoya Phenocycler-Fusion platform.

#### Selection of validation dataset

To generate a validation dataset on the 10X Xenium and Akoya Phenocycler-Fusion platforms, we selected 10 skin tissue blocks from patients profiled in our original cohort to further investigate niche identification. We generated consecutive tissue sections from these blocks (10μm). We selected two sections with the *Sebaceous_immune* niche and three sections with the *Plasma_cell_rich* niche, we also selected five further sections: a section which had low quality in the initial run, a hair follicle section, a non-lesional AD section, and a further lesional AD section and lesional psoriasis section to create our Xenium validation dataset.

#### Immunofluorescence validation with Akoya PhenoCycler-Fusion

Following processing for 10X Xenium Prime 5K Gene Expression workflow, slides were then stained following the PhenoCycler-fusion one step staining protocol for the human IO 60-plex panel in accordance with the manufacturer’s protocol, but without additional acetone and PFA pre-antibody fixation. The PhenoCycler-Fusion system then performs consecutive cycles of dispensing reporters by hybridising the complementary barcodes then acquiring images and removing the reporters using an isothermal wash until all markers are acquired. At the end of the assay, whole slide scans are acquired and were processed using the QuPath software to visualise individual markers as well as co-expression. Images were exported and saved in TIFF format.

#### Visium

We generated spot-based spatial whole transcriptome data using the 10x Genomics Visium (v1 3’) Spatial Gene Expression platform from frozen OCT-embedded human adult skin tissue. Skin samples were sectioned at 15 µm thickness and the optimal tissue permeabilisation time was determined to be 14 minutes for these samples. H&E images were taken using a Hamamatsu NanoZoomer S60v2 Digital slide scanner at 20X magnification. Spatial gene expression libraries were sequenced using an Illumina NovaSeq 6000 to achieve a minimum number of 50,000 read pairs per tissue covered spot.

#### Fibroblast in vitro stimulation

Primary dermal fibroblasts were isolated from healthy skin donated following plastic surgery (REC reference number: 06/Q0704/18). Fibroblasts were grown in Dulbecco Modified Eagle Medium (DMEM) supplemented with 10% fetal bovine serum (FBS) and 1% penicillin-streptomycin (all reagents from Gibco). Cells (50,000 per well) were incubated in DMEM with 1% FBS for 24 hours before cytokine stimulation using 5ng/mL TNF (R&D Systems), 50ng/mL IL-17A (R&D Systems), 1ng/mL IL-1β (R&D Systems), 100ng/mL IL-13 (PeproTech) or complete DMEM (vehicle) for 24 hours at 37°C. Cells were harvested for RNA isolation (QIAGEN) and quantitative PCR was performed to assess gene expression using primers reported in Supplementary Table 4. Transcript levels were normalised to the expression of GAPDH and calculated using the 2Δct method. Data were analysed in GraphPad Prism (version 10.5.0 for Windows) using a Friedman test followed by Dunn’s multiple comparison test.

### Computational analysis

#### Single cell: Atlas data preprocessing and integration

A summary of our processing and integration strategy is shown in Supplementary Fig. 2. Raw scRNA-seq data were downloaded and aligned using STARsolo (GRCh38-2020-A reference) unless already available in local storage^5,127^, and with the exceptions of an AD dataset, provided by authors^71^, and a bullous pemphigoid dataset (non-lesional data used), for which raw counts were not available (Supplementary Table 1). We included publicly-available data generated from fresh skin biopsies using the 10x Genomics Chromium platform.

CellBender v.0.3 was used to correct for ambient mRNA^128^. To remove low-quality cells, we included only cells with >200 genes, >1000 and <300,000 total unique molecular identifiers (UMIs), and a mitochondrial gene percentage of <15%. We calculated doublet scores using scrublet on a per sample basis using scrublet and removed cells with a doublet score of >0.3^129^.

Integration of all cells was performed using single-cell Variational Inference (scVI) using raw counts^23^. We identified highly-variable gene (HVG) per batch using sample ID as the batch key, retaining those expressed in at least 80 batches and targeting ∼6,000 HVGs. Known stress- or hypoxia-related genes were excluded from HVG selection to reduce confounding signals. We also did not consider (for HVG selection) mitochondrial genes; cell cycle genes; and ribosomal genes. In future analyses post-integration, all genes were considered.

The following hyperparameters were used for the scVI model: number of layers: 2; number of latent dimensions: 30; gene likelihood: zero-inflated negative binomial distribution, and dispersion: gene-batch. An early stopping patience of 5 epochs was used. The batch key was “sampleID” and no other covariates were passed to the model for correction.

We constructed a k-nearest neighbours (kNN) graph (k=20) using the scVI embedding and performed community detection (Leiden algorithm) with resolution 1 for the dataset with all skin cells.

#### Differentially expressed gene analysis

For each cluster, we calculated differentially expressed genes using the t-test with the scanpy *rank_genes_groups* function (t-test). Marker genes for each population are shown in Supplementary Figures 3-11, showing gene expression values post-normalisation. We used the shifted logarithm with a scaling factor of 10,000 based on strong performance in a recent benchmarking paper^130^.

For differential expression analysis by cell type (for *MigDC (cDC2)*), we performed pseudobulk differential expression using pertpy and PyDESeq2^131,132^. Lesional samples with ≥100 cells per patient were retained. Counts were aggregated by patient, normalized (10,000 counts per sample), log-transformed, and scaled. Differential expression was tested using the model ∼ disease_overall, comparing psoriasis vs atopic dermatitis, and results visualized via volcano plots. Significance was assessed via Wald tests, and log₂ fold changes were computed for the contrast “Psoriasis vs AD.” For Xenium data, pseudobulk aggregation parameters were

adjusted to min_cells = 20 and min_counts = 1,000 to accommodate lower capture rates and smaller per-sample cell numbers for the fine *MigDC (cDC2)* cell type.

#### Xenium data preprocessing and joint integration

10x Genomics Xenium data was filtered to exclude cells with <10 genes per cell. We then used SCANVI using labelled scRNA-seq and unlabelled Xenium data (Supplementary Fig. 2), restricted to ∼5000 genes found in both datasets. Batch-aware HVG selection was then performed by setting the batch key to sample ID. We selected 2000 HVGs as features for the integration. We used a simpler encoder in the SCANVI model (number of layers: 1; number of latent dimensions: 10). The batch key was sample id (“info_id6”) and a categorical covariate of technology (scRNA vs spatial) was used.

We broadly labelled lineages and repeated feature (HVG) selection and integration by lineage. We then, for each lineage, calculated the kNN graph and performed Leiden clustering. We labelled clusters based on a combination of scRNA-seq annotations and spatial location, with spatial positions helping to refine names for populations such as distinct sebocyte duct and sweat gland channel subsets (Supplementary Fig. 2). Cell coordinates coloured by cell type were visualised using squidpy (<u>sq.pl</u>.spatial_scatter) or scanpy (sc.pl.spatial)^133^. All populations are shown in Supplementary Fig. 2-10 utilising PAGA^134^ initialization for UMAP calculation. For the overall embedding, we repeated SCANI with annotated populations and calculated the kNN graph using k=10. In this model, we excluded the two FFPE samples, leaving 111 samples.

#### Single cell: ACR design

We utilized the ACR design previously described for optimal detection of disease-enriched cell types ^29^. We used non-lesional (excluding AD and psoriasis) data as the atlas control and lesional AD/psoriasis data as query. Hyperparameters are included in the ACR notebook.

For training the non-lesional reference atlas, we used the same pre-processing steps as described above (e.g. ∼6000 HVG selection). Batches with fewer than 2,000 cells

were excluded from HVG consideration to ensure robust gene selection.The model was trained on the reference dataset (non-lesional atlas) with a gene-batch dispersion model for 50 epochs. Then, query datasets were integrated using the scArches framework implemented in scvi-tools. The resulting latent representations were used to compute nearest neighbors (k=20, Euclidean distance) and construct two-dimensional embeddings via UMAP (min_dist = 0.1).

To ensure computational tractability, 10% of cells were subsampled to define Milo neighborhoods. Fine-grained cell annotations were harmonized into seven broad compartments (Myeloid, T/ILC, Endothelium/Pericyte, Keratinocyte, Keratinocyte_Appendage, Fibroblast, and Other). Neighborhoods were counted per sample and tested for differential abundance between psoriasis and atopic dermatitis using a negative binomial generalized linear model. Within each major lineage, fine-grained cell annotations were ordered by mean logFC. Milo differential abundance results were visualized using custom R scripts employing dplyr and ggbeeswarm. Neighborhoods were classified as significant with spatial false discovery rate (Spatial FDR) < 0.1 threshold.

#### Spatial transcriptomics: Xenium niche identification

We utilized 1024 spatially variable genes for NicheCompass^135^ input. The number of neighbours selected per cell was 8. We used a reference-query mapping approach where the reference was healthy/non-lesional and the query data was lesional and post-treatment.

#### Spatial transcriptomics: Connectosomes and neighbourhood enrichment

To calculate neighbourhood enrichment scores, we first constructed an adjacency matrix for indicating which cells were connected using the *spatial_neighbors* functions in squidpy. We determined the neighbourhood as a cell as cells within 10µm of an index cell. We then applied neighborhood enrichment analysis (*nhood_enrichment* function in squidpy) to quantify which cell types were more frequently co-localised than expected by chance. Heatmaps of enrichment scores were visualised using *<u>sq.pl</u>.nhood_enrichment*. To construct connectosomes (shown for *Sebaceous_immune*), we utilised code from ^40^, with a threshold of 0.1 and a minimum cell count per cell type of 20.

#### Spatial transcriptomics: Mintflow

We utilized Mintflow applied to AD spatial transcriptomic data as detailed in ^21^. MGPs were derived utilising the scanpy *rank_genes_groups* function (t-test) applied to spatial clusters. For chemokines within the MGP for the spatial cluster corresponding to the *Tzone-like* niche, we visualised donor-level log1p norm expression values using box plots with *seaborn* and *matplotlib* across disease groups (“Atopic dermatitis” and “Psoriasis”). We included cells from the Tzone-like niche (lesional samples) and excluded remaining donors with fewer than 10 cells in this niche. Donor-level expression between AD and psoriasis was compared using two-sided Mann–Whitney U tests.

#### Single-cell: cell-cell communication analysis

We used spatial transcriptomic data to identify cell types within a niche, and then subset scRNA-seq to only include these cell types. We used CellPhoneDB v5 (method 2) for cell-cell communication analysis^136^. We visualised the results using ktplotspy.

#### Spatial transcriptomics: Gene set enrichment analysis by niche

Gene set enrichment analysis was implemented using Enrichr via gseapy^137^. We used the top differentially expressed genes in the Sebaceous_immune niche. Analysis was performed using the GO Biological Process 2023 database. Adjusted p-values < 0.05 were considered significant. Results were aggregated across gene lists, ranked by significance, and the top six enriched pathways were selected for visualization.

#### Spatial transcriptomics: Spatial foundation model

To extract niche embeddings for validation data, we utilized a spatial foundation model based upon the I-JEPA architecture (to be made available)^138^. This model is transformer-based and thus utilises a distinct approach to NicheCompass, allowing

us to validate the existence of our reported niches with a novel methodology. Benchmarking of this method is ongoing (to be updated), but we compare broadly to tissue sections with niches defined by NicheCompass (Supplementary Fig. 18b) to show that key niches are still recovered: *Tzone-like*, *Sebaceous_immune*, and *Plasma_cell_rich*.

#### Selection of retinoic-acid and fatty acid genes

*PNPLA3* has been reported to promote the extracellular release of retinol through hydrolysis of retinyl esters^139^. Retinoic acid activating genes were taken from multiple, distinct sources^83,84^. For fatty acid genes, we selected genes reportedly associated with sebaceous lipogenesis^140^, with presentation of genes enriched in the sebaceous gland relative to interfollicular keratinocyte populations, as well as genes associated with lipogenesis more broadly^106^.

#### Spatial transcriptomics: GWAS candidate gene module scores

We used candidate genes reported from recent GWAS papers^100–102^ that were available in the Xenium-5k panel. To calculate gene scores, we used the score_genes() function in scanpy. We used arguments of ctrl_size=1000 and n_bins=25. This generated a gene module score for each cell. We then plotted tissue sections coloured by this gene module score, and aggregated scores by niches in violin plots.

For the alternative gene set, we used reported genes for “atopic dermatitis” and “psoriasis” from the Open Targets platform (https://platform.opentargets.org/disease/)^103^. We exported genes with genetic association with the disease, and selected the top 100 genes sorted by global score.

#### Reference-mapping: Alopecia areata, acne vulgaris, and skin T cell atlas

We utilised unseen lesional (inflamed) samples for alopecia areata and acne vulgaris^104,106^. Inputting raw count data, we utilized our workflow for joint integration with our atlas data available on github. We used the scRNA-seq atlas of AD, psoriasis, and non-lesional skin (∼1.7m cells). We applied this same approach to create the skin T cell atlas, but including lesional (inflamed) samples from skin inflammatory and/or fibrotic disorders. We then selected specific clusters for downstream processing (kNN graph and Leiden clustering, as described above). Annotations were refined using manual annotation of differentially expressed genes for identified clusters, as well as the abundance of labelled populations. We then confirmed the identity of cell type populations using the same marker genes established for our skin atlas, shown using only the mapped data (excluding atlas).

For acne, we selected genes reported in recent GWAS to assess expression that showed enrichment in specific cell types and were not commented upon in the original study^99,108^.

#### Spatial transcriptomics: visutils

10x Genomics Visium data was mapped using Spaceranger v.1.3.0 using GRCh38-2020-A reference. We used loupe browser to manually annotate spots as epidermis/dermis/hair follicule, eccrine glands/adipose and remove debris and regions with folded tissue. In addition regions with parakeratosis and perivascular infiltrates were annotated. Dermis and adipose regions of skin have low cell density and as result have low read coverage (few hundreds of UMI) that is below most of widely used cutoffs. To be able to utilize these spots we used the following procedure:

1. We grouped spots by 7 into higher level of hexagonal mesh.
2. Within each group we selected all spots with less than 500 UMI, summed their gene counts and assigned resulting counts to either central spot (if its coverage was lower than 500) or to spot with highest original total counts.
3. We removed all spots that have coverage below 500

This approach allowed us to keep many dermis spots while sacrificing some spatial resolution. Then we used the cell2location (v0.1.3) to deconvolute the cell types predicted to be present in a given spot^141^. We utilised our reference scRNA-seq atlas using sample ID as batch_key. Then we performed deconvolution with the following parameters: detection_alpha=20,N_cells_per_location=30, and max_epochs 50_000. Values of all other parameters were kept default. Following the

cell2location tutorial, we used 5% quantile of posterior distribution (q05_cell_abundance_w_sf) as predicted celltype abundances.

In order to assess cell type abundance or gene expression in dependence to distance to dermis to epidermis junction we using the following procedure:

1. We define dermis to epidermis border as spots annotated as epidermis but contacting with dermis.
2. We calculated Euclidean distance from each spot to the nearest border spot and rounded it in interspot distances (100μm). Distances for all non-epidermis spots were multiplied by -1. We call resultant values “tissue depth”: its absolute value gives distance to the border. Positive values are above/superficial to the border (i.e. epidermis in skin) whereas negative values are deep/inferior to the border (dermis in skin)
3. For each sample and each depth we averaged cell type abundance (predicted by cell2location) and log(1+CP10K) gene expression values that resulted in depth*feature*sample matrix where features can be either cell types or genes.
4. These matrices were used to plot cell type and gene to tissue depth profiles.

## Acknowledgements

The Wellcome Sanger Institute is supported by core funding from the Wellcome Trust (grant 220540/Z/20/A). For the purpose of Open Access, the author has applied a CC BY public copyright licence to any Author Accepted Manuscript version arising from this submission. L.S. is supported by a Wellcome Clinical Research Fellowship PhD student grant and Trinity Internal Graduate Studentship. M.H. is funded by Wellcome (grant WT107931/Z/15/Z) and the NIHR Newcastle Biomedical Research Centre. We acknowledge support of the Atopic Dermatitis Research Network HIH grant 1UM1AI151958 for involvement in scRNA-seq data collection (RSG). We thank Aidan Maartens for critical reading of the manuscript, Fereshteh Torabi for helping with figure design, and Tobi Alegbe for assistance with the Open Targets platform. We thank Lily Wright, Francesca Capon, and Andrew Pink for involvement in sample acquisition and feedback on study design. Biorender was used for select figure design.

## Contributions

LS and MH wrote the manuscript. LS, AF, CS, SM, ML, and MH conceived and designed the study. LS primarily conducted the computational analysis. AF, CA, PM, CH, EF, JM, LG, KC, NG, VS, CS, SM, ML, and MH provided regular feedback during the study. PM developed VisUtils and Visium analysis. AF, KR, CT, BR, HC, BO, TJ, DB,and ES designed, performed and analysed wet laboratory experiments. AF, KR, CT, BR, HC, BO, and ES performed Visium and Xenium ST experiments. AF and KR performed the Akoya Phenocycler validation. TJ performed the fibroblast in vitro stimulation experiment. CA, AF, and LS designed figures. AF, VR, and DB coordinated clinical samples. RG and RX assisted with new atopic dermatitis data. SB developed nichecompass and the spatial foundation model. AA developed mintflow. AB, AP, and MP contributed to data collection, including reprocess_public_10x. DBL and DH developed the webportal. All authors read and critically evaluated the manuscript.

## REFERENCES

1. Rood, J. E., Maartens, A., Hupalowska, A., Teichmann, S. A. & Regev, A. Impact of the Human Cell Atlas on medicine. Nat. Med. 28, 2486–2496 (2022).

2. Marella, S. et al. A transcriptomic atlas of healthy human skin links regional identity to inflammatory disease. bioRxivorg 2025.05.10.653184 (2025) doi:10.1101/2025.05.10.653184.

3. Sikkema, L. et al. An integrated cell atlas of the lung in health and disease. Nat. Med. 29, 1563–1577 (2023).

4. Elmentaite, R. et al. Cells of the human intestinal tract mapped across space and time. Nature 597, 250–255 (2021).

5. Reynolds, G. et al. Developmental cell programs are co-opted in inflammatory skin disease. Science 371, eaba6500 (2021).

6. Boyman, O. et al. Spontaneous development of psoriasis in a new animal model shows an essential role for resident T cells and tumor necrosis factor-alpha. J. Exp. Med. 199, 731–736 (2004).

7. Francis, L., Capon, F., Smith, C. H., Haniffa, M. & Mahil, S. K. Inflammatory memory in psoriasis: From remission to recurrence. J. Allergy Clin. Immunol. 154, 42–50 (2024).

8. Corris, P. A. & Dark, J. H. Aetiology of asthma: lessons from lung transplantation. Lancet 341, 1369–1371 (1993).

9. Song, L., Chen, W., Hou, J., Guo, M. & Yang, J. Spatially resolved mapping of cells associated with human complex traits. Nature 641, 932–941 (2025).

10. Kvastad, L. et al. Spatial transcriptomics and genetically implicated genes identify putative causal tissue structures for complex traits. bioRxiv 2025.05.02.651876 (2025) doi:10.1101/2025.05.02.651876.

11. Kong, L. et al. Single-cell and spatial transcriptomics of stricturing Crohn’s disease highlights a fibrosis-associated network. Nat. Genet. 57, 1742–1753 (2025).

12. Qian, X. et al. Spatial transcriptomics reveals human cortical layer and area specification. Nature 644, 153–163 (2025).

13. Farah, E. N. et al. Spatially organized cellular communities form the developing human heart. Nature 627, 854–864 (2024).

14. Restrepo, P. et al. A single-cell spatial transcriptomic census of human skin anatomy. bioRxiv 2025.09.22.676865 (2025) doi:10.1101/2025.09.22.676865.

15. McGregor, C. et al. Spatial fibroblast niches define Crohn’s fistulae. Nature 1–10 (2025).

16. Tian, J. et al. Global epidemiology of atopic dermatitis: a comprehensive systematic analysis and modelling study. Br. J. Dermatol. 190, 55–61 (2023).

17. Greb, J. E. et al. Psoriasis. Br. Med. J. 2, 856–857 (1954).

18. Weidinger, S., Beck, L. A., Bieber, T., Kabashima, K. & Irvine, A. D. Atopic dermatitis. Nat. Rev. Dis. Primers 4, 1 (2018).

19. Griffiths, C. E. M., Armstrong, A. W., Gudjonsson, J. E. & Barker, J. N. W. N. Psoriasis. Lancet 397, 1301–1315 (2021).

20. Birk, S. et al. Quantitative characterization of cell niches in spatially resolved omics data. Nat. Genet. 57, 897–909 (2025).

21. Akbarnejad, A. et al. Mapping and reprogramming tissue microenvironments with MintFlow. bioRxiv 2025.06.24.661094 (2025) doi:10.1101/2025.06.24.661094.

22. Li, T. et al. WebAtlas pipeline for integrated single-cell and spatial transcriptomic data. Nat. Methods 22, 3–5 (2025).

23. Lopez, R., Regier, J., Cole, M. B., Jordan, M. I. & Yosef, N. Deep generative modeling for single-cell transcriptomics. Nat. Methods 15, 1053–1058 (2018).

24. Cabo, R. et al. ASIC2 is present in human mechanosensory neurons of the dorsal root ganglia and in mechanoreceptors of the glabrous skin. Histochem. Cell Biol. 143, 267–276 (2015).

25. Xu, C. et al. Automatic cell-type harmonization and integration across Human Cell Atlas datasets. Cell 186, 5876–5891.e20 (2023).

26. Xu, C. et al. Probabilistic harmonization and annotation of single-cell transcriptomics data with deep generative models. Mol. Syst. Biol. 17, e9620 (2021).

27. Kolter, J. et al. A subset of skin macrophages contributes to the surveillance and regeneration of local nerves. Immunity 50, 1482–1497.e7 (2019).

28. Zein, J. et al. HSD3B1 genotype identifies glucocorticoid responsiveness in severe asthma. Proc. Natl. Acad. Sci. U. S. A. 117, 2187–2193 (2020).

29. Dann, E. et al. Precise identification of cell states altered in disease using healthy single-cell references. Nat. Genet. 55, 1998–2008 (2023).

30. Lotfollahi, M. et al. Mapping single-cell data to reference atlases by transfer learning. Nat. Biotechnol. 40, 121–130 (2022).

31. Kühnel, I. et al. The activating receptor NKp65 is selectively expressed by human ILC3 and demarcates ILC3 from mature NK cells. Eur. J. Immunol. 54, e2250318 (2024).

32. Zareie, P., Weiss, E. S., Kaplan, D. H. & Mackay, L. K. Cutaneous T cell immunity. Nat. Immunol. 1–9 (2025).

33. Cheuk, S. et al. CD49a expression defines tissue-resident CD8+ T cells poised for cytotoxic function in human skin. Immunity 46, 287–300 (2017).

34. Ma, F. et al. Single cell and spatial sequencing define processes by which keratinocytes and fibroblasts amplify inflammatory responses in psoriasis. Nat. Commun. 14, 1–19 (2023).

35. Matos, T. R. et al. Clinically resolved psoriatic lesions contain psoriasis-specific IL-17-producing αβ T cell clones. J. Clin. Invest. 127, 4031–4041 (2017).

36. Alkon, N. et al. Single-cell RNA sequencing defines disease-specific differences between chronic nodular prurigo and atopic dermatitis. J. Allergy Clin. Immunol. 152, 420–435 (2023).

37. Brunner, P. M. et al. Nonlesional atopic dermatitis skin shares similar T-cell clones with lesional tissues. Allergy 72, 2017–2025 (2017).

38. Bangert, C. et al. Dupilumab-associated head and neck dermatitis shows a pronounced type 22 immune signature mediated by oligoclonally expanded T cells. Nat. Commun. 15, 2839 (2024).

39. Bieber, T. Disease modification in inflammatory skin disorders: opportunities and challenges. Nat. Rev. Drug Discov. 22, 662–680 (2023).

40. Reina-Campos, M. et al. Tissue-resident memory CD8 T cell diversity is spatiotemporally imprinted. Nature 1–10 (2025).

41. McCully, M. L. et al. CCR8 expression defines tissue-resident memory T cells in human skin. J. Immunol. 200, 1639–1650 (2018).

42. Gombert, M. et al. CCL1-CCR8 interactions: an axis mediating the recruitment of T cells and Langerhans-type dendritic cells to sites of atopic skin inflammation. J. Immunol. 174, 5082–5091 (2005).

43. Gordon, K. B. et al. A phase 2 trial of guselkumab versus adalimumab for plaque psoriasis. N. Engl. J. Med. 373, 136–144 (2015).

44. Whitley, S. K., et al. Local IL-23 is required for proliferation and retention of skin-resident memory TH17 cells. Sci. Immunol. 7, eabq3254 (2022).

45. Adler, M., Chavan, A. R. & Medzhitov, R. Tissue biology: In search of a new paradigm. Annu. Rev. Cell Dev. Biol. 39, 67–89 (2023).

46. Suárez-Fariñas, M., Fuentes-Duculan, J., Lowes, M. A. & Krueger, J. G. Resolved psoriasis lesions retain expression of a subset of disease-related genes. J. Invest. Dermatol. 131, 391–400 (2011).

47. Pavel, A. B. et al. Tape strips from early-onset pediatric atopic dermatitis highlight disease abnormalities in nonlesional skin. Allergy 76, 314–325 (2021).

48. Renert-Yuval, Y. et al. Tape strips capture atopic dermatitis-related changes in nonlesional skin throughout maturation. Allergy 77, 3445–3447 (2022).

49. Wang, X.-N. et al. A three-dimensional atlas of human dermal leukocytes, lymphatics, and blood vessels. J. Invest. Dermatol. 134, 965–974 (2014).

50. Dyring-Andersen, B. et al. Spatially and cell-type resolved quantitative proteomic atlas of healthy human skin. Nat. Commun. 11, 5587 (2020).

51. Steele, L. et al. A single-cell and spatial genomics atlas of human skin fibroblasts reveals shared disease-related fibroblast subtypes across tissues. Nat. Immunol. 1–14 (2025).

52. Kolter, J. et al. Sensory neurons shape local macrophage identity via TGF-β signaling. Immunity (2025) doi:10.1016/j.immuni.2025.08.004.

53. Yates, K., Bi, K., Haining, W. N., Cantor, H. & Kim, H.-J. Comparative transcriptome analysis reveals distinct genetic modules associated with Helios expression in intratumoral regulatory T cells. Proc. Natl. Acad. Sci. U. S. A. 115, 2162–2167 (2018).

54. Cohen, J. N., et al. Regulatory T cells in skin mediate immune privilege of the hair follicle stem cell niche. *Sci. Immunol*. 9, eadh0152 (2024).

55. Zagorulya, M. et al. Tissue-specific abundance of interferon-gamma drives regulatory T cells to restrain DC1-mediated priming of cytotoxic T cells against lung cancer. Immunity 56, 386–405.e10 (2023).

56. You, S. et al. Lymphatic-localized Treg-mregDC crosstalk limits antigen trafficking and restrains anti-tumor immunity. Cancer Cell 42, 1415–1433.e12 (2024).

57. Cruz de Casas, P., Knöpper, K., Dey Sarkar, R. & Kastenmüller, W. Same yet different - how lymph node heterogeneity affects immune responses. Nat. Rev. Immunol. 24, 358–374 (2024).

58. Renert-Yuval, Y. et al. Biomarkers in atopic dermatitis-a review on behalf of the International Eczema Council. J. Allergy Clin. Immunol. 147, 1174–1190.e1 (2021).

59. Mailloux, A. W. & Young, M. R. I. NK-dependent increases in CCL22 secretion selectively recruits regulatory T cells to the tumor microenvironment. J. Immunol. 182, 2753–2765 (2009).

60. Filatov, E. et al. CCL22-expressing stem cell-derived islet grafts recruit regulatory T cells in mice. Transplantation (2025) doi:10.1097/TP.0000000000005463.

61. Vestergaard, C. et al. A Th2 chemokine, TARC, produced by keratinocytes may recruit CLA+CCR4+ lymphocytes into lesional atopic dermatitis skin. J. Invest. Dermatol. 115, 640–646 (2000).

62. Rapp, M. et al. CCL22 controls immunity by promoting regulatory T cell communication with dendritic cells in lymph nodes. J. Exp. Med. 216, 1170–1181 (2019).

63. Stutte, S. et al. Requirement of CCL17 for CCR7- and CXCR4-dependent migration of cutaneous dendritic cells. Proc. Natl. Acad. Sci. U. S. A. 107, 8736–8741 (2010).

64. Knipfer, L. et al. A CCL1/CCR8-dependent feed-forward mechanism drives ILC2 functions in type 2-mediated inflammation. J. Exp. Med. 216, 2763–2777 (2019).

65. Kersh, A. E., et al. CXCL9, CXCL10, and CCL19 synergistically recruit T lymphocytes to skin in lichen planus. JCI Insight 9, (2024).

66. Kohrgruber, N. et al. Plasmacytoid dendritic cell recruitment by immobilized CXCR3 ligands. J. Immunol. 173, 6592–6602 (2004).

67. Eidsmo, L. & Martini, E. Human Langerhans cells with pro-inflammatory features relocate within psoriasis lesions. Front. Immunol. 9, 300 (2018).

68. Tanei, R. & Hasegawa, Y. Immunohistopathological analysis of spongiosis formation in atopic dermatitis compared with other skin diseases. Dermatopathology (Basel) 12, 23 (2025).

69. Laky, K., et al. Epithelial-intrinsic defects in TGFβR signaling drive local allergic inflammation manifesting as eosinophilic esophagitis. Sci. Immunol. 8, eabp9940 (2023).

70. He, H. et al. Single-cell transcriptome analysis of human skin identifies novel fibroblast subpopulation and enrichment of immune subsets in atopic dermatitis. J. Allergy Clin. Immunol. 145, 1615–1628 (2020).

71. Fiskin, E., et al. Multi-modal skin atlas identifies a multicellular immune-stromal community associated with altered cornification and specific T cell expansion in atopic dermatitis. bioRxiv (2023) doi:10.1101/2023.10.29.563503.

72. Bangert, C., et al. Persistence of mature dendritic cells, TH2A, and Tc2 cells characterize clinically resolved atopic dermatitis under IL-4Rα blockade. Sci. Immunol. 6, eabe2749 (2021).

73. Zhang, B. et al. Single-cell profiles reveal distinctive immune response in atopic dermatitis in contrast to psoriasis. Allergy 78, 439–453 (2023).

74. Friedrich, M. et al. IL-1-driven stromal-neutrophil interactions define a subset of patients with inflammatory bowel disease that does not respond to therapies. Nat. Med. 27, 1970–1981 (2021).

75. Gao, Y. et al. Cross-tissue human fibroblast atlas reveals myofibroblast subtypes with distinct roles in immune modulation. Cancer Cell 0, (2024).

76. Augustin, M. et al. Unveiling the true costs and societal impacts of moderate-to-severe atopic dermatitis in Europe. J. Eur. Acad. Dermatol. Venereol. 36 **Suppl 7**, 3–16 (2022).

77. Adachi, T. et al. Hair follicle-derived IL-7 and IL-15 mediate skin-resident memory T cell homeostasis and lymphoma. Nat. Med. 21, 1272–1279 (2015).

78. Mackay, L. K. et al. The developmental pathway for CD103(+)CD8+ tissue-resident memory T cells of skin. Nat. Immunol. 14, 1294–1301 (2013).

79. Jabri, B. & Abadie, V. IL-15 functions as a danger signal to regulate tissue-resident T cells and tissue destruction. Nat. Rev. Immunol. 15, 771–783 (2015).

80. Qiu, Z. et al. Retinoic acid signaling during priming licenses intestinal CD103+ CD8 TRM cell differentiation. J. Exp. Med. 220, (2023).

81. Obers, A. et al. Retinoic acid and TGF-β orchestrate organ-specific programs of tissue residency. Immunity 57, 2615–2633.e10 (2024).

82. Pan, Y. et al. Survival of tissue-resident memory T cells requires exogenous lipid uptake and metabolism. Nature 543, 252–256 (2017).

83. Maden, M. Retinoic acid in the development, regeneration and maintenance of the nervous system. Nat. Rev. Neurosci. 8, 755–765 (2007).

84. Rhinn, M. & Dollé, P. Retinoic acid signalling during development. Development 139, 843–858 (2012).

85. Miranda, A. M. A. et al. Selective remodelling of the adipose niche in obesity and weight loss. Nature 644, 769–779 (2025).

86. Devi, K. S. P. et al. PD-1 is requisite for skin TRM cell formation and specification by TGFβ. Nat. Immunol. 26, 1339–1351 (2025).

87. Mani, V. et al. Migratory DCs activate TGF-β to precondition naïve CD8+ T cells for tissue-resident memory fate. Science 366, eaav5728 (2019).

88. Akkaya, M., Kwak, K. & Pierce, S. K. B cell memory: building two walls of protection against pathogens. Nat. Rev. Immunol. 20, 229–238 (2020).

89. Wilson, R. P. et al. IgM plasma cells reside in healthy skin and accumulate with chronic inflammation. J. Invest. Dermatol. 139, 2477–2487 (2019).

90. Gribonika, I. et al. Skin autonomous antibody production regulates host-microbiota interactions. Nature 638, 1043–1053 (2025).

91. Kamberov, Y. G. et al. A genetic basis of variation in eccrine sweat gland and hair follicle density. Proc. Natl. Acad. Sci. U. S. A. 112, 9932–9937 (2015).

92. Metze, D., Kersten, A., Jurecka, W. & Gebhart, W. Immunoglobulins coat microorganisms of skin surface: a comparative immunohistochemical and ultrastructural study of cutaneous and oral microbial symbionts. J. Invest. Dermatol. 96, 439–445 (1991).

93. Madissoon, E. et al. A spatially resolved atlas of the human lung characterizes a gland-associated immune niche. Nat. Genet. 55, 66–77 (2022).

94. Bunker, J. J. & Bendelac, A. IgA responses to Microbiota. Immunity 49, 211–224 (2018).

95. Macpherson, A. J., McCoy, K. D., Johansen, F.-E. & Brandtzaeg, P. The immune geography of IgA induction and function. Mucosal Immunol. 1, 11–22 (2008).

96. Hu, S., Yang, K., Yang, J., Li, M. & Xiong, N. Critical roles of chemokine receptor CCR10 in regulating memory IgA responses in intestines. Proc. Natl. Acad. Sci. U. S. A. 108, E1035–44 (2011).

97. Shang, L. et al. Toll-like receptor signaling in small intestinal epithelium promotes B-cell recruitment and IgA production in lamina propria. Gastroenterology 135, 529–538 (2008).

98. Samiea, A., Celis, G., Yadav, R., Rodda, L. B. & Moreau, J. M. B cells in non-lymphoid tissues. Nat. Rev. Immunol. (2025) doi:10.1038/s41577-025-01137-6.

99. Mitchell, B. L. et al. Genome-wide association meta-analysis identifies 29 new acne susceptibility loci. Nat. Commun. 13, 702 (2022).

100. Budu-Aggrey, A. et al. European and multi-ancestry genome-wide association meta-analysis of atopic dermatitis highlights importance of systemic immune regulation. Nat. Commun. 14, 6172 (2023).

101. Dand, N. et al. GWAS meta-analysis of psoriasis identifies new susceptibility alleles impacting disease mechanisms and therapeutic targets. Nat. Commun. 16, 2051 (2025).

102. Zhang, M. et al. Multi-ancestry genome-wide meta-analysis with 472,819 individuals identifies 32 novel risk loci for psoriasis. J. Transl. Med. 23, 133 (2025).

103. Ochoa, D. et al. The next-generation Open Targets Platform: reimagined, redesigned, rebuilt. Nucleic Acids Res. 51, D1353–D1359 (2023).

104. Ober-Reynolds, B. et al. Integrated single-cell chromatin and transcriptomic analyses of human scalp identify gene-regulatory programs and critical cell types for hair and skin diseases. Nat. Genet. 55, 1288–1300 (2023).

105. Lee, E. Y. et al. Functional interrogation of lymphocyte subsets in alopecia areata using single-cell RNA sequencing. Proc. Natl. Acad. Sci. U. S. A. 120, e2305764120 (2023).

106. Do, T. H., et al. TREM2 macrophages induced by human lipids drive inflammation in acne lesions. Sci. Immunol. 7, eabo2787 (2022).

107. Betz, R. C. et al. Genome-wide meta-analysis in alopecia areata resolves HLA associations and reveals two new susceptibility loci. Nat. Commun. 6, 5966 (2015).

108. Teder-Laving, M. et al. Genome-wide meta-analysis identifies novel loci conferring risk of acne vulgaris. Eur. J. Hum. Genet. 32, 1136–1143 (2024).

109. Kobayashi, T. et al. Homeostatic control of sebaceous glands by innate lymphoid cells regulates commensal bacteria equilibrium. Cell 176, 982–997.e16 (2019).

110. Choa, R. et al. Thymic stromal lymphopoietin induces adipose loss through sebum hypersecretion. Science 373, eabd2893 (2021).

111. Thiboutot, D. Regulation of human sebaceous glands. J. Invest. Dermatol. 123, 1–12 (2004).

112. Alexopoulos, A., Kakourou, T., Orfanou, I., Xaidara, A. & Chrousos, G. Retrospective analysis of the relationship between infantile seborrheic dermatitis and atopic dermatitis. Pediatr. Dermatol. 31, 125–130 (2014).

113. Silverberg, N. B. Typical and atypical clinical appearance of atopic dermatitis. Clin. Dermatol. 35, 354–359 (2017).

114. Gori, N., Ippoliti, E., Peris, K. & Chiricozzi, A. Head and neck atopic dermatitis: still a challenging manifestation in the biologic era. Expert Opin. Biol. Ther. 23, 575–577 (2023).

115. Bousbaine, D. et al. Discovery and engineering of the antibody response to a prominent skin commensal. Nature 1–3 (2024).

116. Bemark, M. et al. Limited clonal relatedness between gut IgA plasma cells and memory B cells after oral immunization. Nat. Commun. 7, 12698 (2016).

117. Bartnikas, L. M. et al. Epicutaneous sensitization results in IgE-dependent intestinal mast cell expansion and food-induced anaphylaxis. J. Allergy Clin. Immunol. 131, 451–60.e1–6 (2013).

118. Francis, L. et al. Single-cell analysis of psoriasis resolution demonstrates an inflammatory fibroblast state targeted by IL-23 blockade. Nat. Commun. 15, 913 (2024).

119. Castillo, R. L., et al. Spatial transcriptomics stratifies psoriatic disease severity by emergent cellular ecosystems. Sci. Immunol. 8, eabq7991 (2023).

120. Mitamura, Y. et al. Spatial transcriptomics combined with single-cell RNA-sequencing unravels the complex inflammatory cell network in atopic dermatitis. Allergy 78, 2215–2231 (2023).

121. Schäbitz, A. et al. Spatial transcriptomics landscape of lesions from non-communicable inflammatory skin diseases. Nat. Commun. 13, 7729 (2022).

122. He, H. et al. Mild atopic dermatitis lacks systemic inflammation and shows reduced nonlesional skin abnormalities. J. Allergy Clin. Immunol. 147, 1369–1380 (2021).

123. Hughes, T. K. et al. Second-Strand Synthesis-based massively parallel scRNA-seq reveals cellular states and molecular features of human inflammatory skin pathologies. Immunity 53, 878–894.e7 (2020).

124. Yu, Z. et al. Genetic variation across and within individuals. Nat. Rev. Genet. 25, 548–562 (2024).

125. Ding, J. et al. An esophagus cell atlas reveals dynamic rewiring during active eosinophilic esophagitis and remission. Nat. Commun. 15, 3344 (2024).

126. Hanifin, J. M. et al. The eczema area and severity index (EASI): assessment of reliability in atopic dermatitis: EASI: assessment of reliability in AD. Exp. Dermatol. 10, 11–18 (2001).

127. Ganier, C. et al. Multiscale spatial mapping of cell populations across anatomical sites in healthy human skin and basal cell carcinoma. Proc. Natl. Acad. Sci. U. S. A. 121, e2313326120 (2024).

128. Fleming, S. J. et al. Unsupervised removal of systematic background noise from droplet-based single-cell experiments using CellBender. Nat. Methods 20, 1323–1335 (2023).

129. Wolock, S. L., Lopez, R. & Klein, A. M. Scrublet: Computational identification of cell Doublets in Single-cell transcriptomic data. Cell Syst. 8, 281–291.e9 (2019).

130. Ahlmann-Eltze, C. & Huber, W. Comparison of transformations for single-cell RNA-seq data. Nat. Methods 20, 665–672 (2023).

131. Heumos, L., et al. Pertpy: an end-to-end framework for perturbation analysis. *bioRxiv* (2024) doi:10.1101/2024.08.04.606516.

132. Muzellec, B., Teleńczuk, M., Cabeli, V. & Andreux, M. PyDESeq2: a python package for bulk RNA-seq differential expression analysis. Bioinformatics 39, (2023).

133. Palla, G. et al. Squidpy: a scalable framework for spatial omics analysis. Nat. Methods 19, 171–178 (2022).

134. Wolf, F. A. et al. PAGA: graph abstraction reconciles clustering with trajectory inference through a topology preserving map of single cells. Genome Biol. 20, 59 (2019).

135. Birk, S. et al. Large-scale characterization of cell niches in spatial atlases using bio-inspired graph learning. bioRxiv 2024.02.21.581428 (2024) doi:10.1101/2024.02.21.581428.

136. Troulé, K. et al. CellPhoneDB v5: inferring cell-cell communication from single-cell multiomics data. arXiv [q-bio.CB] (2023).

137. Fang, Z., Liu, X. & Peltz, G. GSEApy: a comprehensive package for performing gene set enrichment analysis in Python. Bioinformatics 39, btac757 (2023).

138. Assran, M. et al. Self-supervised learning from images with a Joint-Embedding Predictive Architecture. arXiv [cs.CV] (2023).

139. Pirazzi, C. et al. PNPLA3 has retinyl-palmitate lipase activity in human hepatic stellate cells. Hum. Mol. Genet. 23, 4077–4085 (2014).

140. Schmidt, M., Binder, H. & Schneider, M. R. The metabolic underpinnings of sebaceous lipogenesis. *Commun*. Biol. 8, 670 (2025).

141. Kleshchevnikov, V. et al. Cell2location maps fine-grained cell types in spatial transcriptomics. Nat. Biotechnol. 40, 661–671 (2022).

142. Mall, M. A. et al. Cystic fibrosis. Nat. Rev. Dis. Primers 10, 53 (2024).

