## Supplementary Data Fig for "Hidden immune memory niches in inflammatory skin diseases"

Supplementary Fig. 1: 107 skin cell types

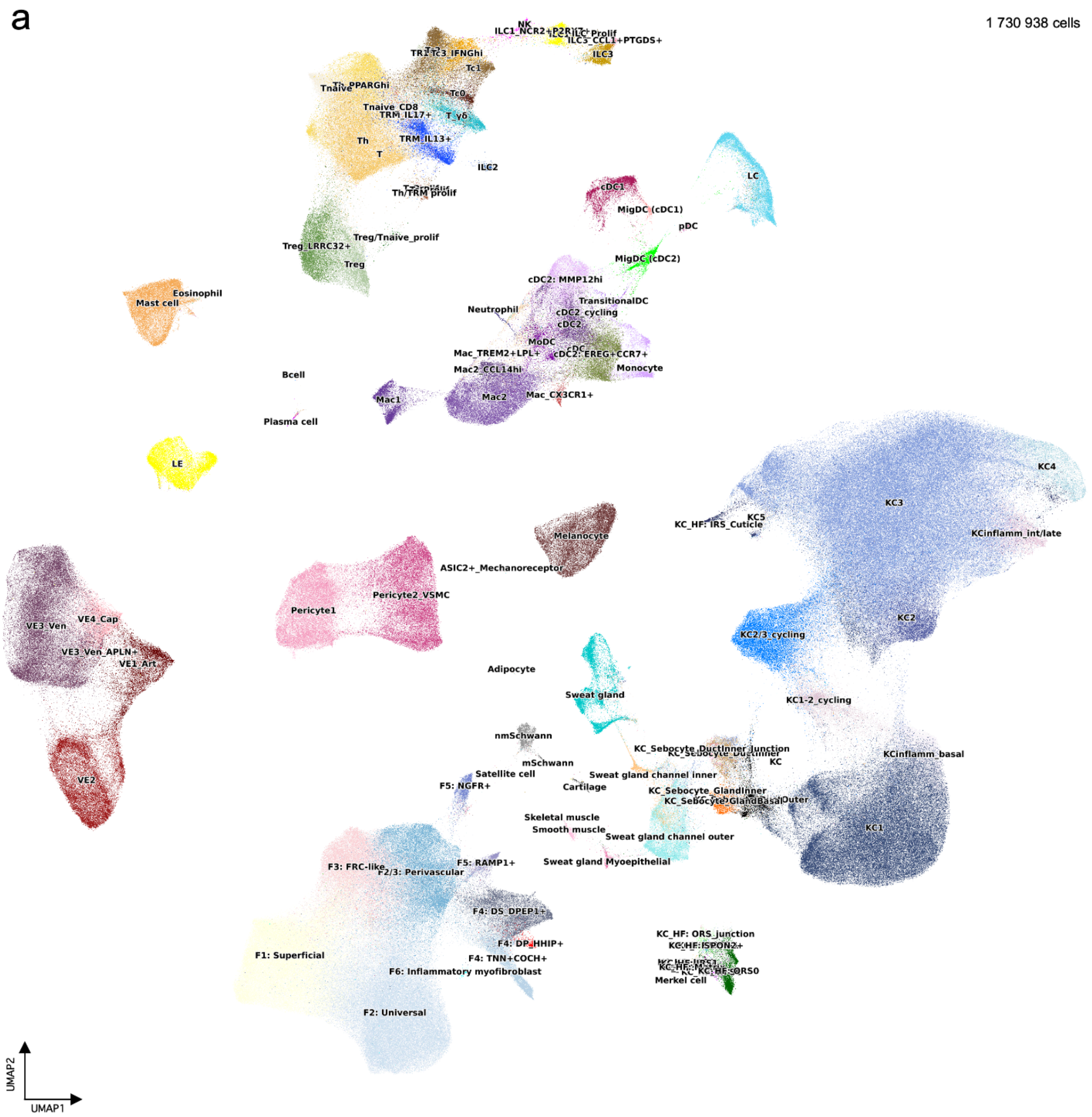


**Supplementary Figure 1 | 107 skin cell types. a,** UMAP visualisation of scRNA-seq dataset (lesional atopic dermatitis, lesional psoriasis, and non-lesional).

Supplementary Fig. 2: T cell populations (excluding ILC)

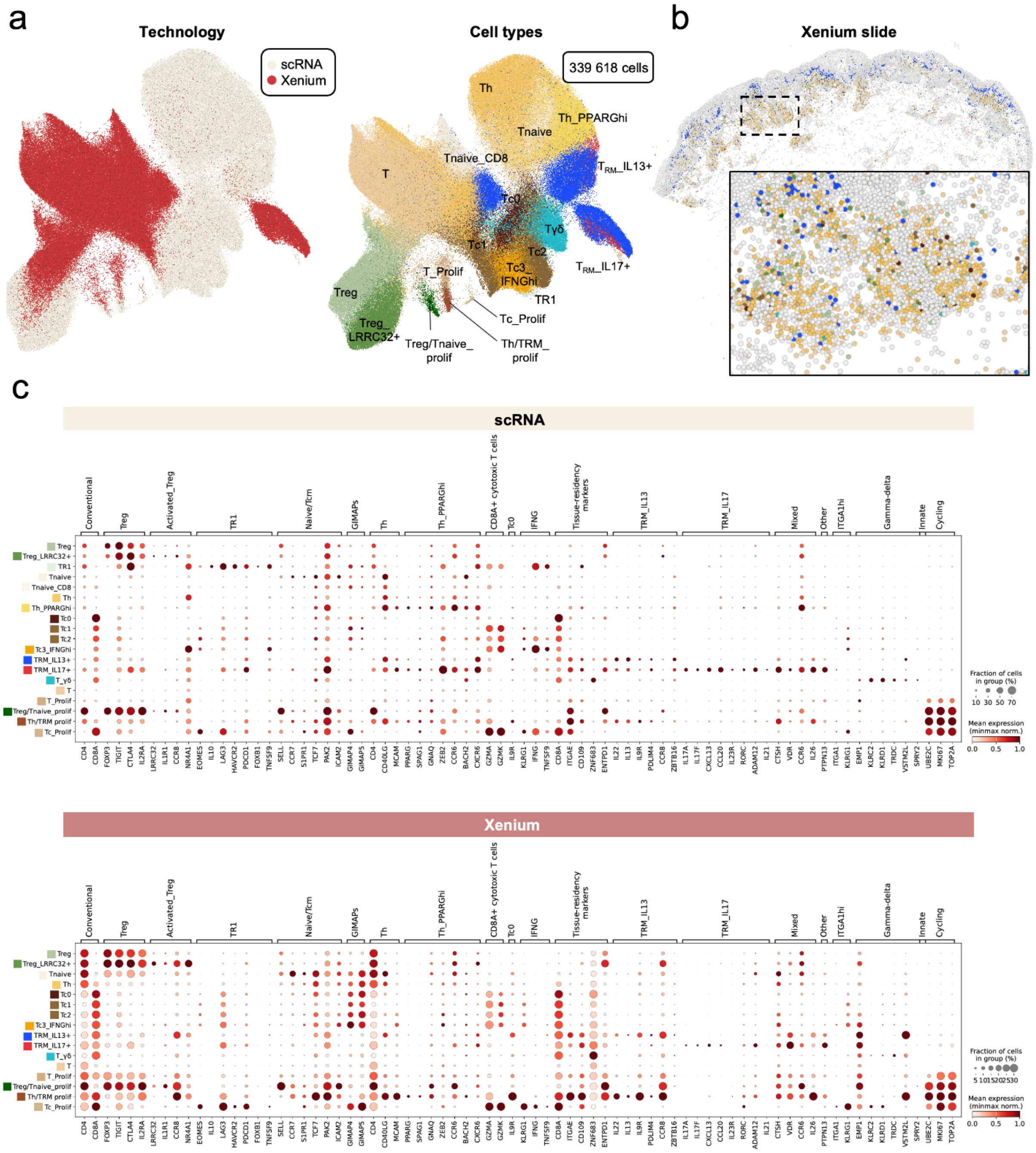


**Supplementary Figure 2 | T subsets (excluding innate lymphoid cells). a,** UMAP visualisation of cells by technology (left) and cell type (right). **b,** Xenium slide section (top right) coloured by cell subsets shown in panel a. **c,** Dot plot of same marker genes in scRNA-seq and Xenium 5k data.

Supplementary Fig. 3: Innate lymphoid cell (ILC) populations

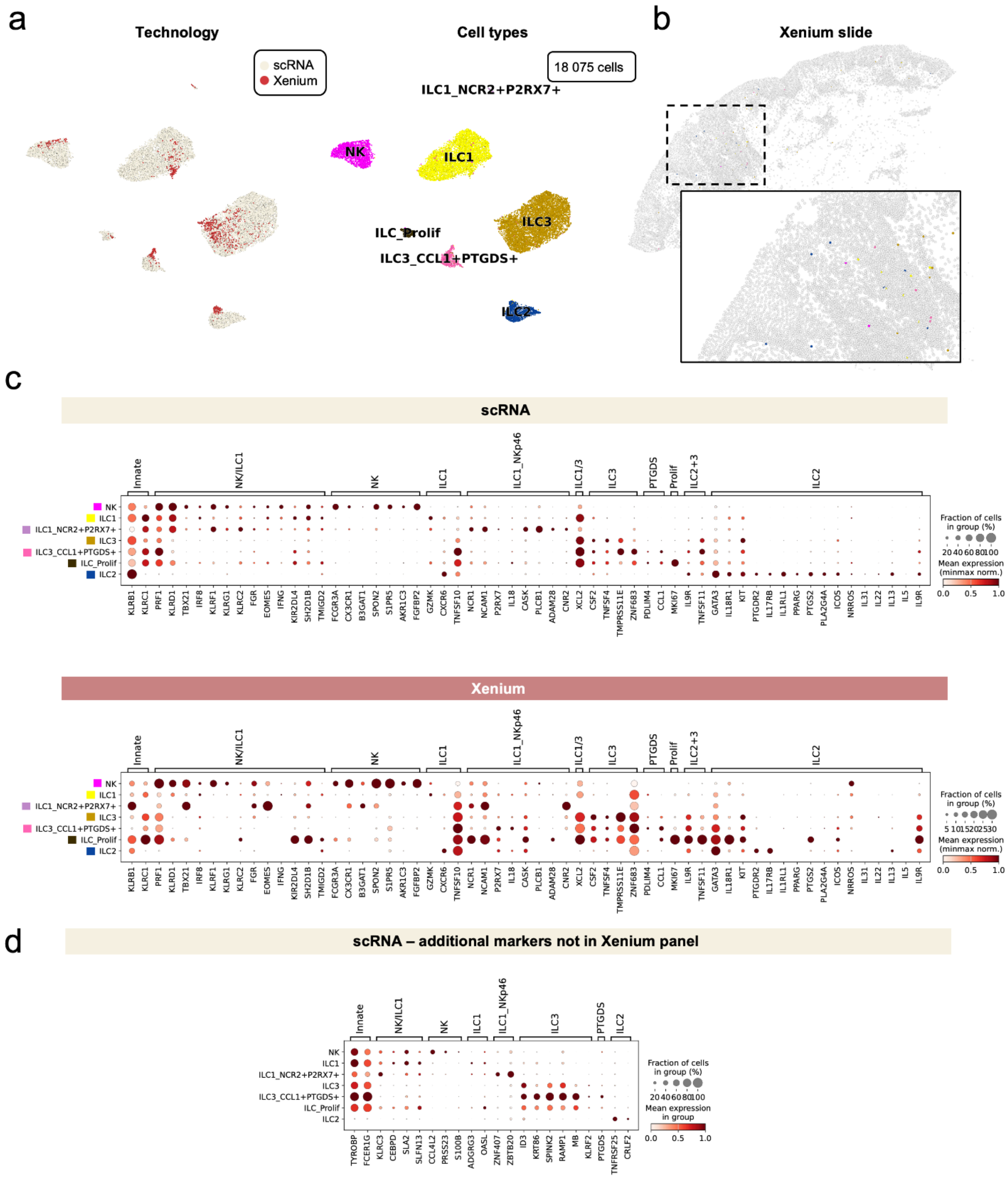


**Supplementary Figure 3 | Innate lymphoid cell (ILC) subsets. a,** UMAP visualisation of cells by technology (left) and cell type (right). **b,** Xenium slide section (top right) coloured by cell subsets shown in panel a. **c,** Dot plot of same marker genes in scRNA-seq and Xenium 5k data. **d,** Dot plot of marker genes only present in scRNA-seq data (not present in Xenium 5k panel).

Supplementary Fig. 4: Keratinocytes

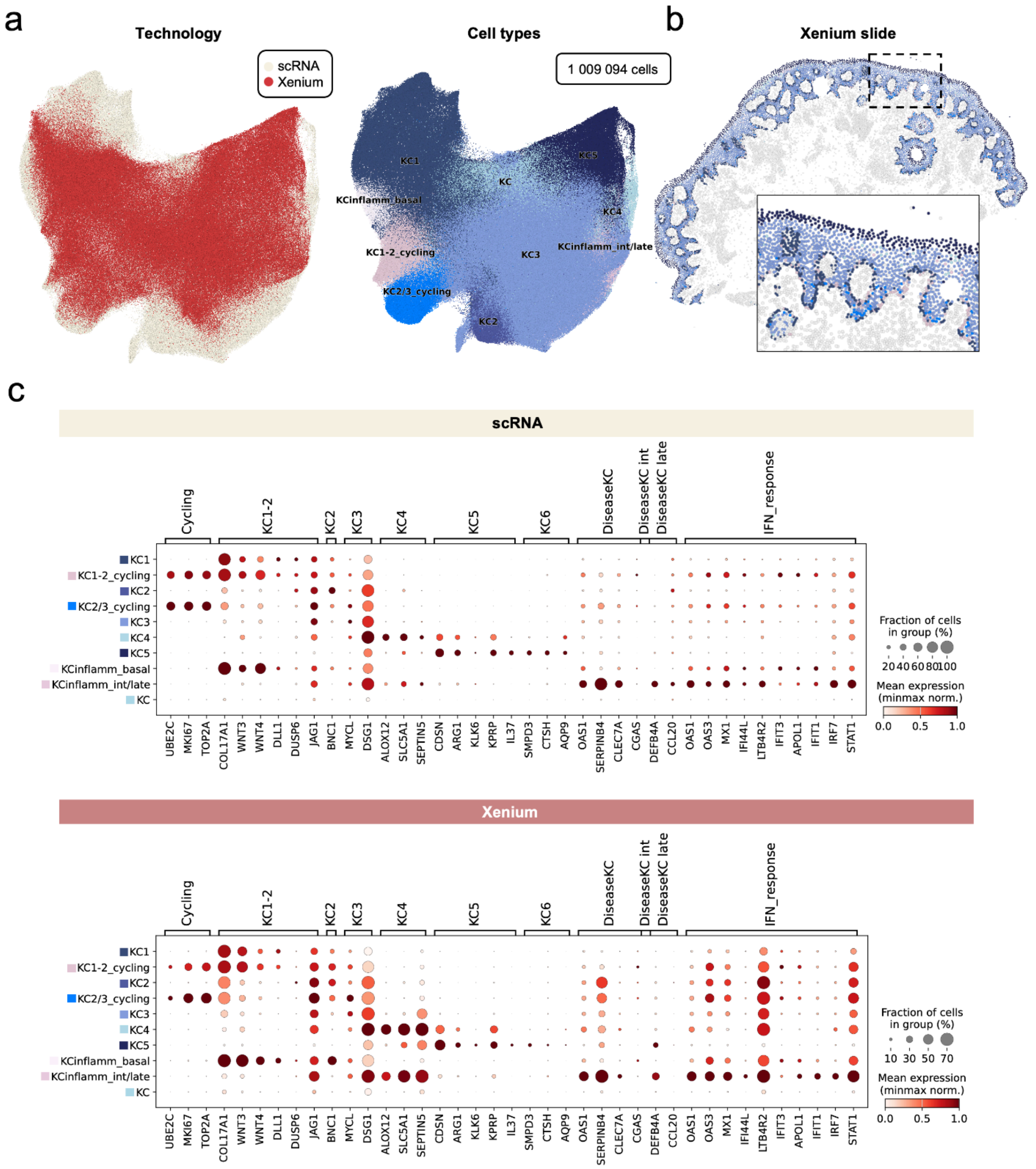


**Supplementary Figure 4 | Keratinocyte subsets. a,** UMAP visualisation of cells by technology (left) and cell type (right). **b,** Xenium slide section (top right) coloured by cell subsets shown in panel a. **c,** Dot plot of same marker genes in scRNA-seq and Xenium 5k data.

Supplementary Fig. 5: Fibroblasts

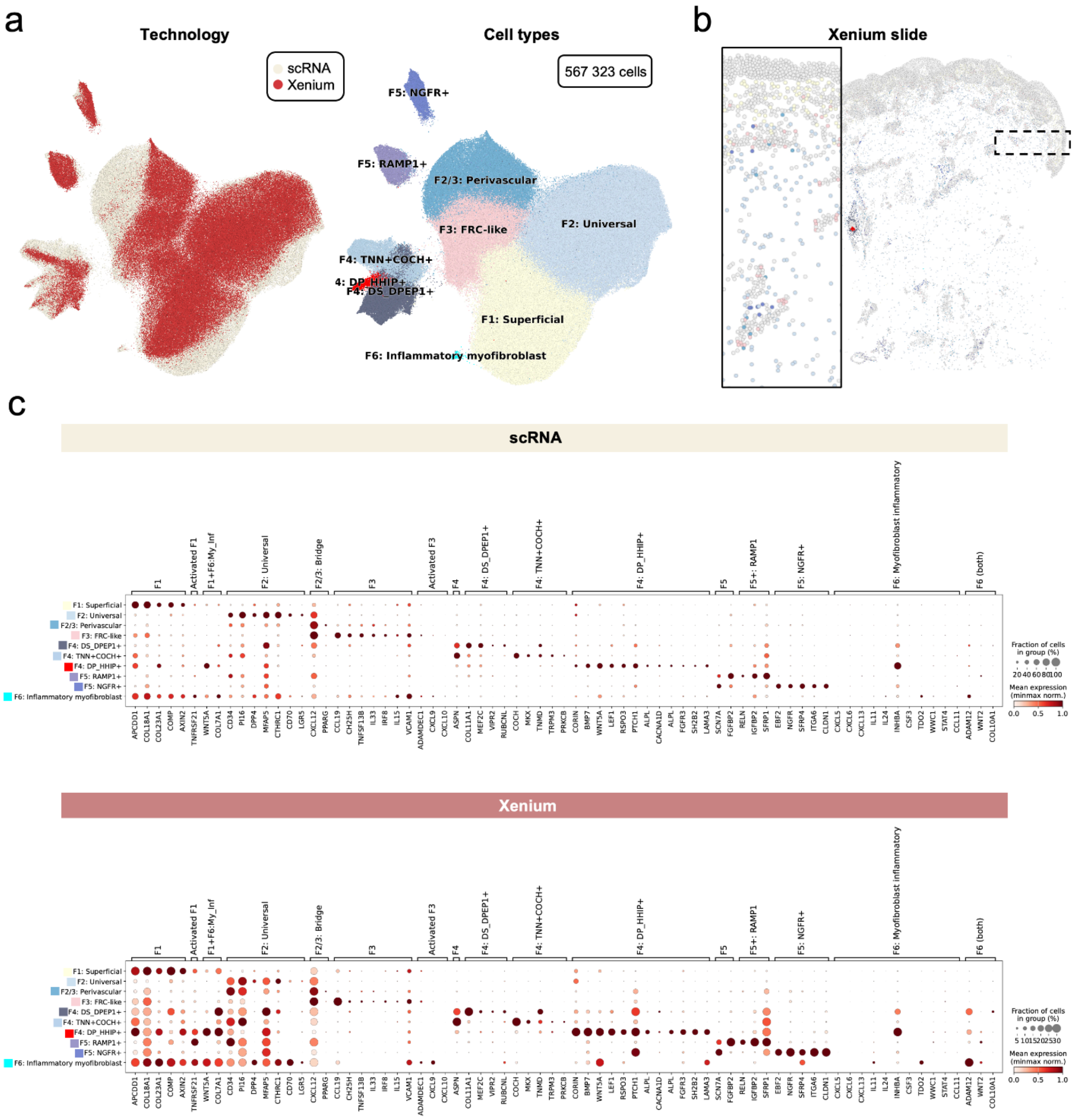


**Supplementary Figure 5 | Fibroblast subsets. a,** UMAP visualisation of cells by technology (left) and cell type (right). **b,** Xenium slide section (top right) coloured by cell subsets shown in panel a. **c,** Dot plot of same marker genes in scRNA-seq and Xenium 5k data.

Supplementary Fig. 6: Endothelium + pericyte + muscle populations

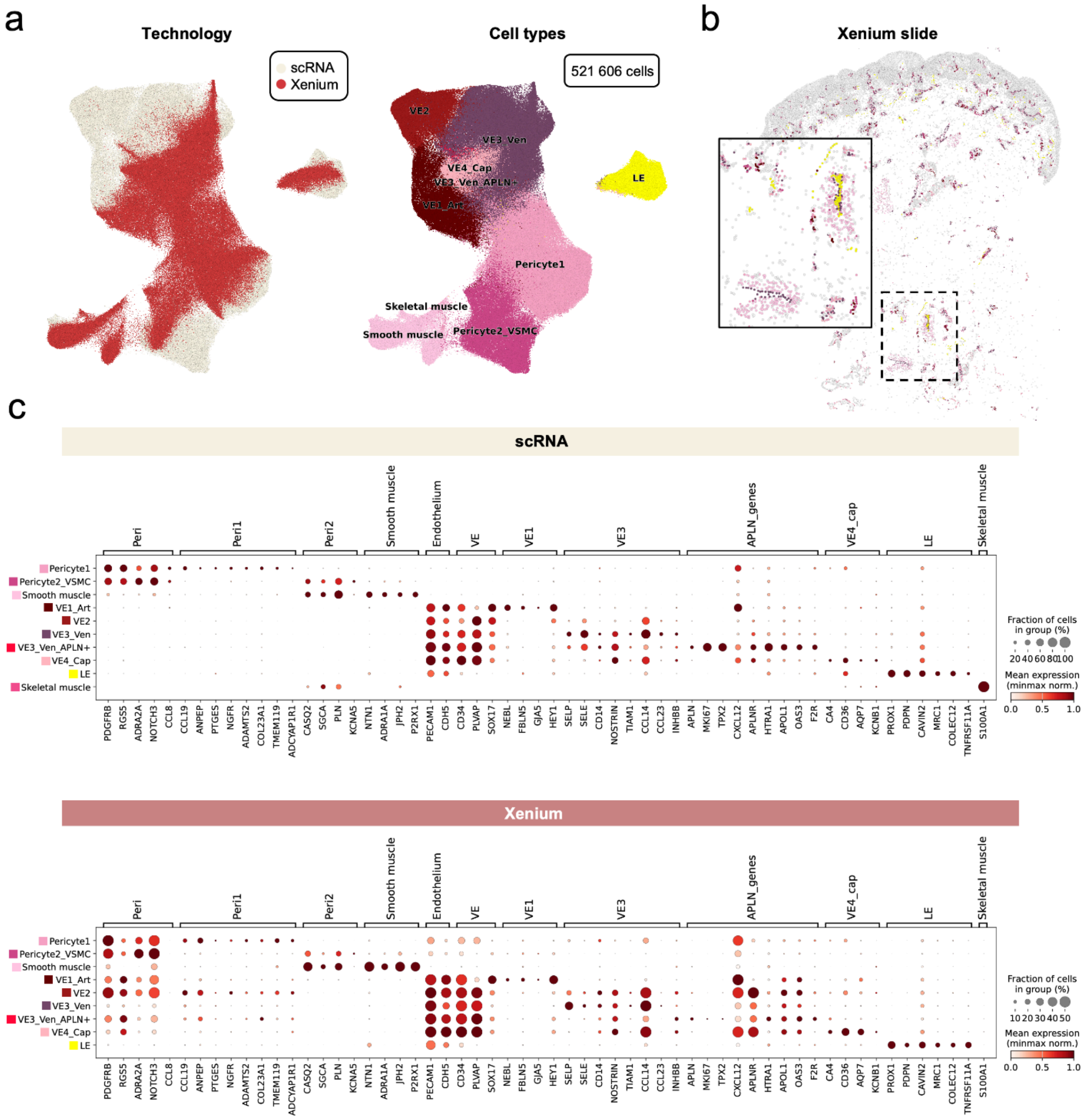


**Supplementary Figure 6 | Endothelium/pericyte/muscle subsets. a,** UMAP visualisation of cells by technology (left) and cell type (right). **b,** Xenium slide section (top right) coloured by cell subsets shown in panel a. **c,** Dot plot of same marker genes in scRNA-seq and Xenium 5k data.

Supplementary Fig. 7: Hair follicle

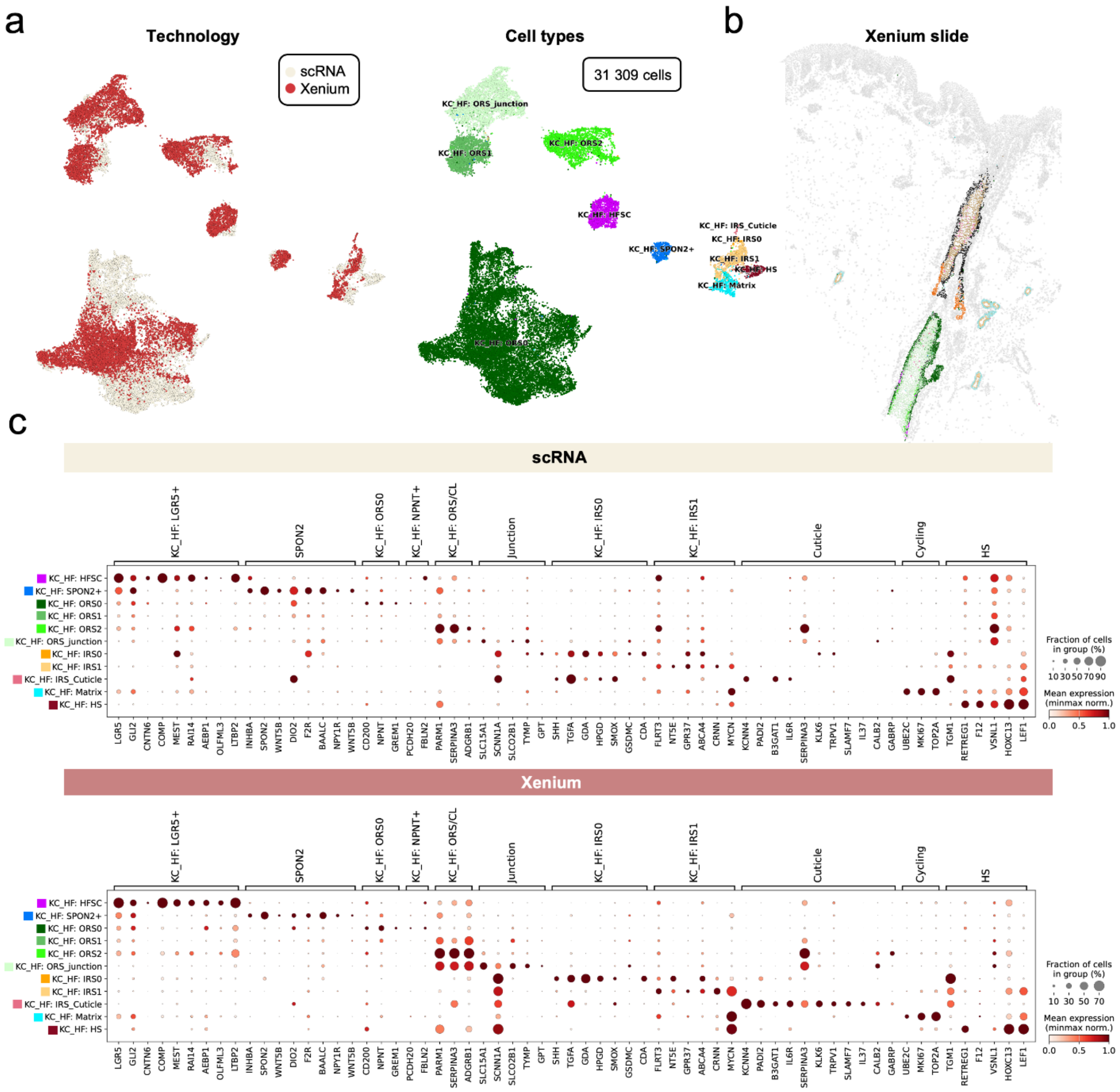


**Supplementary Figure 7 | Hair follicle subsets. a,** UMAP visualisation of cells by technology (left) and cell type (right). **b,** Xenium slide section (top right) coloured by cell subsets shown in panel a. **c,** Dot plot of same marker genes in scRNA-seq and Xenium 5k data.

Supplementary Fig. 8: Sebaceous gland

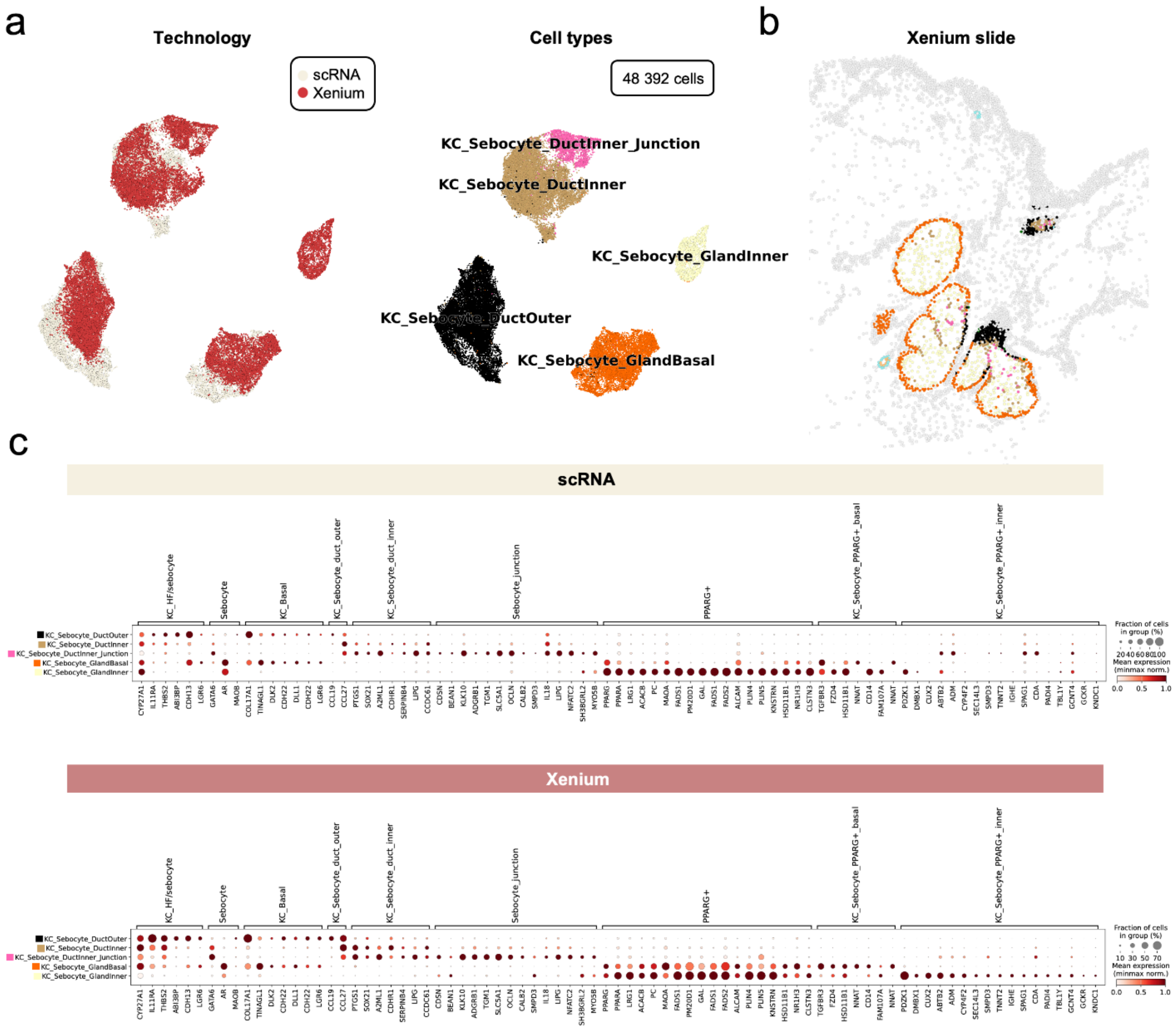


**Supplementary Figure 8 | Sebaceous subsets. a,** UMAP visualisation of cells by technology (left) and cell type (right). **b,** Xenium slide section (top right) coloured by cell subsets shown in panel a. **c,** Dot plot of same marker genes in scRNA-seq and Xenium 5k data.

Supplementary Fig. 9: Sweat gland

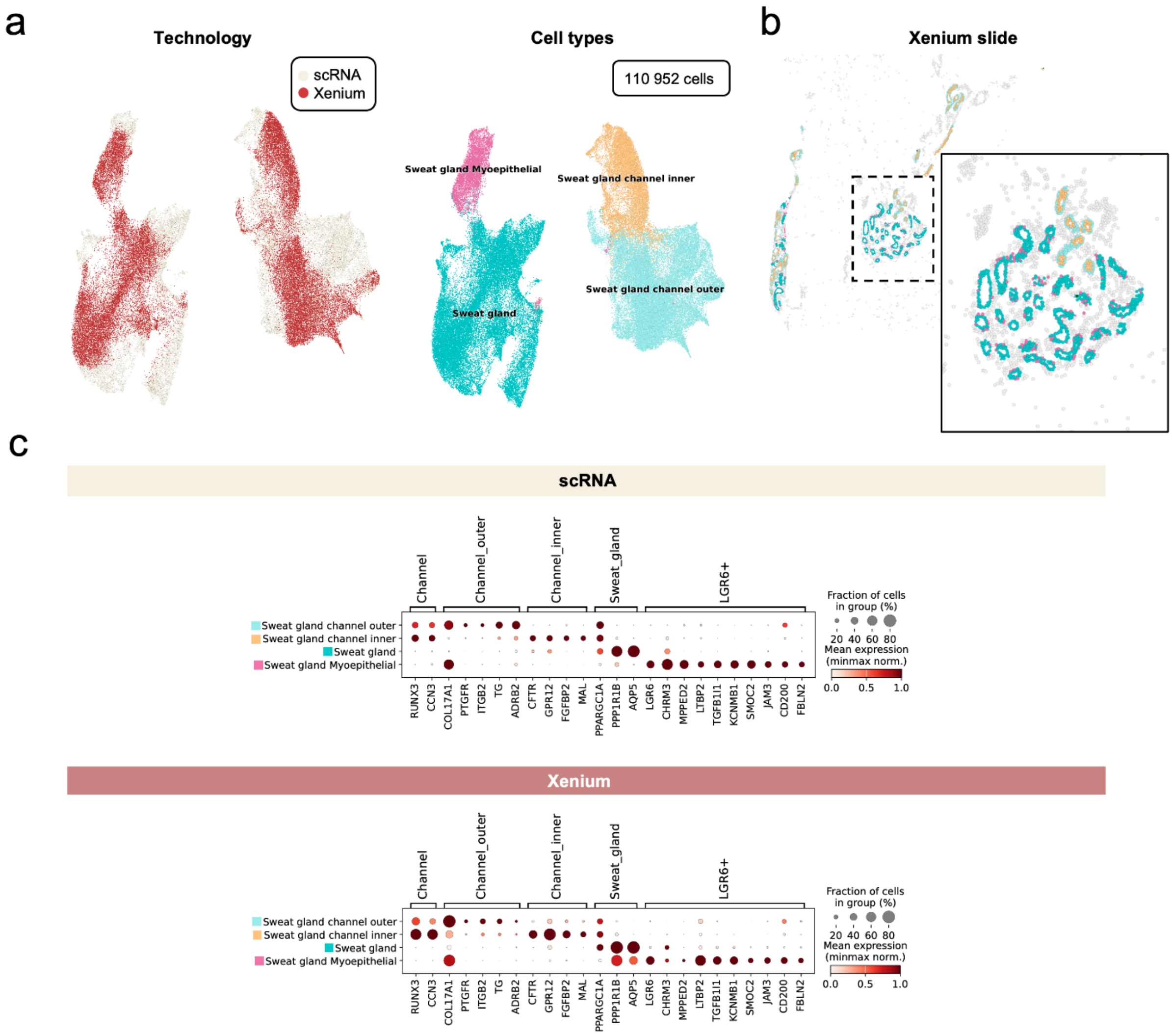


**Supplementary Figure 9 | Sweat gland subsets. a,** UMAP visualisation of cells by technology (left) and cell type (right). **b,** Xenium slide section (top right) coloured by cell subsets shown in panel a. **c,** Dot plot of same marker genes in scRNA-seq and Xenium 5k data.

Supplementary Fig. 10: Myeloid

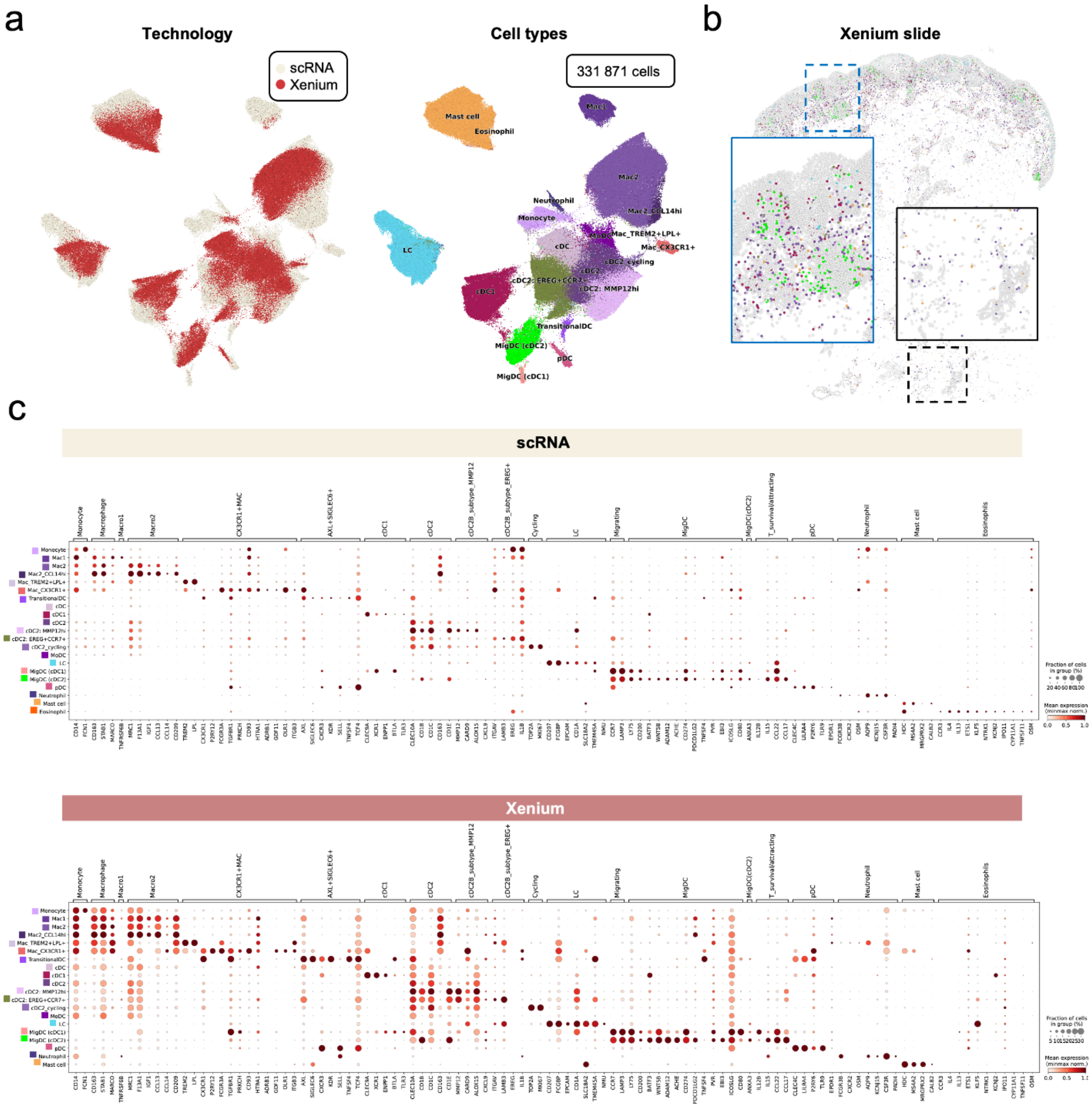


**Supplementary Figure 10 | Myeloid subsets. a,** UMAP visualisation of cells by technology (left) and cell type (right). **b,** Xenium slide section (top right) coloured by cell subsets shown in panel a. **c,** Dot plot of same marker genes in scRNA-seq and Xenium 5k data.

Supplementary Fig. 11: Schwann and other populations

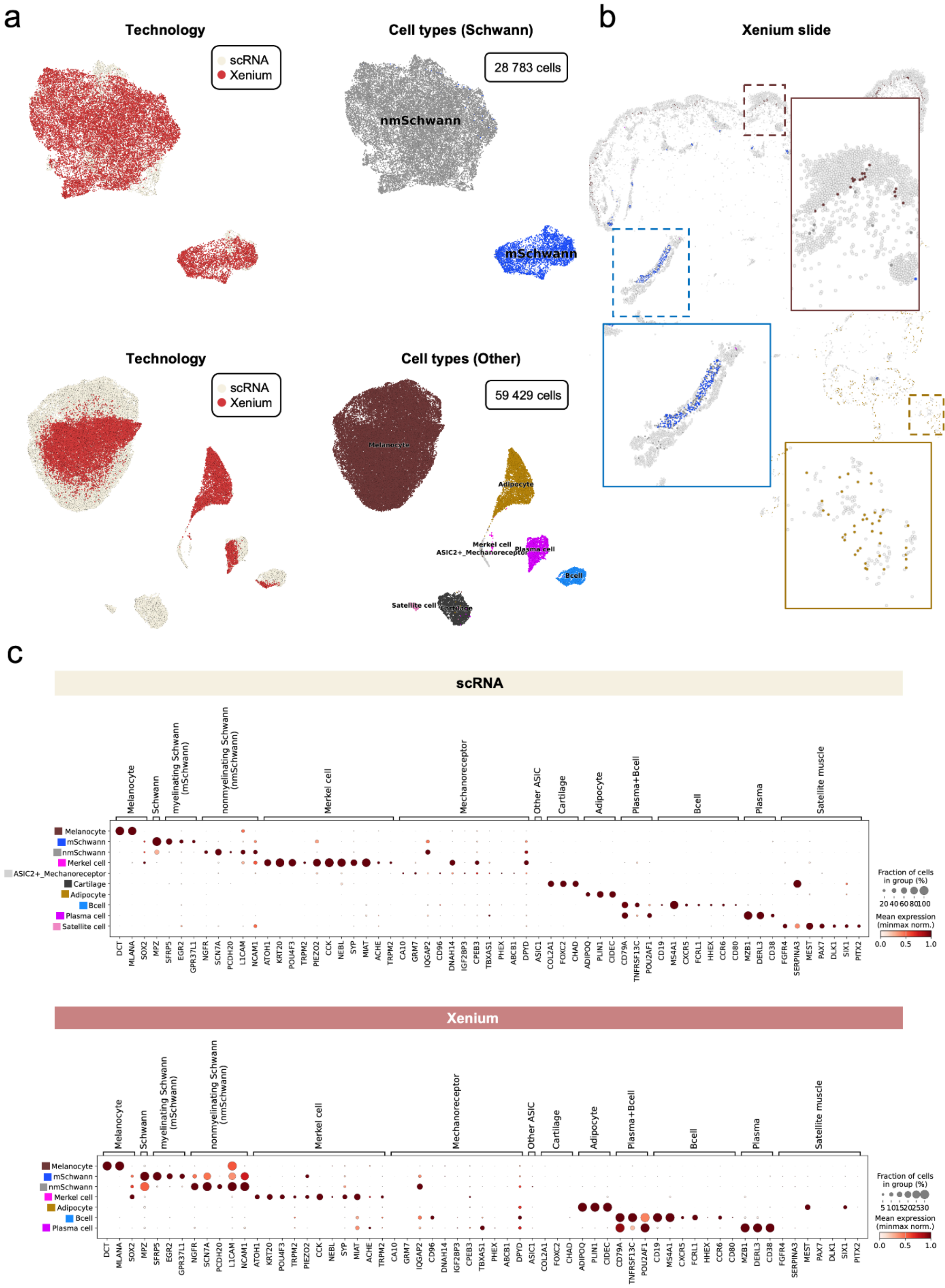


**Supplementary Figure 11 | Other subsets. a,** UMAP visualisation of cells by technology (left) and cell type (right). **b,** Xenium slide section (top right) coloured by cell subsets shown in panel a. **c,** Dot plot of same marker genes in scRNA-seq and Xenium 5k data.

Supplementary Fig. 12: Milo results and epidermal interaction maps.

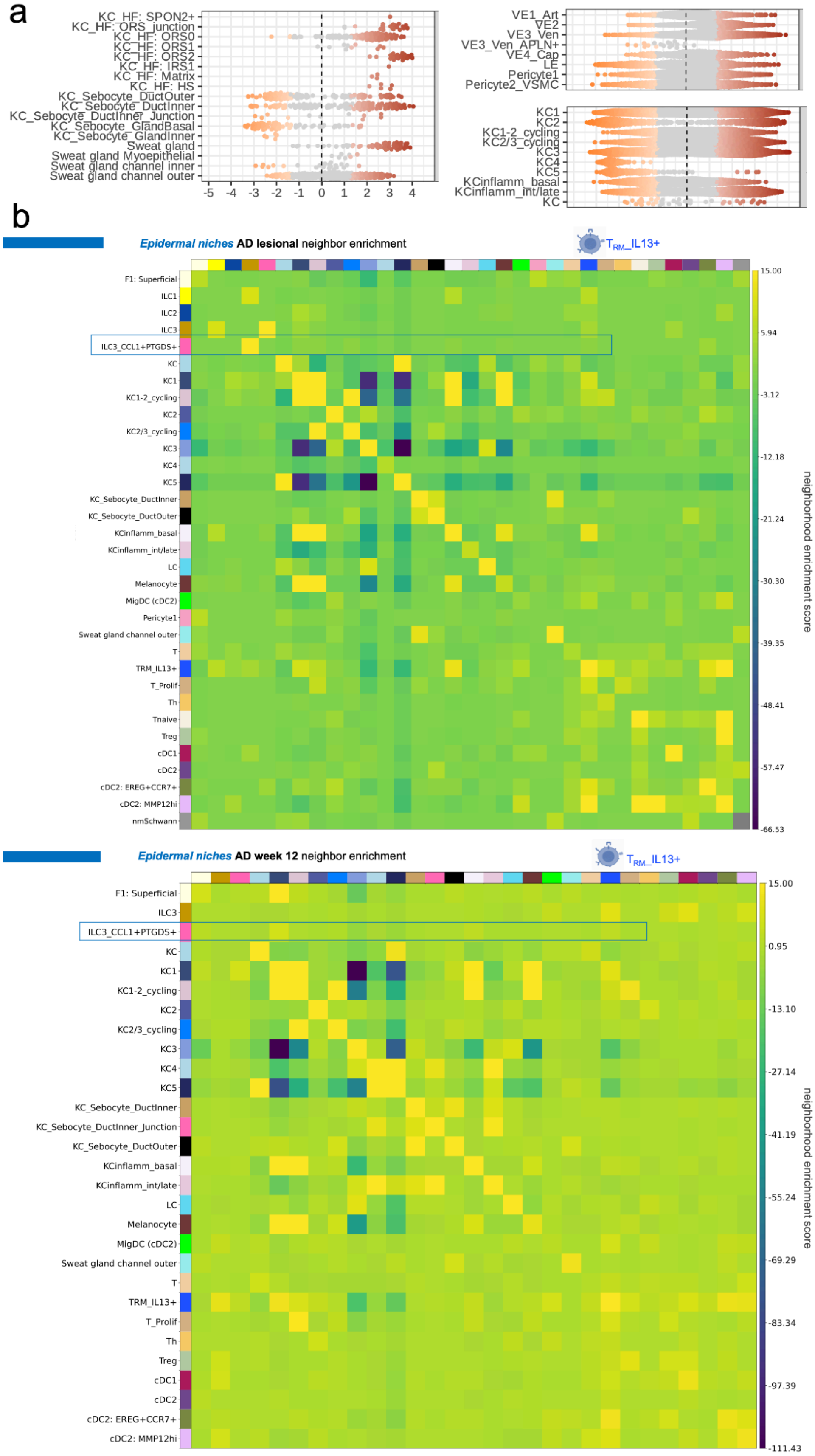


**Supplementary Fig. 12 | Milo results and epidermal interaction maps. a,** Differential abundance results from Milo for lesional AD skin vs lesional psoriasis skin for populations not shown in Fig. 1f. Coloured regions represent significant differences (false discovery rate 0.1). **b,** Neighbourhood enrichment plot for epidermis populations (defined from epidermal niches) in lesional and post-treatment AD skin. Red boxes indicate *ILC3_CCL1+PTGDS+* cells, with a relative enrichment with *TRM_IL13+* cells. AD: atopic dermatitis.

**Supplementary Fig. 13: Identification of cell types in external datasets**

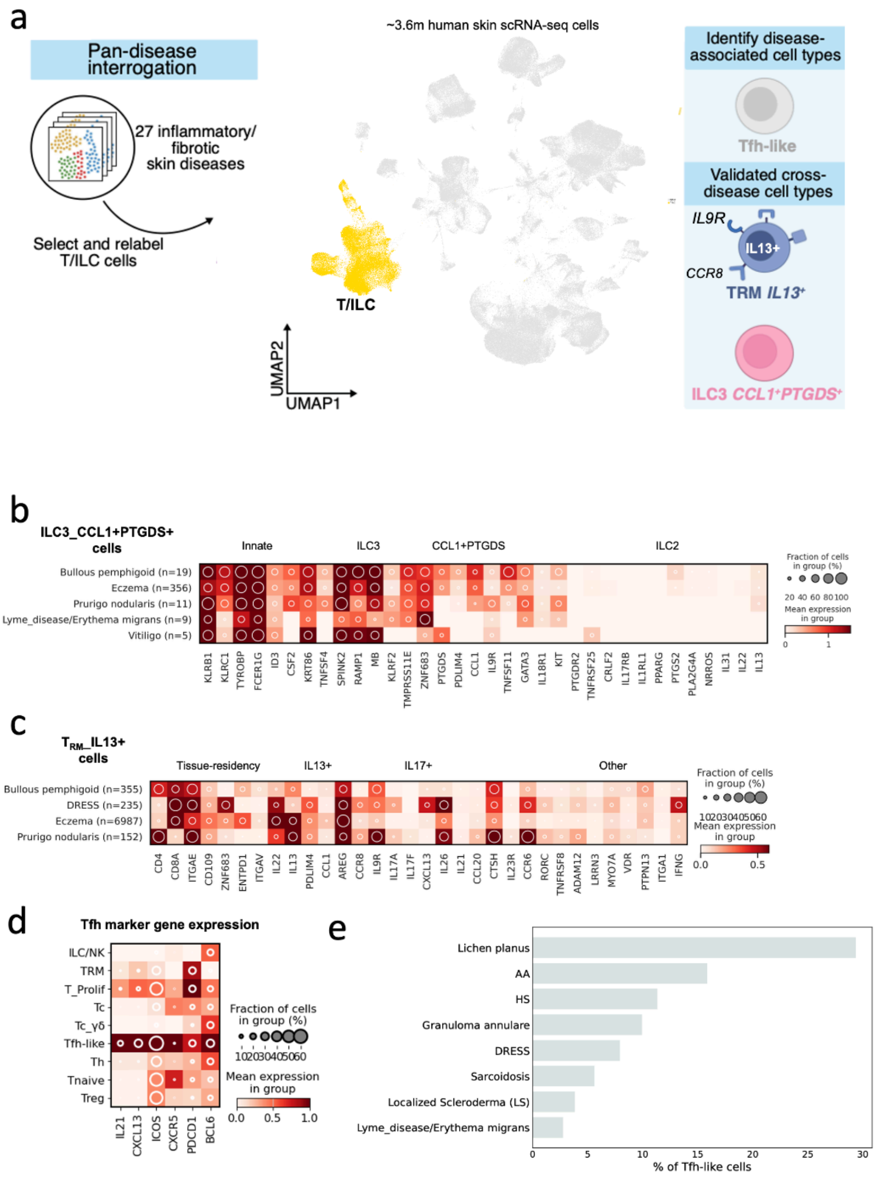


**Supplementary Fig. 13 | Identification of cell types in external datasets. a,** Schematic of methodology for skin T cell atlas generation (see Methods) and summary of novel cell states. **b,** Dotplot-heatmap combined of marker genes for *ILC3_CC1+PTGDS+* cells by disease. **c,** Dotplot-heatmap combined of marker genes for $T_{RM}$ cells by disease. **d,** Dotplot of Tfh marker genes. **e,** Prevalence of Tfh-like cells by patient status. We observed high prevalence in diseases with known bona fide tertiary lymphoid structures (hidradenitis suppurativa), as well as additional diseases such as lichen planus. Tfh: T follicular helper

Supplementary Fig. 14: Spatial cell distribution in non-lesional skin.

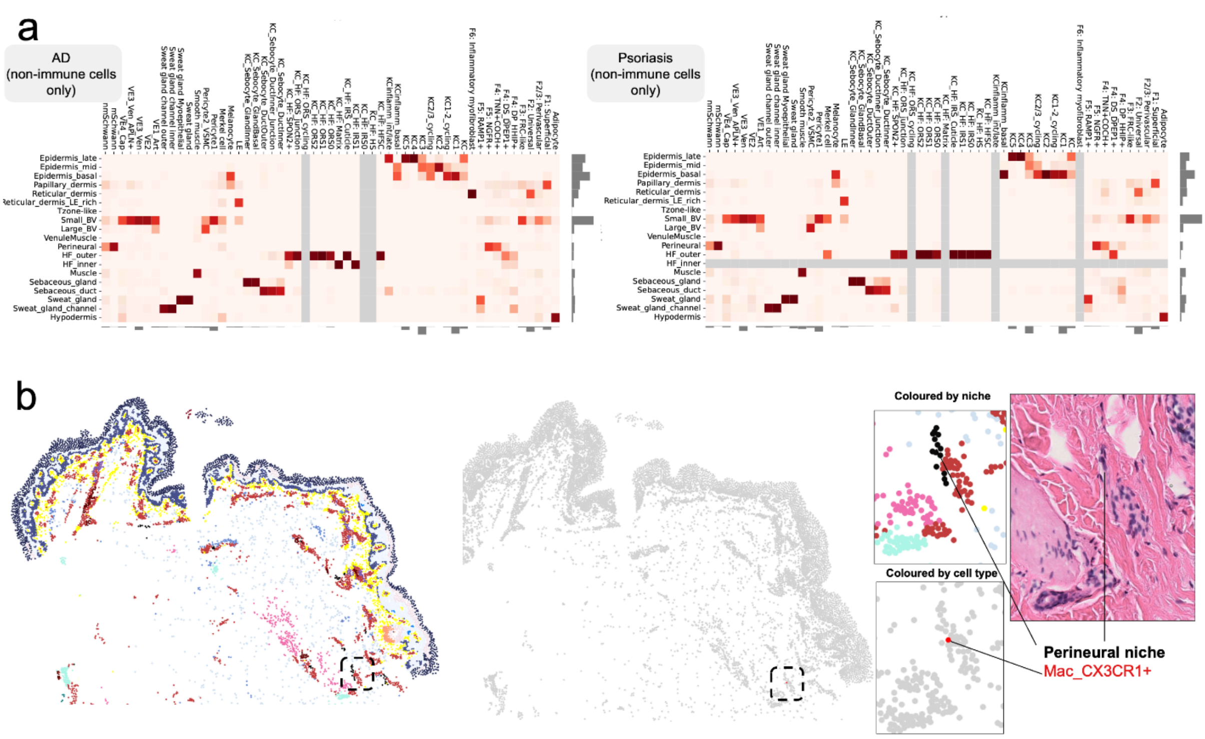


**Supplementary Fig. 14 | Spatial cell distribution in non-lesional skin. a,** Heatmap showing cell type distribution for non-immune cells by niche for non-lesional AD and psoriasis skin. **b,** Xenium 5k section of non-lesional skin coloured by niche (left) and cell type (other vs Mac_CX3CR1+). We identified the rare *Mac_CX3CR1+* population in the perineural niche, consistent with a neural region on H&E.

Supplementary Fig. 15: VisUtils predictions of cell types in non-lesional and healthy skin

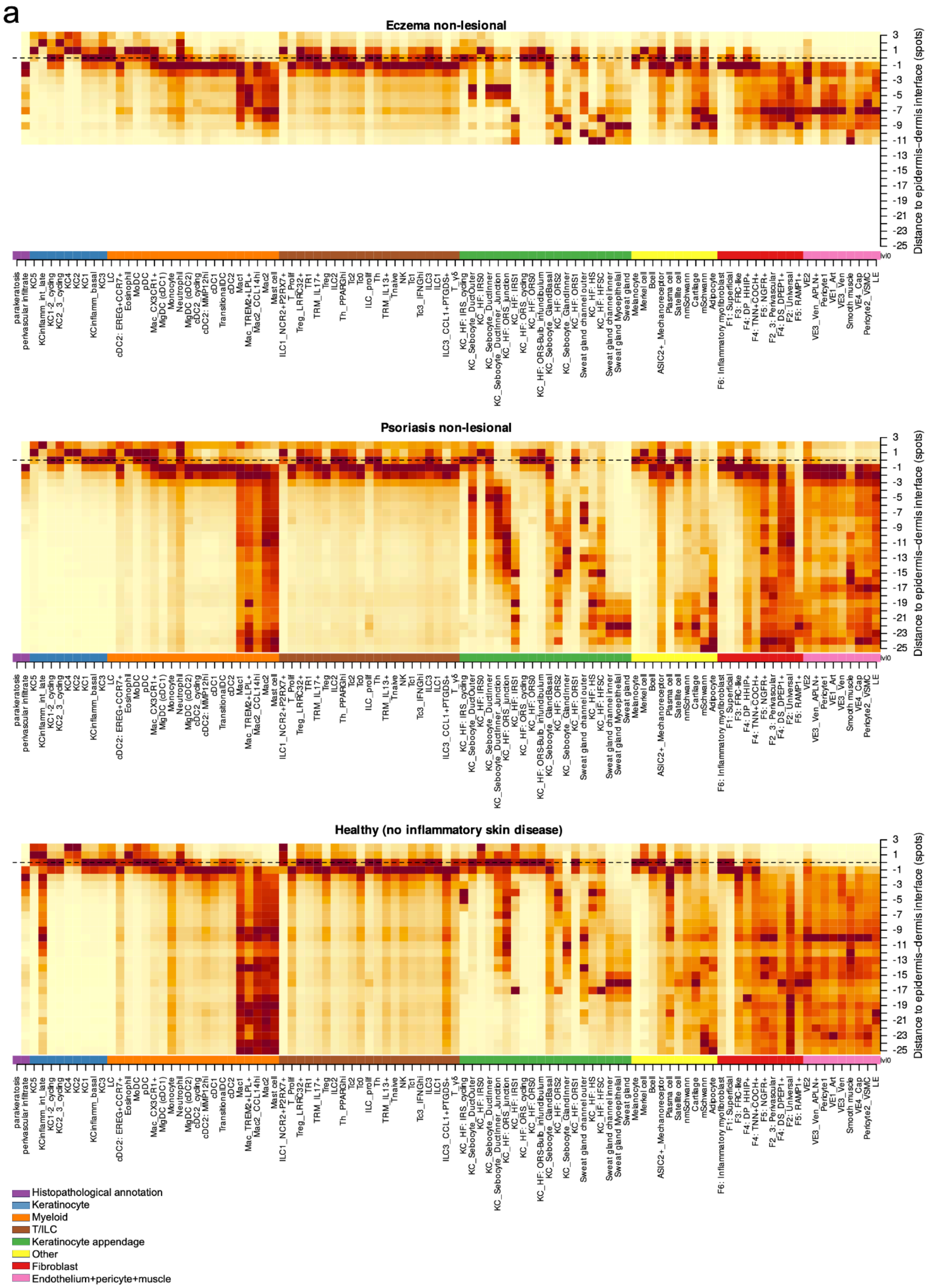


**Supplementary Figure 15 | VisUtils predictions of cell types in non-lesional and healthy skin. a,** Heatmap of cell abundance predictions by cell depth utilizing VisUtils in

non-lesional (atopic dermatitis (Eczema) and psoriasis) and healthy skin. Parakeratosis and perivascular infiltrate represent histopathological annotations of Visium spots.

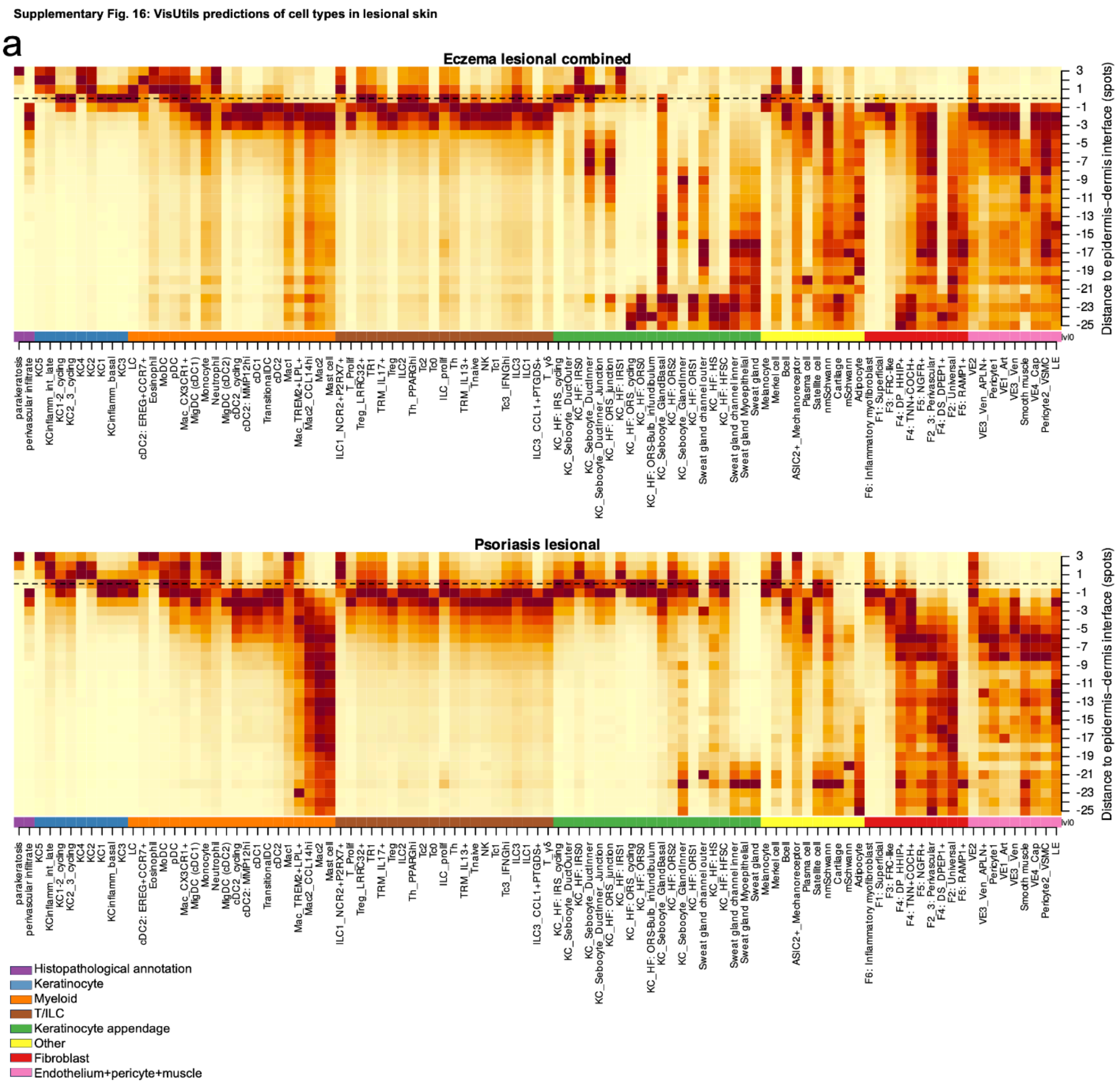


**Supplementary Figure 16 | VisUtils predictions of cell types in lesional skin. a,** Heatmap of cell abundance predictions by cell depth utilizing VisUtils in lesional skin (atopic dermatitis (Eczema) and psoriasis). Parakeratosis and perivascular infiltrate represent histopathological annotations of Visium spots.

Supplementary Fig. 17: Tzone-like niche and association with blood vessel network.

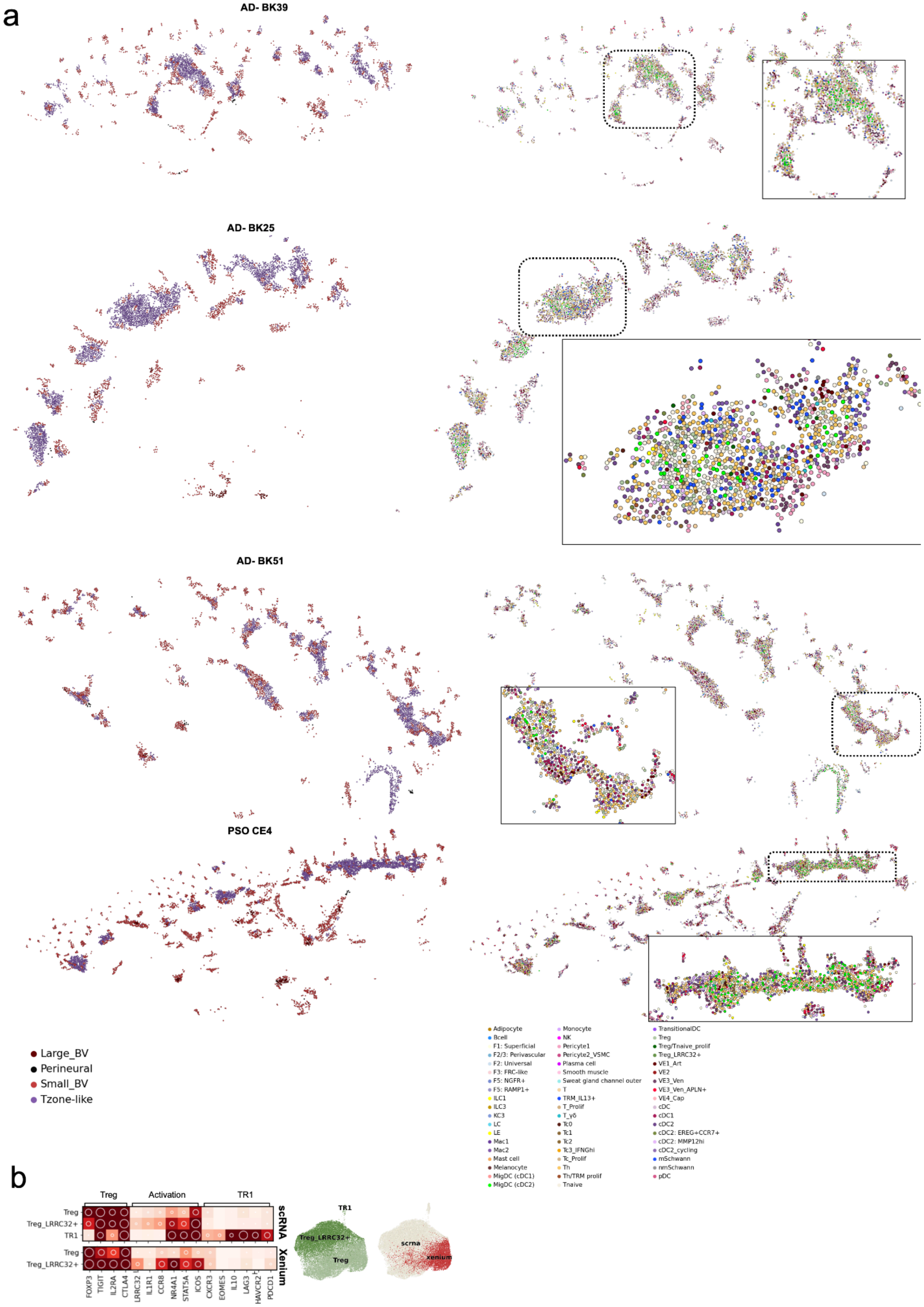


**Supplementary Fig. 17 | Tzone-like niche and association with blood vessel network.**
**a,** Xenium 5k tissue sections showing cells belonging to the *Tzone-like*, *Small_BV*,

*Large_BV*, or *Perineural* niches, coloured by niche (left column). The *Tzone-like* was typically identified next to the superficial vascular network, consistent with a site of extravasation of immune cells from blood. Same tissue sections coloured by cell type (right column). **b,** UMAP visualisation of Treg subsets for joint integration of scRNA-seq and Xenium data and dotplot-heatmap combined of Treg subtype markers in scRNA-seq and Xenium data from atlas.

**Supplementary Fig. 18: Plasma_cell_rich niche supporting genes.**

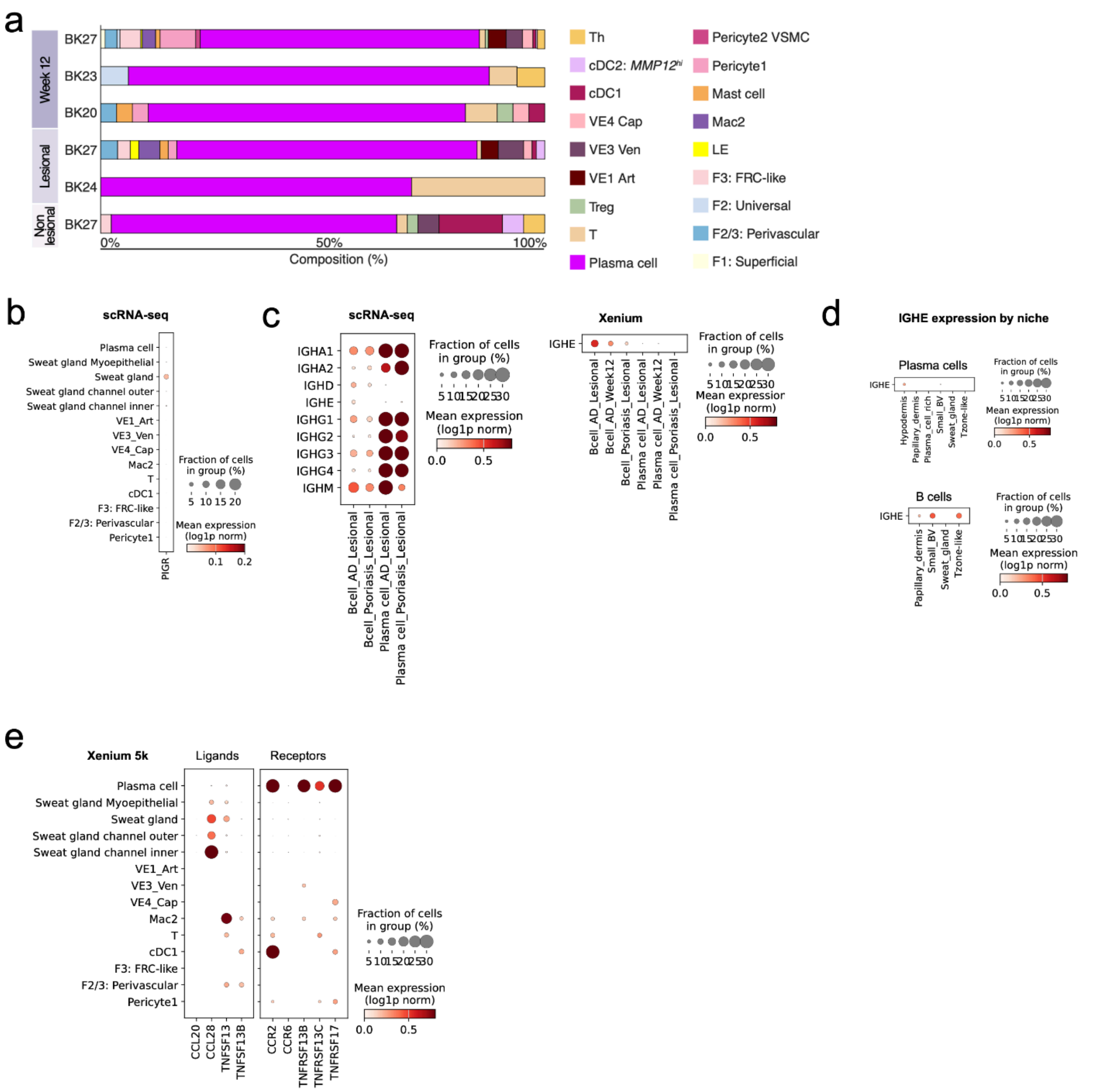


**Supplementary Fig. 18 | Plasma_cell_rich niche supporting genes.** **a,** Composition of *Plasma_cell_rich* niche by tissue section (minimum 10 cells). Cell types with <10 cell counts across all sections are excluded. **b,** Expression of *PIGR* in cell types found within the *Sweat_gland* niche, showing highest expression in sweat gland cells (scRNA-seq data). **c,**

Expression of immunoglobulin-related genes for B cells and plasma cells in scRNA-seq and Xenium data. The Xenium panel only included the *IGHE* gene. **d,** Expression of *IGHE* in plasma cells and B cells in Xenium data by niche (niches with minimum of 10 plasma/B cells) **e,** Expression of receptor/ligands reported for lung gland-associated immune niche in cell types within *Sweat_gland* niche for genes within Xenium 5k panel.

**Supplementary Fig. 19: Niche identification across different methodologies.**

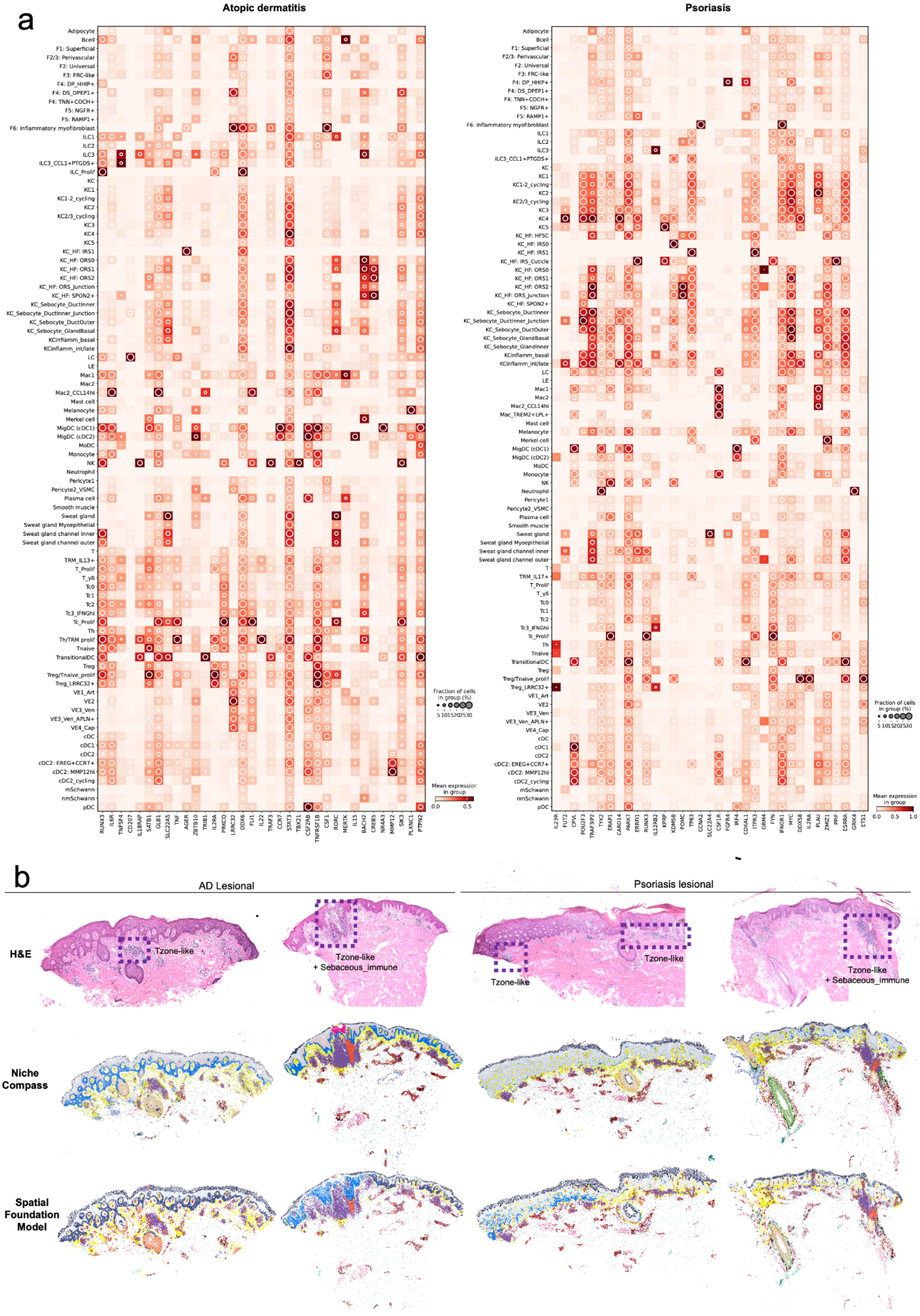


**Supplementary Fig. 19 | Niche identification across different methodologies. a,** Combined heatmap/dotplot of candidate genes for atopic dermatitis and psoriasis by cell type. **b,** Comparison of niches identified by NicheCompass and spatial foundation model (NEMO), showing consistent detection of the *Tzone-like* and *Sebaceous_immune* niches. These tissue sections were chosen as they previously shown in Fig. 3b and Extended Fig. 6b.

**Supplementary Fig. 20: Resolving high-resolution cell types in external datasets**

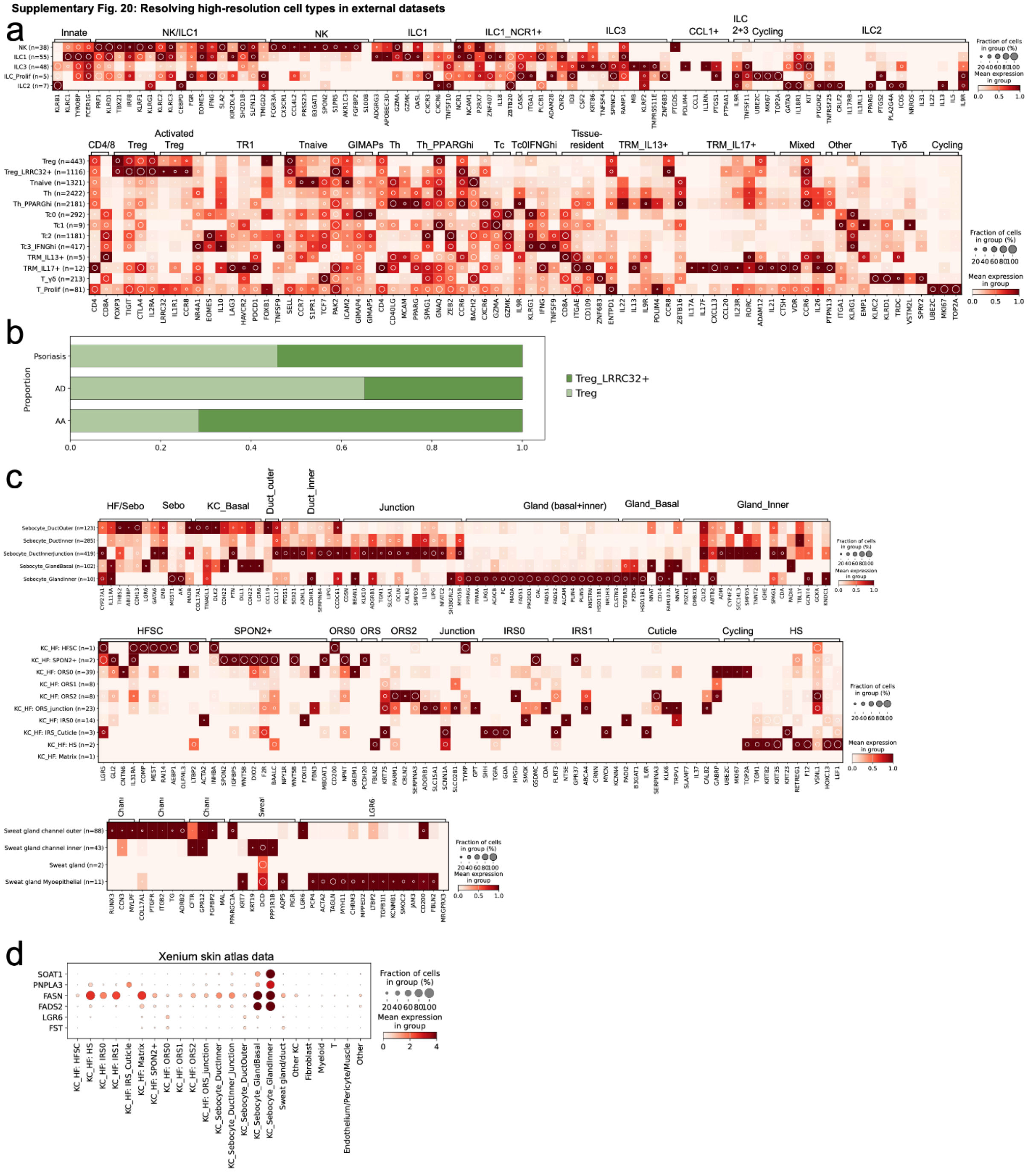


**Supplementary Fig. 20 | Resolving high-resolution cell types in external datasets. a,** Marker genes for T/ILC cells (as utilised for our original cell types) applied to alopecia areata dataset, showing consistent gene expression even  for rare cell types such as innate lymphoid cells. **b,** Proportion of Treg subtype by disease for psoriasis, AD, and AA. AA: Alopecia areata. AD: atopic dermatitis. **c,** Marker genes for keratinocyte appendage cells (as utilised for our original cell types) applied to acne dataset. **d,** Dot plot of candidate genes for acne genetic risk loci utilising improved annotations obtained for sebocyte cell populations in Xenium atlas data. ILC: innate lymphoid cell

**Supplementary Fig. 21: A queryable spatially-resolved resource of human skin at high-resolution.**

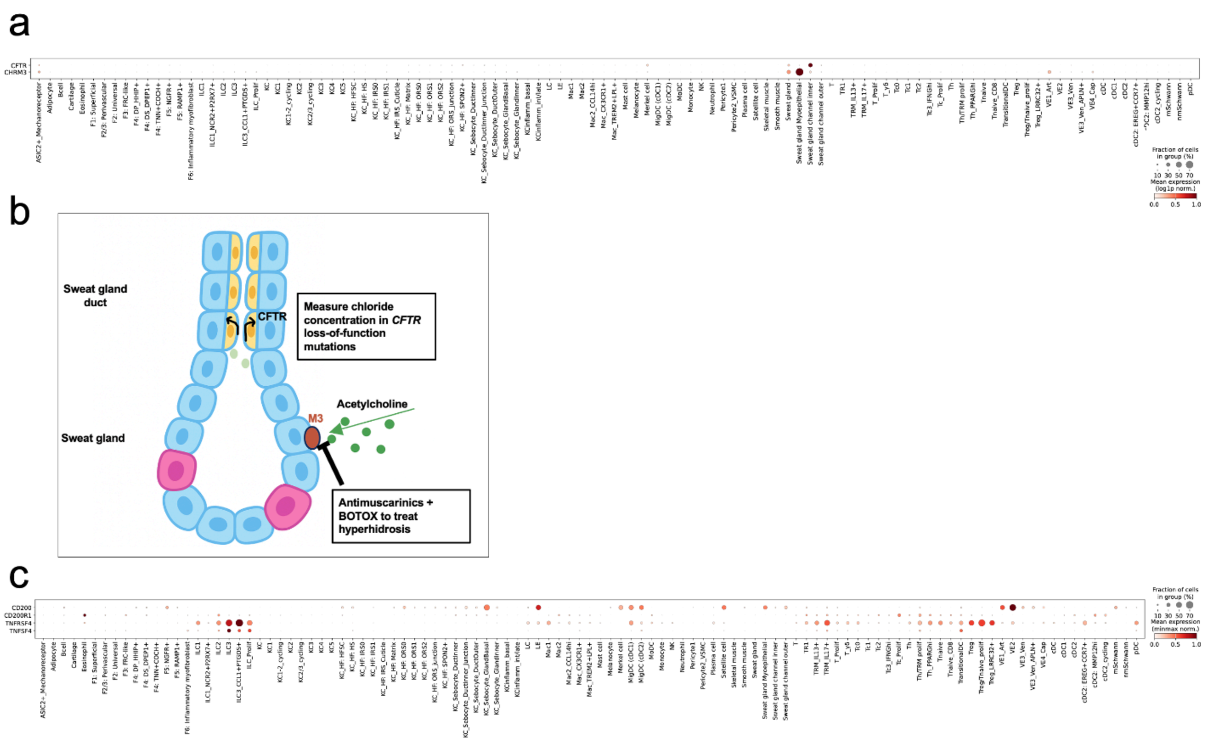


**Supplementary Fig. 21 | A queryable spatially-resolved resource of human skin at high-resolution. a,** Dot plot of *CFTR* and *CHRM3* expression in human skin. **b,** Schematic of utilising cell-specific level expression of *CFTR* and *CHRM3* to predict genetic investigations and treatment effects. **c,** Dot plot of expression of potential therapeutic targets in inflammatory skin disease in human skin (*TNFSF4*-*TNFRSF4*, *CD200*-*CD200R1*)).

**Supplementary Note 1**

Our atlas also serves as an interactive queryable resource for cell-specific queries, such as predicting safety of drug targets and/or planning diagnosis of genetic disorders. To illustrate this, we first describe two known examples for the sweat gland: diagnosis of *CFTR* mutations (cystic fibrosis) and treatment with M3 (antimuscarinic) inhibitors. CFTR regulates the movement of chloride ions from the lumen into the cell[142] and was uniquely expressed at high levels by *Sweat_gland_channel_inner* cells (Supplementary Fig. 21a). This finding suggests that abnormal chloride concentrations would be observed in cystic fibrosis, and consistently, the chloride sweat test is a first-line investigation for cystic fibrosis[142]. The M3 muscarinic receptor is encoded by *CHRM3,* which was expressed in the skin by both sweat gland and myoepithelial cells (Supplementary Fig. 21a). This finding suggests that reduced sweating would be a side-effect of drugs targeting this receptor, such as anticholinergic drugs. Consistently, impaired sweating (anhidrosis) is a recognised side-effect of anticholinergic medications and hyperhidrosis (excessive sweating) can be treated by anti-muscarinic treatments or other drugs that block the release of the M3 ligand (Supplementary Fig. 21b).

Novel drug targets are required for inflammatory skin diseases. We queried the expression of two potential target interactions[39], OX40-OX40L (encoded by *TNFRSF4* and *TNFSF4*, respectively) and CD200-CD200R The inhibitory immune checkpoint structure CD200R has been reported to be expressed on antigen-expressing cells and T cells[39]. We found expression on T cells (notably *ILC2s*) and antigen-presenting cells (particularly *cDC2: MMP12hi* cells), but notably, we also found high expression in eosinophils (Supplementary Fig. 21c). Expression of *TNFSF4* was observed on ILC subsets as well as *TransitionalDCs*.
